# Phylogenomics and ancestral reconstruction of Korarchaeota reveals genomic adaptation to habitat switching

**DOI:** 10.1101/2023.09.28.559970

**Authors:** Guillaume Tahon, Stephan Köstlbacher, Erik A. Pelve, Brett J. Baker, Jimmy H. Saw, Laura Eme, Daniel Tamarit, Max Emil Schön, Thijs J. G. Ettema

**Author notes:** **Correspondence:** Guillaume Tahon Thijs J. G. Ettema.

## Abstract

Our knowledge of archaeal diversity and evolution has expanded rapidly in the past decade. However, hardly any genomes of the phylum Korarchaeota have been obtained due to the difficulty in accessing their natural habitats and – possibly – their limited abundance. As a result, many aspects of Korarchaeota biology, physiology and evolution remain enigmatic. Here, we expand this phylum with five high-quality metagenome-assembled genomes. This improved taxon sampling combined with sophisticated phylogenomic analyses robustly places Korarchaeota at the base of TACK and Asgard clades, revisiting the phylum’s long-assumed position. Furthermore, we observe a clear split between terrestrial and marine thermal clades. Gene tree-aware ancestral reconstructions suggest that the last Korarchaeota common ancestor was a thermophilic autotroph. In contrast, Korarchaeaceae, the lineage where environmental transitions occurred, shifted towards a heterotrophic lifestyle. Terrestrial Korarchaeota gained many *cas* and CARF genes indicating they may need to manage viral infections. Together, our study provides new insights into these early diverging Archaea and suggests that gradual gene gain and loss shaped their adaptation to different thermal environments.

**Importance:** Korarchaeota are an ancient group of archaea, but their biology, physiology and evolution have remained obscure. Analysis of five novel Korarchaeota MAGs, and publicly available reference data provides robust phylogenomic evidence that Korarchaeota are placed at the base of Asgard archaea and TACK, revisiting the phylum’s long-assumed position. Gene content reconstruction suggests a versatile thermophilic and autotrophic last Korarchaeota common ancestor. Environmental distribution surveying of public databases places all Korarchaeota in thermophilic environments and indicates that their habitat is limited to hydrothermal vents and hot springs. Our modeling indicates at least two transitions linked to habitat switching between these environments in the evolutionary history of Korarchaeota. Both are linked to a significant alteration of the inferred ancestral gene content, including a shift towards a heterotrophic and potential scavenging lifestyle. Furthermore, hot spring Korarchaeota acquired various genes participating in resistance to viruses, suggesting they may need to manage frequent viral threats.

## Introduction

Archaea were only recognized as a third domain of life in 1977 (1), and were for a long time made up of only two phyla – Crenarchaeota and Euryarchaeota – represented by very few cultured strains (2). The third archaeal phylum to be discovered and the first one to be proposed without a cultured representative was *Candidatus* Korarchaeota (sometimes referred to as Korarchaeia (3) – hereafter referred to as Korarchaeota) (4). Amplicon sequences of the lineage were first sampled from Obsidian Pool, a near-neutral pH hot spring in the Hayden Valley of Yellowstone National Park (5). Subsequent 16S rRNA gene surveys and phylogenetic analyses supported the separation of this strain from the classical archaeal groups Euryarchaeota and Crenarchaeota, inspiring the name Korarchaeota, for the early divergence of the taxon in archaeal evolution (“koros/kore” referring to a young man/woman in Greek) (4). In 2008, ultrathin filamentous cells isolated from an anaerobic enrichment culture inoculated with Obsidian Pool sediment allowed initial characterization of a Korarchaeon - “*Candidatus* Korarchaeum cryptofilum” OPF8 (6). Its 1.59 Mb complete genome suggested an anaerobic heterotrophic lifestyle relying on a simple mode of peptide fermentation for generating biomass and energy. Also, the inability to synthesize various essential biomolecules indicated a potential dependency on other community members.

With the development of high-throughput DNA sequencing technologies and phylogenetic approaches in the four decades since the recognition of Archaea, the number and diversity of lineages described has increased exponentially. Currently, Archaea comprise four main clades (i.e., TACK, DPANN, Asgard and Euryarchaeota) which encompass over 30 phylum-level lineages and at least 50,000 species (See G. Tahon et al. (7) for a detailed overview). However, Korarchaeota have not undergone nearly the same expansion. Nearly 30 years after its initial discovery, very few Korarchaeota genomes have been sequenced, all of which have been recovered from either marine hydrothermal vents or terrestrial hot springs (Table S2) (8). Furthermore, all but the *Ca.* Korarchaeum cryptofilum OPF8 genome were published after 2018. As a result, many aspects of Korarchaeota biology, physiology and evolution have remained obscure. In addition, their phylogenetic position has been difficult to establish precisely. Initially, the two original environmental korarchaeotal 16S rRNA gene sequences branched deeply within Crenarchaeota, as a sister group to Eukarya or branching off before the common ancestor of Cren- and Euryarchaeota (4). Subsequent phylogenomic analyses based on concatenated sets of marker genes have not managed to unequivocally resolve the position of Korarchaeota. Depending on the phylogenetic method and dataset used, the phylum was placed either at the base of the TACK superphylum, as a sister group of Crenarchaeota inside TACK or as a sister lineage to Asgardarchaeota (9–13). However, the varying and frequently low-supported position of Korarchaeota within Archaea may be explained in part because of the use of less sophisticated models of evolution at the time, as well as by the lack of sequence representative of the korarchaeotal diversity. The latter led to the existence of a long branch at the base of Korarchaeota, which is well-known to yield phylogenetic reconstruction artefacts (14).

In this project, we expand the Korarchaeota genomic and taxonomic diversity with five distinct draft genomes obtained from sediment collected from the Little Hot Creek terrestrial hot spring (California, USA) and the Taketomi Island shallow submarine hydrothermal field of the Southern Ryukyu Archipelago (Japan). Using sophisticated phylogenomic analyses relying on multiple marker protein sets, we do not only accurately resolve the placement of Korarchaeota within the archaeal domain, but also the internal topology of the phylum. Furthermore, using gene tree-species tree reconciliation methods, we infer the evolution gene content along the korarchaeotal tree and describe how living in different marine and aquatic terrestrial thermal habitats shaped the genetic composition of the Korarchaeota.

## Materials and Methods

### Sampling

Samples were collected from the Taketomi Island shallow submarine hydrothermal field (TIV) of the Southern Ryukyu Archipelago (N24°20.9’ E124°06.10’) (15, 16) and from the Little Hot Creek terrestrial hot spring (N37°41.436’ W118°50.664’) (17, 18). Samples from Taketomi Island were collected as previously described by K. Zaremba-Niedzwiedzka et al. (19). Briefly, a sediment core was collected at 18 m depth, 5 m from the hydrothermal vent outflow by scuba diving, and material from the 10-15 cm section below the seafloor was extracted and stored at –25 °C and –80 °C. The core temperatures at 0, 20 and 30 cm below the sea floor were determined to be 31.0, 57.6 and 59.0 °C, respectively (RMT-2000; RIGO, Tokyo, Japan).

The Little Hot Creek (LHC) sample was collected as previously described (17, 18, 20). Briefly, cells from 80 °C sediment collected from the LHC4 hot spring in the Long Valley Caldera, California, USA, were separated by Nycodenz density centrifugation and stored either on ice (4 °C, in 10% v/v ethanol) or dry ice (80 °C, in 6% w/v betaine).

### DNA extraction and sequencing

DNA from the TIV sample was extracted using the Power Max Soil DNA isolation kit (MoBio Labs) and purified with the Aurora system (Boreal Genomics).

Metagenome libraries from the sample were prepared from 1 ng purified DNA using the Nextera XT DNA library kit and sequenced with the Illumina HiSeq 2000 instrument at the Uppsala SNP&SEQ Technology Platform, generating 49 Gb of 2×250 bp paired-end reads, as previously described by Zaremba-Niedzwiedzka et al. (2017).

Cells from the LHC samples were sorted within 1 week of collection and subsequently lysed. The DNA was amplified using multiple displacement amplification (MDA) in a microfluidic device and individual cells were screened with PCR, as previously described by J. A. Dodsworth et al. (20). Subsequently, 1 µl of the original single-amplified genome (SAG) DNA of the Korarchaeon was used in another round of MDA to generate additional DNA of this SAG for library preparation (18). The MDA reaction was performed using the Qiagen REPLI-g Mini kit and purified with the QIAamp DNA Mini kit (Qiagen, Hilden, Germany). Next, a sequencing library was generated from 1 ng of the resulting SAG DNA with the NexteraXT library preparation kit (Illumina, San Diego, CA, USA) and the library was then sequenced on an Illumina MiSeq instrument (2×300 bp), resulting in 4.9 Gbp of raw data.

Additionally, DNA was extracted from LHC sediment using the FastDNA Spin Kit for Soil (MP Biomedicals, Solon, Ohio, USA). Two metagenome libraries were generated using the Nextera DNA Sample Preparation kit (Illumina): a MiSeq and a HiSeq library which generated 7.3 and 10.8 Gbp, respectively.

### Assembly and binning

Illumina adapters in the raw reads of each metagenome dataset were removed using Scythe (https://github.com/vsbuffalo/scythe) with default parameters. Low quality regions in the sequences were trimmed with Sickle (https://github.com/najoshi/sickle) with the -*q 30* parameter to only keep reads with a PHRED score above 30. Afterwards, reads were assembled using SPAdes (version 3.5.0; https://github.com/ablab/spades) with the following settings *--meta -k 21, 33, 55*; and IBDA-UA (version 1.1.1; (21)) with the following settings: *--maxk 124*. Assembled contigs larger than 1 kbp were binned with the Emergent Self-Organizing Maps approach (ESOM) (22) with the following parameters: *-min 5000 -max 10000 -t robust*. korarchaeotal bins were visually inspected and manually selected using Databionic ESOM Tools (23). Bins containing SSU genes displaying high similarity to the *Ca.* K. cryptofilum SSU gene sequence were selected for further processing. A description of the *Ca*. Korarchaeota archaeon LHC assembly can be found in J. H. Saw et al. (18).

### Cleaning of new and reference Korarchaeota bins

The five korarchaeotal bins (one from LHC and four from TIV) were screened for non-korarchaeotal contigs using the R package multi-metagenome (https://github.com/MadsAlbertsen/multi-metagenome) (24). Contigs with deviant coverage, GC- or kmer content or with marker genes not phylogenetically clustered to other korarchaeotal genes were manually inspected and removed if deemed not likely to belong to the korarchaeotal clade.

Together with the five newly assembled Korarchaeota metagenome-assembled genomes (MAGs), a total of 34 publicly available MAGs classified as Korarchaeota in the NCBI (8), SILVA (25) and GTDB (3) databases were downloaded from NCBI in October 2020. The completeness and contamination levels of all MAGs were determined using CheckM v1.1.3 (26) and CheckM2 v1.0.1 (27). Additionally, MAGs were screened using an in-house RP15 pipeline, as previously described (28). Briefly, this pipeline was used to identify contigs that contain between 3 and 15 ribosomal proteins (RPs). The RPs found on these ‘*ribocontigs’* are then aligned to the orthologous RPs from a set of reference taxa. Subsequently, the separate alignments are trimmed, concatenated, and used to infer a phylogenetic tree using FastTree 2 (*-wag model*) (29) and IQ-TREE 2.1.2 (*LG+C60+F+G4 model*, *-bb 1000 -bcor 0.98*) (30).

Based on the position of ribocontigs in this tree, we observed that five of the publicly available MAGs clustered outside of Korarchaeota and we removed them from subsequent analyses.

### Phylogeny of 16S rRNA genes

To construct a phylogenetic maximum likelihood (ML) tree of the TACK superphylum, partial 16S rRNA gene sequences were extracted from Korarchaeota MAGs using barrnap v0.9 (https://github.com/tseemann/barrnap) with the flags *--kingdom arc/bac/euk*. Additionally, a total of 305 korarchaeotal sequences (Table S3) were retrieved from the SILVA Ref NR database SSU r138.1 (25), together with 2367 near full-length reference sequences (>1300 bp) representing the other taxa encompassed in the TACK superphylum. Asgardarchaeaota 16S rRNA gene sequences (>1300 bp; 305 sequences) were retrieved to be used as an outgroup. Several Korarchaeota 16S rRNA gene sequences retrieved from the MAGs were already present in the SILVA database and therefore removed from the dataset to avoid duplicate sequence entries.

The dataset was aligned using MAFFT v7.310 (*--auto*) (31). Subsequently, the alignment was processed using trimAl v1.4.rev22 to remove all columns with gaps in more than 80% of the sequences (*-gt 0.2*) (32). The resulting alignment was then used to construct a ML tree using IQ-TREE 2.1.2 (30), with settings: “*-m GTR+F -bb 1000 -altr 1000 --bcor 0.98*” (33).

The initial phylogenetic tree revealed that several publicly available Korarchaeota MAGs contained contigs harboring non-korarchaeotal 16S rRNA gene copies. These contigs were removed from the MAGs before downstream analyses. Additionally, two

Korarchaeota 16S rRNA gene sequences from the SILVA database were shown not to cluster with this phylum. These non-Korarchaeota 16S rRNA gene sequences were removed from the dataset after which the above steps were repeated.

If publicly available and documented, the sample origin (i.e., country and environment) from which the MAGs and 16S rRNA gene sequences originate was mapped onto the phylogenetic tree to visualize in which environments korarchaeotal clades have been detected. All trees were visualized using iTOL v6 (34).

### Environmental distribution assessment

To detect the presence of Korarchaeota in the environment, the 317 Korarchaeota 16S rRNA gene sequences were subjected to an integrated microbial NGS platform (IMNGS) search using a similarity threshold of 90% and a minimum size of 200 bp (35). From the results, sequences from datasets with less than 10.000 reads were deleted. The remaining Sequence Read Archive (SRA) sequence hits returned by the analysis were added to the aforementioned 16S rRNA gene alignment using the *-- add* command in MAFFT. This alignment was then used to construct a ML tree using the GTR+F model and ultrafast bootstrap support values as described above. Only SRA sequences clustering with Korarchaeota were retained from the IMNGS results. The origin of these SRA datasets was retrieved from NCBI and used to create an overview of the relative abundance of Korarchaeota in the environment (Table S4).

### 16S rRNA gene primer coverage of Korarchaeota

To analyze how well primers previously used to obtain Korarchaeota 16S rRNA gene sequences target different Korarchaeota clades, amplification primer information was, if documented, retrieved for the 294 PCR-amplicon-derived Korarchaeota 16S rRNA gene sequences in SILVA (Table S3). These primer sequences were then used in an *in silico* analysis which included all 317 Korarchaeota 16S rRNA gene sequences, together with 12 recently designed universal primers to determine primer coverage of different korarchaeotal lineages as described in G. Tahon et al. (7) (Table S5).

### Selection of Korarchaeota MAGs for species tree inference

To investigate the taxonomic diversity of Korarchaeota, average amino acid identity (AAI) of verified Korarchaeota MAGs was calculated using EzAAI (https://github.com/endixk/ezaai) (Table S6). Additionally, the 32 Korarchaeota MAGs were used as input for a phylogenetic analysis pipeline using GToTree (36) with settings “*-H Archaea -G 0.25”,* together with a previous in-house selection of 119 archaeal proteomes spanning archaeal diversity with an emphasis on Asgard archaeal representatives (Table S1). DPANN archaea were not included because they are only remotely related to TACK archaea and tend to be long-branching taxa in phylogenies, which can artefactually distort the inferred topology. The concatenated alignment obtained by GToTree was subsequently used to construct a ML phylogenetic tree with IQ-TREE v2.1.2 using 1000 ultrafast bootstraps and aLRT under the LG+C60+F+G4 model. Using this tree, the 16S rRNA gene tree, CheckM results, AAI values and the RP15 pipeline results, a subset of 18 MAGs - covering all major korarchaeotal lineages (Table S2) - was selected for amalgamated likelihood estimation (ALE) (37).

### Phylogenetic analyses of concatenated protein alignments for species tree inference

Two sets of phylogenetic markers were used to infer the species tree. The first one (RP56) is based on a previously published dataset of 56 RPs (19). The second one (NM54, for ‘new markers’) corresponds to 54 proteins extracted from a set of 200 markers previously identified as core archaeal proteins that can be used to robustly infer the tree of archaea (38, 39).

For both RP56 and NM54 marker sets, we gathered orthologs from all korarchaeotal predicted proteomes - predicted using prodigal (40) - as well as the other 119 archaeal proteomes. To retrieve orthologs from the set of selected taxa, we used sequences from the previously published alignments (19, 38) as queries for BLASTp against all proteomes. For each marker, the best BLAST hit from each proteome was added to the dataset. For the first iteration, each dataset was aligned using MAFFT-linsi (41) and ambiguously aligned positions were trimmed using BMGE 1.12 (*-m BLOSUM30*) (42). ML individual protein phylogenies were inferred using IQ-TREE v1.5.5 (43) under the LG+C20+F+G4 substitution model with 1000 ultrafast bootstraps and were manually inspected.

Individual marker phylogenies were inspected for erroneous inclusion of paralogous or contaminated sequences and to identify putative horizontal gene transfers. After deleting dubious sequences, datasets were realigned and trimmed, and phylogenies re-estimated as described above.

Trimmed alignments were then concatenated into two supermatrices (RP56 and NM54 supermatrices). For each dataset, a phylogeny was inferred using IQ-TREE under the LG+C60+F+G4 model. Statistical support for branches was calculated using 1000 replicates for ultrafast bootstrap and aLRT analyses.

The RP56 and NM54 supermatrix alignments were used to create four more alignments: an SR4-recoded alignment (44), an alignment where 50% of the fastest evolving sites were removed, a χ^2^-trimmed (28, 45) alignment where the 50% most heterogeneous sites were removed in steps of 1% and a custom recoded alignment. For the latter we calculated custom reduced amino acid schemes using the minmax chisq method (https://www.mathstat.dal.ca/~tsusko/doc/minmax-chisq.pdf) (44) and used these to recode sequence alignments.

For the 8 recoded or trimmed alignments, a phylogenetic ML tree was inferred using 1000 ultrafast bootstraps and aLRT under the GTR+G4+C60SR4 model (SR4-recoded alignments) or LG+C60+F+G4 model (other alignments). The ten trees (i.e., two supermatrices, five alignments each) were subsequently used to infer 100 non-parametric bootstrap trees using the posterior mean site frequency (PMSF) method under the transfer bootstrap expectation (TBE) interpretation (*-b 100 --tbe*) (46). Additionally, we ran four independent PhyloBayes-MPI v1.7a (47) Monte Carlo Markov Chains (*-cat -gtr*) for each of the alignments. Results were checked to determine whether convergence (maxdiff <0.3) and/or sufficient effective sample size (effsize[>[300) was reached. Subsequently, posterior predictive checks were performed with PhyloBayes-MPI to determine whether the inferred phylogenetic models adequately captured the across-taxa compositional heterogeneity and site-specific pattern diversity present in the alignments.

### Homology detection and protein family phylogenetic analysis

We used the annotation of arCOGs from EggNOG v5.0 (using eggnog-mapper v2.0.5) (48) to assign all 290,917 proteins from 137 taxa (both MAGs and reference genomes, Tables S1-2) into orthologous groups. All proteins without arCOG annotation were subjected to *de novo* clustering using OrthoFinder v2.4.0 *(-os -M msa*) (49), resulting in a total of 52,724 clusters (9,241 arCOG and 43,483 *de novo*). For all protein clusters with at least four members (i.e. 9,108) multiple sequence alignments were computed using MAFFT v7.475 E-INS-I (41) and subsequently processed with trimAl v1.4.1 to filter gap-rich alignment columns (*-gt 0.01*) (32).

Single-gene trees were inferred from these 9,108 alignments with IQ-TREE v2.0.3 with 1000 ultrafast bootstraps (*-bb 1000 -bnni -wbtl*) (30, 33). A model test was performed (*-m TESTNEW -mset LG -madd LG+C10,.., LG+C60*) for each tree inference (50). For clusters with only two or three members, only a single topology is possible for an unrooted tree and the unrooted gene trees were created for them.

### Gene tree-aware ancestral reconstruction

ALE v0.4 (37) was applied to reconcile single gene trees with the species trees based on ribosomal markers (RP56) and the new marker set (NM54). Firstly, conditional clade probabilities were computed from the bootstrap samples (*ALEobserve*) and 100 reconciliations with the species tree were sampled (*ALEml_undated*). We thereby adjusted the reconstructed genome copy number with the extinction probability per cluster within ALE (see github.com/maxemil/ALE/commit/136b78e).

We applied a threshold of 0.3 to the raw reconciliation frequencies that are the output of ALE, meaning that events (loss, transfer, origination or duplication) and presence/absence (copies) were counted as such if they had a frequency of at least 0.3 (51). Singleton clusters were counted as originations at the corresponding species node.

### Annotation of carbohydrate-active enzymes and transporters

Presence and distribution of glycosyl hydrolases, glycosyl transferases, polysaccharide lyases, carboxyl esterases, carbohydrate-binding modules and auxiliary activities were identified through the dbCAN2 meta server (52), using the CAZy database (July 2021). Carbohydrate-active enzymes predicted with at least two of the integrated tools were used. Transporters were identified using TransAAP (July 2021) (53).

### Korarchaeota optimal growth temperature

The optimal growth temperature of Korarchaeota MAGs was predicted using OGT_prediction (54).

Inference of growth temperature of the last Korarchaeota common ancestor was performed using the alignment and ML tree of the nucleoside diphosphate kinase sequences from the MAGs, obtained during the protein family phylogenetic analysis. Ancestral sequences and their unfolding midpoint temperatures (T_m_) were predicted using FireProt^ASR^ (55) and ProTstab2 (56), respectively.

### Reverse gyrase phylogeny

To create a phylogenetic tree of reverse gyrase, all corresponding protein sequences were retrieved from UniProt Release 2023_02 (57). Reverse gyrase sequences from the 137 MAGs, if not yet present in UniProt, were added to this dataset. Sequences were aligned with MAFFT v7.310 (*--auto*) after which the alignment was trimmed with trimAl v1.4.rev22 to remove all columns with gaps in more than 80% of the sequences (*-gt 0.2*). The ML phylogenetic tree was then generated using IQ-TREE v2.0.3 (*-bb 1000 -alrt 1000*) with the model LG+C60+F+R8, as chosen by the Bayesian information criterion (BIC) using ModelFinder Plus *(-m MFP -mset JTT, JTT+C10..C60, WAG, WAG+C10..C60, LG, LG+C10..C60, Q.pfam, Q.pfam+C10…+C60 -mrate G4, G6, G8, R4, R6, R8 -mfreq F*) (50).

### Relative evolutionary divergence

To determine a robust phylogenomic placement of a potential novel family-level lineage placed at the base of Korarchaeota by F. Vulcano et al. (58), seven MAGs from lineages Kg_2, Kg_3 and Kg_5, with a completeness estimation above 75% were added to our dataset containing the original 137 MAGs, and MAGs classified as *Ca*. Njordarchaeales. The single MAG from lineage Kg_4 could not be included due to its low completeness. The expanded dataset was subjected to the same analyses as before to create a NM54 supermatrix alignment. This alignment was χ^2^-trimmed as mentioned before, after which it was used to create a non-parametric bootstrap tree using the PMSF method under the TBE interpretation. The resulting ML tree was used as input for a phylogenomic analysis using the Castor package (59). Relative evolutionary divergence values were calculated for all nodes of the tree to determine whether this new lineage was part of the phylum Korarchaeota.

### Code availability

All scripts needed to reproduce the protein clustering and running ALE are available at https://github.com/maxemil/korarchaeota-scripts and https://github.com/maxemil/ALE-pipeline.

### Data availability

Sequencing raw reads and draft genome assemblies for the five new Korarchaeota MAGs have been deposited in the NCBI GenBank database under the accession numbers PRJNA983117 (TIV) and PRJNA983228 (LHC), respectively.

## Results and Discussion

### Contamination and misclassification among Korarchaeota

A first round of quality control using all contigs that contained (part of) the ribosomal protein (RP) operon, contigs that harbor partial 16S rRNA genes, and 76 single-copy archaeal marker proteins (see methods for detail) revealed that several of the publicly available metagenome-assembled genomes (MAGs) assigned as Korarchaeota were prone to contamination (Table S2). One MAG contained a 5 kbps contig harboring a bacterial 16S rRNA gene. Another MAG contained a contig harboring bacterial RPs. Finally, one MAG turned out to be a hybrid of bacterial and fungal contigs. In addition, some MAGs did not truly belong to Korarchaeota. Among these, three MAGs were found to represent *Ca*. Njordarchaeales, a recently described clade belonging to the Asgard archaea (39, 60), rather than Korarchaeota. A third MAG (DRAE01.1) was found to represent a yet undescribed lineage in TACK. Although sometimes classified as a lineage at the base of Korarchaeota, this placement was not supported and resulted from inferior phylogenomic analyses (3, 58). These findings underline that automated results generated by commonly used binning software, quality control tools and phylogenomic placement pipelines such as CheckM (26) and GTDB-tk (61) cannot be blindly trusted, because these tools do not perform phylogenetic validation of marker genes. For example, hybrid MAGs may have high completeness and low contamination values and even have robust phylogenetic placement. Therefore, thorough quality control and manual curation of MAGs is needed prior to their public deposit and classification. Phylogenetic analyses of the MAGs verified to be Korarchaeota, based on the 16S rRNA gene, contigs encoding at least 3 ribosomal proteins encoded on of a conserved ribosomal protein operon, and 76 single-copy archaeal marker proteins all revealed that Korarchaeota formed a monophyletic group in Archaea, however, with an unstable phylogenetic position and internal topology (Figs. S1-3). This urged us to perform in-depth phylogenomic analyses to resolve the phylogenetic position of Korarchaeota and its internal topology.

### Korarchaeota are placed at the base of the ‘ATACK superphylum’

The availability of 32 *bona fide* Korarchaeota MAGs in October 2020 allowed us to robustly place the phylum in the archaeal tree of life. To this end, we subsampled the dataset to reduce genomic redundancy of highly similar taxa in order to reduce computational burden. More Korarchaeota MAGs became publicly available after our initial selection (8, 58, 62). The vast majority of these comprised taxa already covered by our dataset. Some MAGs, however, represented a novel order at the base of Korarchaeota (58), as supported by estimation of relative evolutionary divergence values (Fig. S4). These MAGs became public when our project was in its final stage and could thus not be included in the analyses anymore. Further analysis of these MAGs indicated they did not change our findings.

Based on our initial phylogenomic tree including all MAGs (Fig. S3) and average amino acid identity (AAI) (Table S6) (63), we observed that the 32 MAGs could be grouped into 20 species, eight genera, four families and one order (Table S2, Fig. S3). The five MAGs obtained from in-house sequencing efforts all represented different species – four of which were novel – belonging to different genera. A total of 18 MAGs from all genera were then selected (with completeness >65%) for downstream analyses. For each taxon, we here propose *Candidatus* names that will be used throughout the text. Details on the taxonomic descriptions are given in the Supplementary Note. All names and the relevant nomenclatural types have been proposed under the SeqCode (64).

A first set of phylogenomic trees was inferred using 56 concatenated conserved ribosomal proteins (RP56) (39). In addition to the original concatenation, we performed four treatments aiming to reduce potential phylogenetic reconstruction artefacts yielded by saturation and compositionally biased sites (39, 65). For this, we first recoded our alignments into four character-states (SR4-recoding or custom-recoding) (44). We also removed the 50% fastest-evolving sites. Finally, we used a χ^2^ trimmer to remove the 50% most compositionally heterogeneous sites (28). The initial alignment and its four derivations (i.e., SR4-recoded, custom recoded, 50% fastest evolving sites removed, and χ^2^-trimmed) were then used to infer maximum likelihood (ML) trees using the posterior mean site frequency (PMSF) method (Figs. S5-9), and for phylogenetic reconstruction using PhyloBayes-MPI (Figs. S10-14).

Phylogenies based on all variations of the supermatrix, whether reconstructed in ML or Bayesian framework, showed Korarchaeota branching at the base of the TACK superphylum, although statistical support ranged from weak to high (80-99% bootstrap support; 0.69-1 posterior probability). The only exception was the Bayesian inference based on the RP56-SR4-recoded dataset, which placed Korarchaeota nested within TACK. Regarding the internal topology, we consistently recovered two distinct groups of MAGs originating from hot springs, representing the genera *Ca*. Korarchaeum and *Ca*. Methanodesulfokora, respectively. However, these were never found to be monophyletic, since the hydrothermal vent MAG representing the genus *Ca*. Fumariihydrokora always placed as sister-lineage to *Ca*. Korarchaeum. The rest of relationships between deep branches within the Korarchaeota varied depending on the analysis. In particular, we observed various placements of the hydrothermal vent genera *Ca*. Calidiprofundikora. and *Ca*. Sedimenticalidikora which are characterized by high genomic GC content (Fig. 2). This is in part reflected by the fact that, out of the five RP56 alignments, only the χ^2^-trimmed one was appropriately modelled by the sophisticated CAT-GTR model of evolution, based on bayesian posterior prediction tests (Table S7). And even in this case, the monophyly between the two clades from hot spring environments was poorly supported (Figs. S8 and S13).

**Fig. 1.**
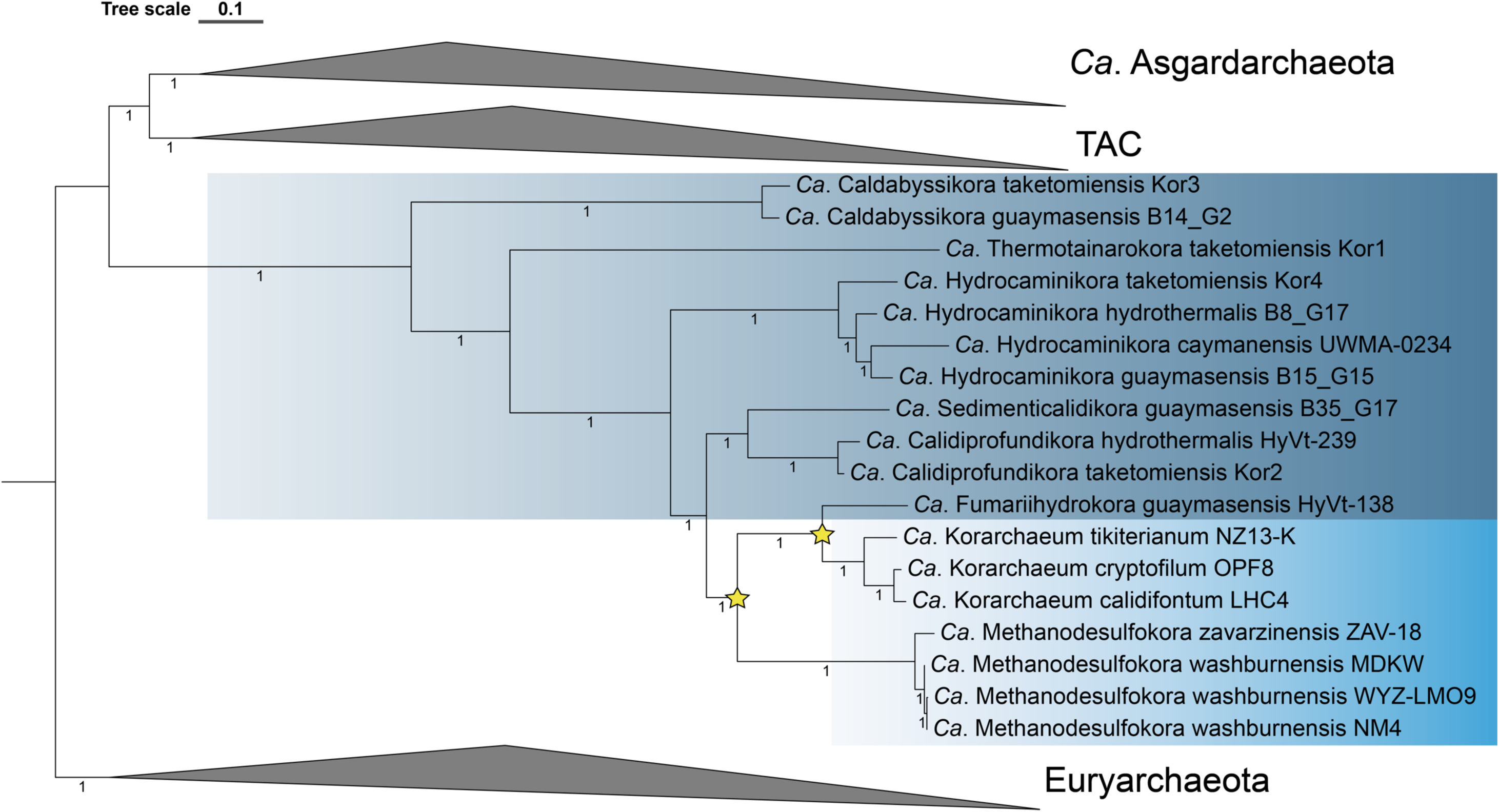
Phylogenetic tree corresponding to the Bayesian consensus tree (4 chains) reconstructed from the χ2-trimmed alignment (50% most heterogeneous sites removed; 7594 alignment positions) of 54 concatenated new marker proteins (NM54). *Ca*. Korarchaeota clades originating from hot springs and hydrothermal vents have a light and dark blue background shading, respectively. The two transitions between hydrothermal vents and hot springs are indicated with a star symbol. Scale bar indicates 0.1 substitutions per site.

**Fig. 2.**
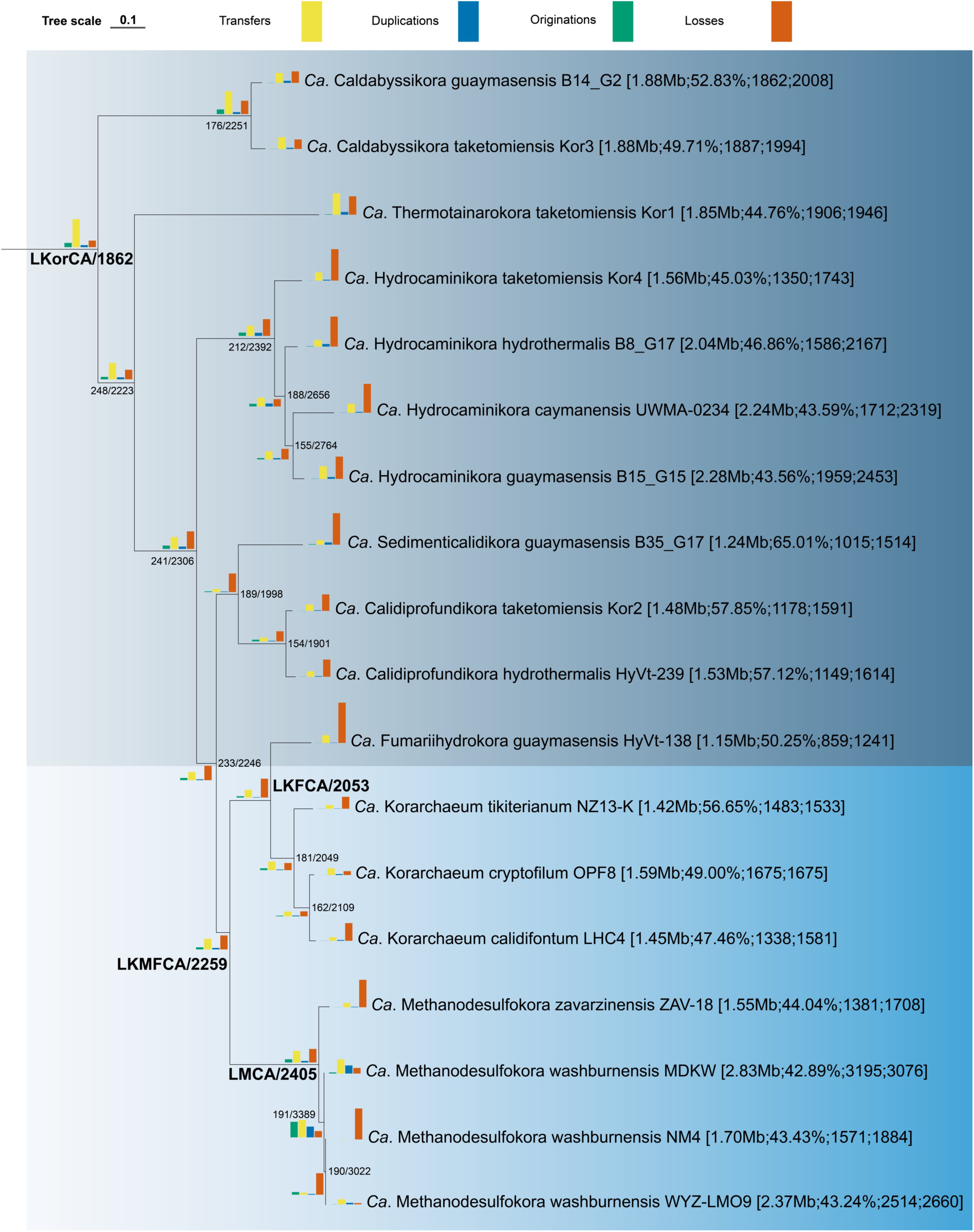
Overview of the Korarchaeota clade from Fig. 1. Internal nodes are annotated with the ancestor name/number code and the number of inferred coding sequences from the ALE analysis. Branches are annotated with bars representing the number of originations, transfers, duplications and losses. Bar heights in the legend correspond to 1000 such events each. Raw values can be found in Table S9. Terminal branches are labelled with the MAG name and accession number. Additional MAG statistics are given between square brackets: estimated genome size; GC content; number of CDS; estimated number of CDS.

As previously shown, RPs can yield phylogenomic reconstruction artefacts depending on the lineage considered (39). Despite them being highly conserved across the different domains of life, they are short in length (66) resulting in short concatenated alignments (i.e. 3575-7158 alignment positions in our case) and thus limited phylogenetic signal (Table S7). Furthermore, RPs are all part of a unique multisubunit complex, meaning they have undergone strong co-evolution. As a result, the assumptions of most evolutionary models of independence of evolution at sites are violated. Additionally, this implies that any departure from an average amino-acid composition will be propagated across subunits, and the noise/signal ratio amplified during phylogenetic reconstruction. This is particularly relevant since the amino-acid composition of RPs is strongly influenced by adaptation to high temperature environments, even more than the rest of the proteome (39). All Korarchaeota genomes were obtained from thermal environments, and using genomic information we estimated phylum members to have an optimal growth temperature of 70-85 °C (Table S2). One recently published Korarchaeota MAG was suggested to be mesophilic (62). However, the authors did not determine any optimal growth parameters of the organism. Here, we determined the optimal growth temperature of this MAG to be 79.3 °C, a value similar that of all other known Korarchaeota. Therefore, it can be assumed that this organism is a thermophile and not a mesophile. As a result of this adaptation to high temperature, the amino acid composition of their RPs has evolved strong thermostability-related signatures, such as a high ratio of charged versus polar amino acids, which can yield phylogenomic artefacts (39). An illustrative example of how problematic this can be is the recent case of the so-called *Ca*. Panguiarchaeales (67). Although described as a novel korarchaeotal lineage, phylogenomic and AAI analyses (Fig. S15, Table S8) show that all genomes described under this lineage name correspond to a low diverse taxon that instead belongs to the *Ca.* Njordarchaeales, a hyperthermophilic Asgard archaeal order (60). A similar problem arises in the work of J. Pan et al. (62), where several *Ca*. Njordarchaeales MAGs were considered as Korarchaeota (i.e., 3300022188_28 and GCA_011042815.1). *Ca*. Njordarchaeales have been shown to artefactually branch outside Asgard archaea as a sister lineage to Korarchaeota when inferring phylogenies based on RPs (39). The above examples show the importance of constructing robust phylogenomic trees including a broad diversity of reference data. Failure to do so (e.g., the work of Y. Liu and M. Li (68)) ultimately leads to incorrect results and conclusions.

To overcome limitations and inconsistencies specific to RP datasets, a second supermatrix alignment was constructed based on 54 core archaeal proteins that can be robustly used to infer the tree of Archaea (39). This new marker (NM54) supermatrix alignment and its four derivations were subjected to the same analyses as the RP56 alignments. Contrary to the RP-based phylogenies, all resulting NM54-based trees showed Korarchaeota not at the base of or within TACK, but instead at the base of a large superphylum composed of ‘TAC’ (i.e., TACK clade without Korarchaeota) and Asgard archaea, with high support (Figs. S16-24); altogether, they compose the informally named ATACK superphylum. Given this revisited position of Korarchaeota in the tree of Archaea using robust and sophisticated phylogenomic analyses, we show Korarchaeota to be a phylum-level lineage and not a class (i.e., *Ca*. Korarchaeia) of the Thermoproteota, as suggested by D. H. Parks et al. (3). Additionally, the internal Korarchaeota topology of the NM54 supermatrix alignments always showed a clear highly supported grouping of MAGs from hot springs, with the hydrothermal vent lineage *Ca*. Fumariihydrokora interrupting their monophyly. The χ^2^-trimmed alignment was again the only one appropriately modelled based on posterior prediction tests (Table 1). Contrary to its RP56 counterpart, the consensus tree of the four PhyloBayes-MPI Monte Carlo Markov Chains now showed maximum support for branches within Korarchaeota (Fig. 1). Additionally, the PhyloBayes sampling statistics showed sufficient effective sample size, and posterior prediction tests indicated that the model adequately captured the site-specific pattern diversity present in the NM54-χ^2^-trimmed alignment (Table 1). Therefore, we opted to use this alignment and the resulting phylogeny as species tree for the ancestral reconstruction analyses (below). We also performed phylogenomic analyses of our dataset (of the untreated and four treated alignments) expanded with *Ca*. Njordarchaeales (including the so-called *Ca*. Panguiarchaeales) MAGs. In the NM54 phylogenomic trees, contrary to the RP56 marker set, these lineages are not misplaced outside of the Asgard archaea and next to Korarchaeota but remain in place within the Asgard archaea (Fig. S25), thus showing that the NM54 marker set is better suited to resolve deep evolutionary relationships compared to marker sets based on ribosomal proteins. This highlights the importance of performing such sophisticated phylogenomic analyses prior to ALE. The different yet more robust and accurate placement of Korarchaeota in our study, compared to F. Vulcano et al. (58), J. Pan et al. (62) and Y. Liu and M. Li (68) undoubtedly impacts downstream analyses. For example, the previously estimated first divergence time of Korarchaeota (62) – when placed at the base of TACK – will greatly differ. With Korarchaeota placed at the base of ATACK, their estimated first divergence time would be in the Archean, long before the Great Oxidation Event. Therefore, Korarchaeota evolution was very unlikely driven by the availability of oxygen.

**Table 1.** Overview of Bayesian phylogenetic analyses results for the five New Marker (NM54) alignments.

| | Untreated | SR4 recoded | Custom recoded | 50% least heterogeneous sites ($\chi^2$-trimmed) | 50% fastest evolving sites trimmed |
| --- | --- | --- | --- | --- | --- |
| *General alignment statistics* |  |  |  |  |  |
| No. of sites | 15142 | 15142 | 15142 | 7594 | 7571 |
| Model of evolution | LG+C60+F+G4 | GTR+G4+C60SR4 | LG+C60+F+G4 | LG+C60+F+G4 | LG+C60+F+G4 |
| No. of informative sites | 13862 | 11810 | 13427 | 6341 | 6291 |
| Missing data (%) | 8.80 | 8.80 | 8.80 | 8.59 | 8.51 |
| *PhyloBayes sampling statistics* |  |  |  |  |  |
| Chains used for optimal Maxdiff | 4 | 2 | 2 | 4 | 2 |
| Cycles | 53599 - 54546 | 69504 - 69901 | 67722 - 68016 | 69911 - 70485 | 68356 - 70038 |
| Burnin | 10000 | 20000 | 20000 | 20000 | 20000 |
| Step | 30 | 30 | 30 | 30 | 30 |
| Effective sampling size | >287 | >15702 | >16772 | >3399 | >1510 |
| Optimal Maxdiff | 1 | 0.049 | 0.217 | 1 | 0.053 |
| Maxdiff 4 chains | 1 | 0.397 | 1 | 1 | 1 |
| *Posterior predictive tests* |  |  |  |  |  |
| Maximum squared heterogeneity (p-value) | 0 | 0 | 0 | 0.36 - 0.41 | 0 |
| Maximum squared heterogeneity (z-score) | 185.41 - 195.53 | 45.74 - 49.51 | 36.98 - 39.85 | 0.06 - 0.13 | 33.33 - 35.38 |
| Mean squared heterogeneity (p-value) | 0 | 0 | 0 | 1 | 0 |
| Mean squared heterogeneity (z-score) | 464.51 - 502.73 | 97.00 - 100.48 | 131.87 - 140.52 | -7.38 - -7.15 | 73.08 - 76.04 |
| Mean diversity per site (p-value) | 0 | 0.47 - 0.50 | 0.04 - 0.06 | 0 | 0 |
| Mean diversity per site (z-score) | 13.32 - 13.87 | 0.03 - 0.09 | 1.56 - 1.68 | 7.87 - 8.01 | 6.81 - 7.20 |

### Korarchaeota have a limited and environmentally confined distribution

Interestingly, all *bona fide* Korarchaeota MAGs originated from either hot springs or hydrothermal vents (Table S2), as was the case for all korarchaeotal 16S rRNA gene sequences from SILVA (Table S3). Because of the limited number of MAGs and 16S rRNA gene sequences, we also explored the phylogenetic diversity of the phylum using ∼280000 16S rRNA gene amplicon datasets included in the IMNGS platform (35). Only 161 datasets were found to contain Korarchaeota sequences (Table S4), as verified with a phylogenetic analysis (Fig. S2). Of these, only 5 and 24 had a Korarchaeota relative abundance of more than 1% and 0.1%, respectively. All but 4 datasets originated from known hydrothermal environments (Fig. S26). Together, these results suggest that Korarchaeota exhibit a high level of endemism and low abundance, even in their niche.

To investigate potential explanations for the restricted distribution and limited abundance of Korarchaeota, metadata was gathered for all MAGs, 16S rRNA gene sequences and IMNGS datasets. The majority of amplicon-based 16S rRNA gene sequences were obtained using Korarchaeota-specific and universal primers that were designed at least one decade ago (Table S5). An *in silico* primer coverage analysis showed that these primers almost exclusively target *Ca.* Korarchaeaceae with no or relatively few mismatches (Table S5). At the time of designing, only 16S rRNA gene sequences from these clades were available, which explains why these primers fail to target more distant Korarchaeota lineages. Various more recently developed universal primers, however, target the Archaea much better (Table S5) (7). Accordingly, the IMNGS datasets containing the highest Korarchaeota relative abundances were generated using these universal primers. Altogether, this suggests that the limited availability of Korarchaeota data is the result not only of their true low abundance and restricted distribution, but also of limited sequencing depth and the use of inaccurate primers.

All phylogenomic trees (Figs. S3-25) showed the presence of two distinct, yet low phylogenetically diverse clusters of MAGs originating from hot springs, representing the genera *Ca*. Korarchaeum and *Ca*. Methanodesulfokora. Each hot spring cluster was composed of a clade of multiple closely related MAGs from USA hot springs, with a non-USA (i.e., Russia and New Zealand, respectively) MAG at their base. Hydrothermal vent MAGs, on the other hand, form six distinct genera, their species originating either from the Taketomi Island shallow submarine hydrothermal field, Guaymas Basin or the Mid-Cayman Rise.

We wanted to check if the environmental and geographical split observed on Korarchaeota MAG phylogenies could be expanded to the larger amount of 16S rRNA gene sequence data available. The Korarchaeota topology of the 16S rRNA gene trees (Figs. S1-2) revealed that, compared to hot spring lineages, only few sequences grouped with the hydrothermal vent clades – especially the basal ones – which could be explained by primer bias and undersampling of these latter environments (see above). Overall, the two 16S rRNA gene trees were congruent with the phylogenomic ones. However, the relationships between hot spring and hydrothermal vent sequences were less well resolved. The hot spring lineages of *Ca.* Methanodesulfokora and *Ca.* Korarchaeum were comprised of closely related subclusters originating from Icelandic, Russian, USA and New-Zealand hot springs, and Malaysian, New Zealand, Japanese, Russian, Chinese, Bulgarian and USA locations, respectively (Table S3). Although similar in temperature and pH, these locations have vastly different chemistries. Therefore, despite their lower phylogenetic diversity, hot spring Korarchaeota could still be genetically distinct. To date, however, no MAGs are available from most of these sites. Interestingly, *Ca.* Methanodesulfokora contained one single hydrothermal vent sequence (accession number AF411237.1), whereas *Ca*. Fumariihydrokora, close relative of *Ca*. Korarchaeum, contained a single hot spring sequence (accession number DQ300332.1). These sequences originated respectively from a coastal shallow water hydrothermal vent and a geothermal well less than 100m from the beach, thus possibly representing environments with mixed characteristics. Nevertheless, biogeographic clustering suggests the existence of geographic variants.

All analyses support a Korarchaeota niche-invasion from marine to terrestrial geothermal habitats and a high degree of endemism. How this transition exactly took place and how genetically similar Korarchaeota from distant locations are remains unclear. Genetically similar korarchaeotal hot spring populations in the USA have been suggested to follow a spatial distribution resulting from a historical connection or an invasion-type event (69). Although these hypotheses could explain similar phylotypes in locations that are geographically close to each other, they fail to explain the clustering of phylotypes from distant locations. Instead, Korarchaeota probably adapted to specific environmental conditions of their respective habitats, and spread globally. For example, many hot springs where Korarchaeota have been detected share multiple environmental conditions (69, 70) that may have influenced the evolutionary origin of the two major hot spring phylotypes.

### The last Korarchaeota common ancestor: a versatile anaerobic autotrophic thermophile

To explore the genetic basis of the transitions between hydrothermal vents and hot springs or *vice versa*, and more generally how ancestral gene content evolved along the Korarchaeota species tree, we reconciled the histories of gene families and species tree using ALE (37). First, all proteins from the 137 MAGs were clustered into gene families, after which single-gene trees were inferred for all gene families.

Gene family histories were then inferred by reconciling gene trees with the species tree obtained from the NM54-χ^2^-trimmed alignment (Fig. 1) to infer four types of ancestral events (i.e., gene transfer, loss, duplication, and origination) as well as their resulting gene copy numbers on all ancestral nodes of the species tree (Fig. 2, Table S9). First, we screened the inferred gene content of the last Korarchaeota common ancestor (LKorCA).

The ALE analysis suggested that LKorCA produced biomass autotrophically by assimilating inorganic carbon via the Wood-Ljungdahl pathway, thus turning CO_2_ into acetyl-CoA (Fig. 3). Apart from the carbon monoxide dehydrogenase/acetyl-CoA synthase, LKorCA also encoded the genetic potential of pyruvate:ferredoxin oxidoreductase (*porABDG*). This key enzyme in anaerobic metabolism indicates that LKorCA could further reduce and carboxylate acetyl-CoA to pyruvate and *vice versa* (71).

**Fig. 3.**
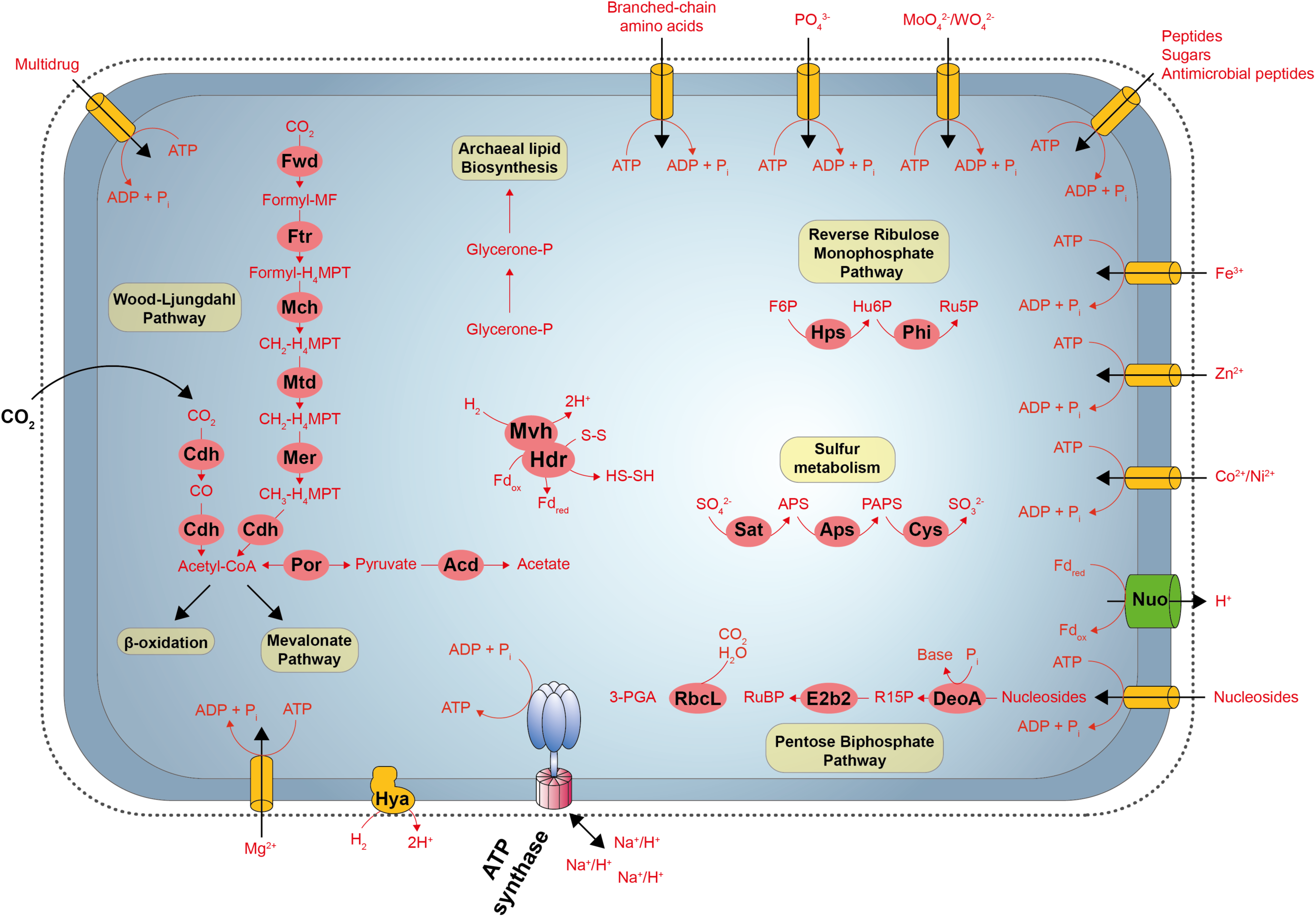
Ancestral gene content reconstruction of the Last Korarchaeota common ancestor (LKorCA). Genes were considered present if it was inferred by ALE with a copy number ≥ 0.3. For protein complexes, at least half the subunits had to be inferred as mentioned before.

LKorCA also contained a single *rbcL* copy, which encodes the archaeal type III ribulose-1, 5-bisphosphate carboxylase-oxygenase (RuBisCO) of the Calvin-Benson-Bassham (CBB) cycle. However, a complete CBB pathway was not present, which is common in archaeal genome sequences (72). Instead, the presence of thymidine phosphorylase (*deoA*) and eukaryotic translation initiation factor 2B (*e2b2*) hint that *rbcL* is involved in the utilization of ribonucleotides and ribonucleosides via the pentose-bisphosphate pathway (Fig. 3) (72, 73).

Inference of 3-hexulose-6-phosphate synthase (*hps*) and 6-phospho-3-hexuloisomerase (*phi*) in LKorCA suggests LKorCA could potentially rely on the reverse ribulose monophosphate pathway for biosynthesizing ribulose 5-phosphate in absence of the pentose phosphate pathway (74).

The inferred genetic potential of LKorCA also showed evidence for assimilatory sulfate reduction from sulfate to sulfite, a trait often found in conjunction with the Wood-Ljungdahl pathway (Fig. 3). LKorCA likely dwelled in deep-sea hydrothermal ecosystems. In these anoxic sediments niches carbon and sulfur cycling are intimately linked, and the diffusion of sulfate from seawater makes sulfate reduction a highly favorable process (75). Therefore, LKorCA might have played an important role in sulfur cycling. For nitrogen intake, LKorCA relied on a glutamine synthetase for the conversion of ammonium to glutamine. Phosphate, saccharides, metallic cations and peptides could be imported using ABC-type transporters. The presence of a complete branched-chain amino acid transport system (*liv* complex) suggests LKorCA could take up and utilize protein-degradation products from its environment. LKorCA also contained genes for the degradation of various amino acids, including alanine, serine and aspartate. The presence of genes encoding proteins for β-oxidation indicate at the genetic capacity of LKorCA to degrade fatty acids (Fig. 3).

LKorCA also encoded several cytoplasmic and membrane-bound complexes which are relevant for energy conservation. Presence of the respiratory complex NADH-ubiquinone oxidoreductase (*nuo*) would allow reoxidizing reduced ferrodoxin, which can in turn be used as electron donor, as supported by the absence of *nuoEFG*, the subunits required for the binding and oxidation of NADH (76). The Mvh/Hdr complex potentially coupled the endergonic reduction of ferredoxin by H_2_ to the exergonic reduction of a disulfide bond (77). Presence of a FAD-containing hydrogenase (*glcD*) could also indicate the use of HdrD/GlcD for heterodisulfide reduction (76). Hya and Mvh [NiFe]-type hydrogenases act in mediating the oxidation of H_2_, while an archaeal ATPase (Ntp) ensures energy conservation (Fig. 3). In addition, LKorCA showed genetic potential of HypA-F which are involved in the maturation of hydrogenases (78).

Although a complete *de novo* pyrimidine synthesis pathway was inferred in LKorCA, it lacked the purine counterpart. Therefore, LKorCA may have relied on transporting nucleosides across the membrane using the ABC uptake system encoded by *nupABC* and *bmpA* (Fig. 3).

Reverse gyrase, an enzyme which induces positive supercoiling in closed circular DNA in an ATP-dependent manner (79) and unique to (hyper)thermophilic organisms, was inferred as absent in LKorCA. However, all internal Korarchaeota ancestors and MAGs have a reverse gyrase. Phylogenetic analysis revealed that Korarchaeota reverse gyrase sequences are not monophyletic (Fig. S27). Sequences from *Ca*. Caldabyssikora were distinctly placed from those of the other Korarchaeota. Since Korarchaeota reverse gyrase sequences are not monophyletic, we cannot exclude that LKorCA already had it based on our inferences and that it might have been replaced in the last common ancestor of *Ca*. Caldabyssikora.

Although potential absence of reverse gyrase could indicate that LKorCA was not a hyperthermophile, this is very unlikely. Firstly, reverse gyrase is not necessarily a prerequisite for life at high temperatures, because the growth rate of a reverse gyrase knockout of *Pyrococcus furiosus* was only impacted above 90 °C (80). Therefore, the enzyme might not have been essential even if LKorCA had a thermophilic lifestyle. Secondly, the last common ancestors of each of the major archaeal clades were inferred to be thermophiles (81). Thirdly, nucleoside diphosphate kinase is a well conserved enzyme of which the stability is directly related to the host organism’s environmental temperature (82). The unfolding midpoint temperature of the reconstructed korarchaeotal ancestral nucleoside diphosphate kinase sequence (T_m_ = 78 °C) was more stable than that of the hyperthermophilic archaeon *Archaeoglobus fulgidus* DSM4304 (T_m_ = 62 °C) and thus indicated LKorCA was a thermophile.

LKorCA contains several genes encoding antioxidant enzymes (e.g., superoxide reductase, rubrerythrin). Previously, J. Pan et al. (62) suggested these genes to be absent from the *Ca*. Caldabyssikora (i.e., Kor 7) and *Ca*. Thermotainarokora (i.e., Kor 8). However, our extended Korarchaeota dataset including five newly constructed in-house MAGs, together with robust sophisticated phylogenomic analyses indicate that this is not the case.

### A first environmental transition is coupled to the switch to a likely heterotrophic lifestyle and higher virus predation

The NM54-χ^2^-trimmed alignment species tree indicated two environmental transitions, both appearing within the *Ca*. Korarchaeaceae family (Fig. 1). The first transition – from hydrothermal vents to hot springs – appeared at the branch before the diversification of the hot spring genera *Ca*. Korarchaeum, *Ca*. Methanodesulfokora and the hydrothermal vent genus Ca. Fumariihydrokora (‘last *Ca*. Korarchaeum, Methanodesulfokora and Fumariihydrokora common ancestor’ - LKMFCA). Upon this transition, the genetic repertoire of Korarchaeota was significantly altered (Fig. 4). Contrary to LKorCA, LKMFCA was not an autotroph. The Wood-Ljungdahl pathway, apart from *mch*, was lost in the last Korarchaeaceae ancestor and replaced by the glycolysis pathway. The latter, however, lacks the genes required for the reversible transformation between glucose and glucose-6-phosphate. LKMFCA’s lifestyle shifted towards that of fermentation, as could be observed by the gain of the *acs* gene allowing the reversible conversion of acetyl-CoA into acetate, and formate acetyltransferase.

**Fig. 4.**
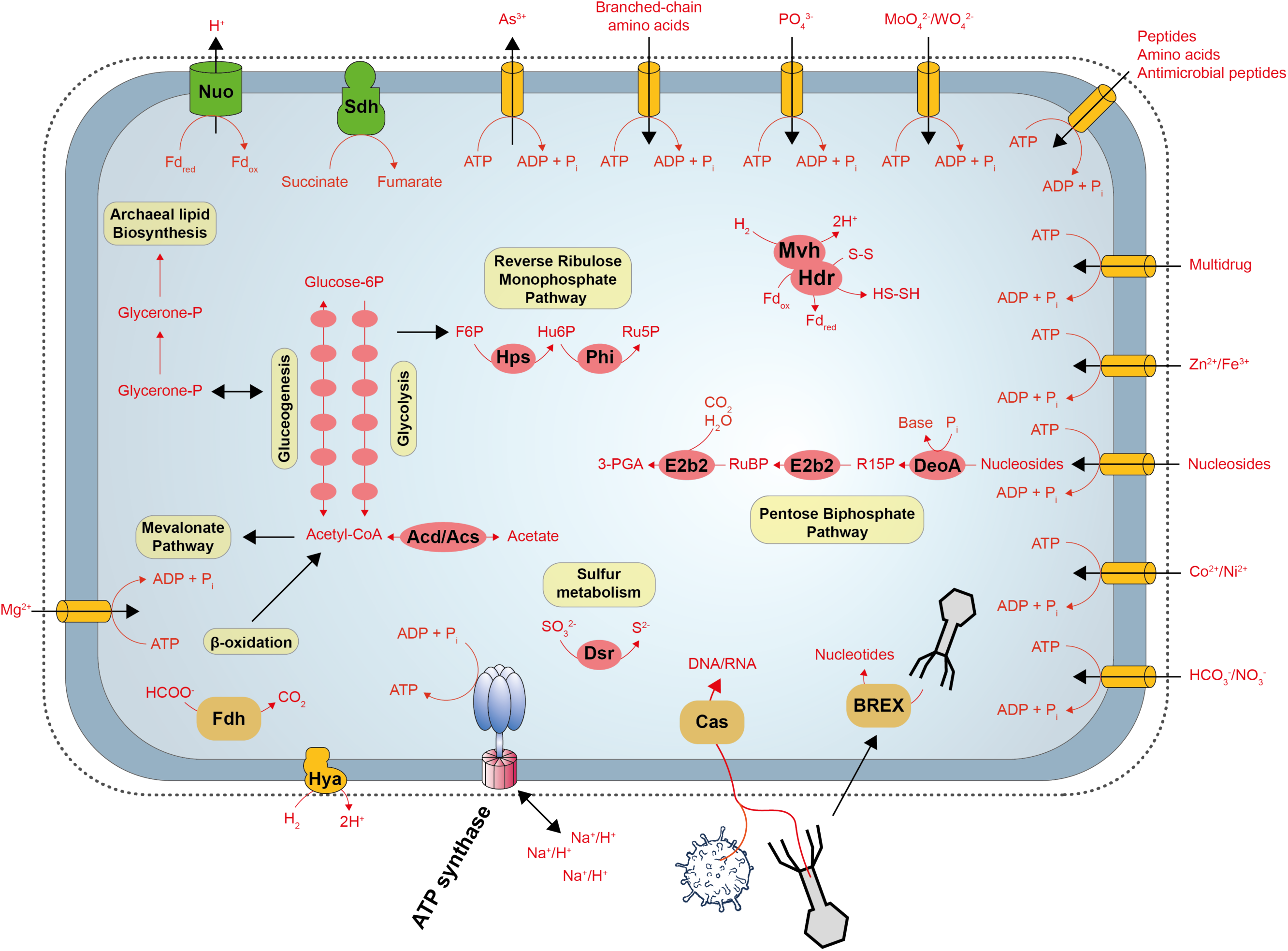
Ancestral gene content reconstruction of the Last *Ca*. Korarchaeum, *Ca*. Methanodesulfokora and *Ca*. Fumariihydrokora common ancestor (LKMFCA), which highlights the first transition from hydrothermal vents to hot springs. Genes were considered present if it was inferred by ALE with a copy number ≥ 0.3. For protein complexes, at least half the subunits had to be inferred as mentioned before.

Likewise, a shift took place in the sulfur metabolism, from assimilatory to dissimilatory (DsrAB) reduction of sulfur compounds (Fig. 4). This complex, together with the Mvh hydrogenase, indicates that hydrogen may have been used as an electron donor for the reduction of sulfite (83). The nitrogen metabolism repertoire was significantly expanded, including the glutamate dehydrogenase and glutamate synthase allowing the conversion of ammonia and glutamine, respectively, to glutamate. Additionally, presence of nitrate reductase suggests the ability to reduce nitrate to nitrite. A formate dehydrogenase allowed the conversion of formate to carbon dioxide. Compared to LKorCA, the genetic repertoire of LKMFCA also enlarged with various glycosyltransferases, mainly of families GT2 and GT4 which are commonly involved in processes such as N-glycosylation, cell wall biogenesis, cellulose, and sucrose biosynthesis (84). Where the electron transport chain of LKorCA consisted of only Complex I, that of LKMFCA was expanded with Complex II, the succinate dehydrogenase. The environmental switch also coincided with major changes in the amino acid metabolism of LKMFCA. Many of the biosynthesis pathways gained genes (e.g., LeuA-D), whereas genes required for degradation were lost. Pathways related to the metabolism of cofactors and vitamins were also subject to gene gain and now included a complete NAD^+^ biosynthesis pathway and greatly expanded thiamine metabolism. Furthermore, LKMFCA gained multiple transporter systems including those for arsenite efflux and transport of metal ions, amino acids, cations, tungstate and nitrate/bicarbonate. As a result, LKMFCA had an increased potential to scavenge its environment for nutrients and to efflux toxins out of its cell to survive (85). Interestingly, at the time of environmental transition, LKMFCA gained many genes participating in resistance to viruses, particularly those of type I and III *cas*, and CARF systems (86, 87). Given that both environments are hyperthermophilic, it is unclear exactly why these genes became enriched in hot spring Korarchaeota. One potential explanation could be a different virome, although further research is needed to answer this question. Several of the inferred type I and III genes (e.g. Cmr4g7, Cmr1g7) are known to be associated with hyperthermophiles, indicating they may be essential for managing viral infections in the hot spring habitats where many archaeal viruses reside (88, 89). Others, such as the BREX system, might allow phage adsorption while blocking its DNA replication (90). As such, the phage could be used as a source of nucleotides in support of growth of LKMFCA (Fig. 4).

### A reverse transition towards a scavenging or symbiont lifestyle

A second environmental transition, this time from hot spring to hydrothermal vent adaptation, happened before the diversification of *Ca*. Korarchaeum and Ca. Fumariihydrokora (LKFCA). While the central carbon metabolism of LKFCA did not change compared to LKMFCA, many changes, particularly gene losses, characterized this second transition (Fig. 5). LKFCA lost the ability to perform dissimilatory reduction of sulfur compounds and to reversibly convert acetyl-CoA into acetate. Likewise, the inferred genome of LKFCA is devoid of thymidine phosphorylase (*deoA*), the eukaryotic translation initiation factor 2B (*e2b2*) and the archaeal type III RuBisCO (*rbcL*) and thus lost the potential to use ribonucleotides and ribonucleosides via the pentose-bisphosphate pathway. The thiamine metabolism lost various key genes and was likely no longer functional. Given the absence of the *acs* and *mobAB* genes, the interconversion of acetate and acetyl-CoA, and synthesis of enzymes belonging to the DMSO reductase family were not possible anymore, respectively. Additionally, the repertoire of LKFCA no longer contained a tungstate ABC transporter. Despite many losses, LKFCA also gained various novel genes and thus pathways. Like in the transition to LKMFCA, many genes involved in amino acid biosynthesis were gained. Pathways for the biosynthesis of valine, leucine, isoleucine, lysine and phenylalanine expanded as well. A newly obtained spermidine/putrescine ABC transporter allowed regulating these polyamines which play vital roles in ion homeostasis and cell proliferation.

**Fig. 5.**
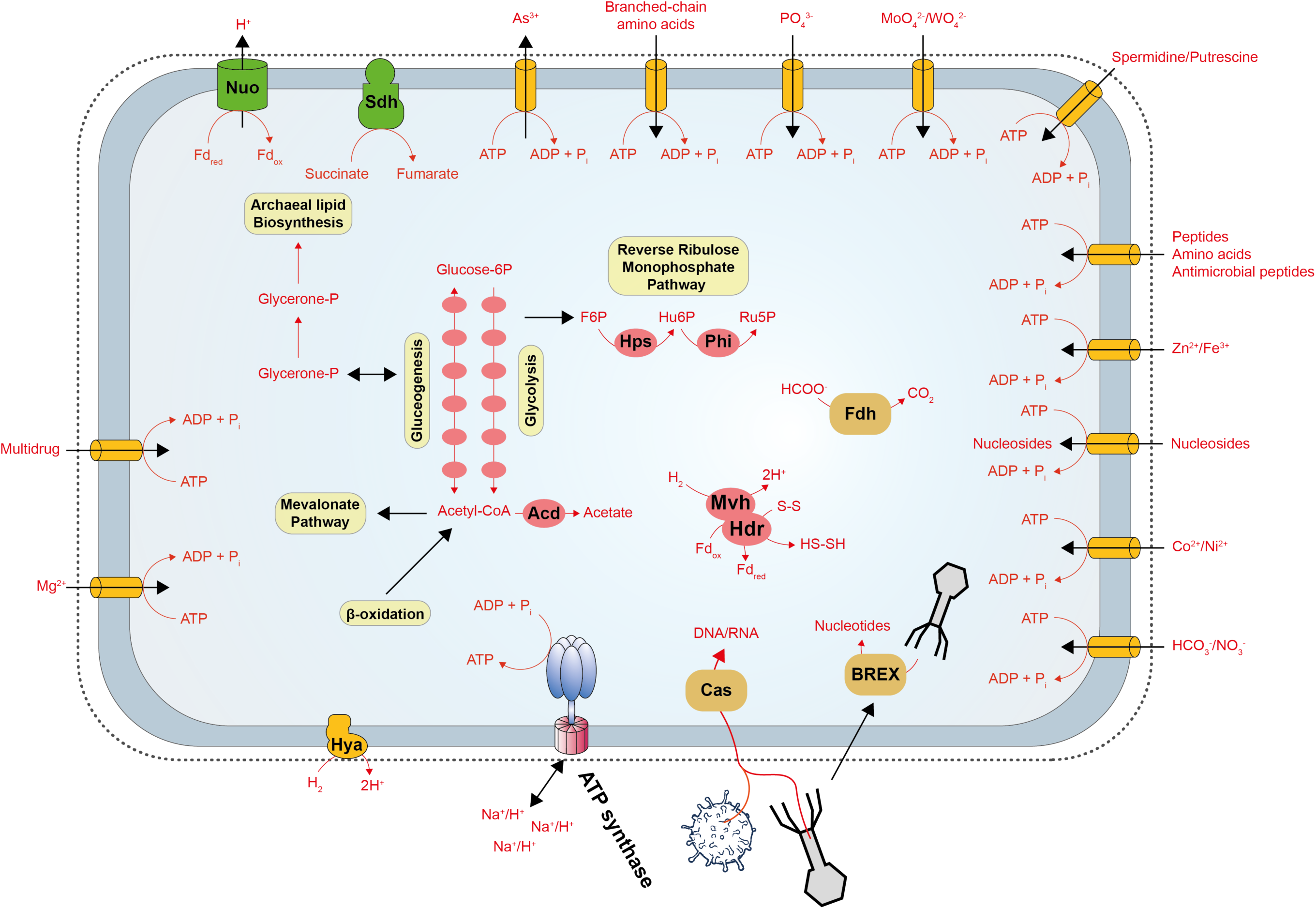
Ancestral gene content reconstruction of the Last *Ca*. Korarchaeum and *Ca*. Fumariihydrokora common ancestor (LKFCA), which highlights the second transition from hydrothermal vents to hot springs. Genes were considered present if it was inferred by ALE with a copy number ≥ 0.3. For protein complexes, at least half the subunits had to be inferred as mentioned before.

Taken together, the second transition indicated that LKFCA underwent even more metabolic changes towards a simplified genomic machinery. Contrary to the once rather metabolically complex LKorCA, LKFCA transitioned to a simple anaerobic heterotrophic lifestyle of peptides and amino acids (6). Whether LKFCA is just a very efficient scavenger or a symbiont, or perhaps both, awaits to be proven with the cultivation of Korarchaeum strains.

### The last Methanodesulfokora common ancestor: a virus-combatting methanogen

The last *Ca*. Methanodesulfokora common ancestor (LMCA) followed a different evolutionary trajectory. Where LKFCA lost the *rbcL*, *e2be*, *deoA* and *dsrAB* genes, and several key genes of the thiamine metabolism, LMCA kept this genomic potential. LMCA also had an expanded amino acid metabolism, yet less extensive than LKFCA. Several more type I and type III *cas* system genes participating in resistance to viruses were acquired, indicating that *Ca*. Methanodesulfokora spp. may need to manage additional viral threats in the hot springs where they reside (Fig. 6). Interestingly, we inferred in LMCA a whole suite of genes encoding methanol methyltransferase (MtaABC) and methyl-coenzyme M reductase (McrABG) for methanol reduction and methane production. Although *dsr* and *mcr* genes could hint at a sulfite-dependent, anaerobic oxidation of methane to methanol (83), recent cultivation of a strain of a *Ca*. Methanodesulfokora washburnensis has validated its methylotrophic methanogenic potential (Personal communication with Roland Hatzenpichler). Using genomic data, only *Ca*. M. washburnensis MAGs – all obtained from USA hot springs – appear to have the Dsr, Mcr and Mta complexes. Whether they are truly absent in the other *Ca*. Methanodesulfokora species, currently comprises a single MAG obtained from a Russian hot spring, needs to be further investigated by recovery of more complete MAGs and cultivated strains.

**Fig. 6.**
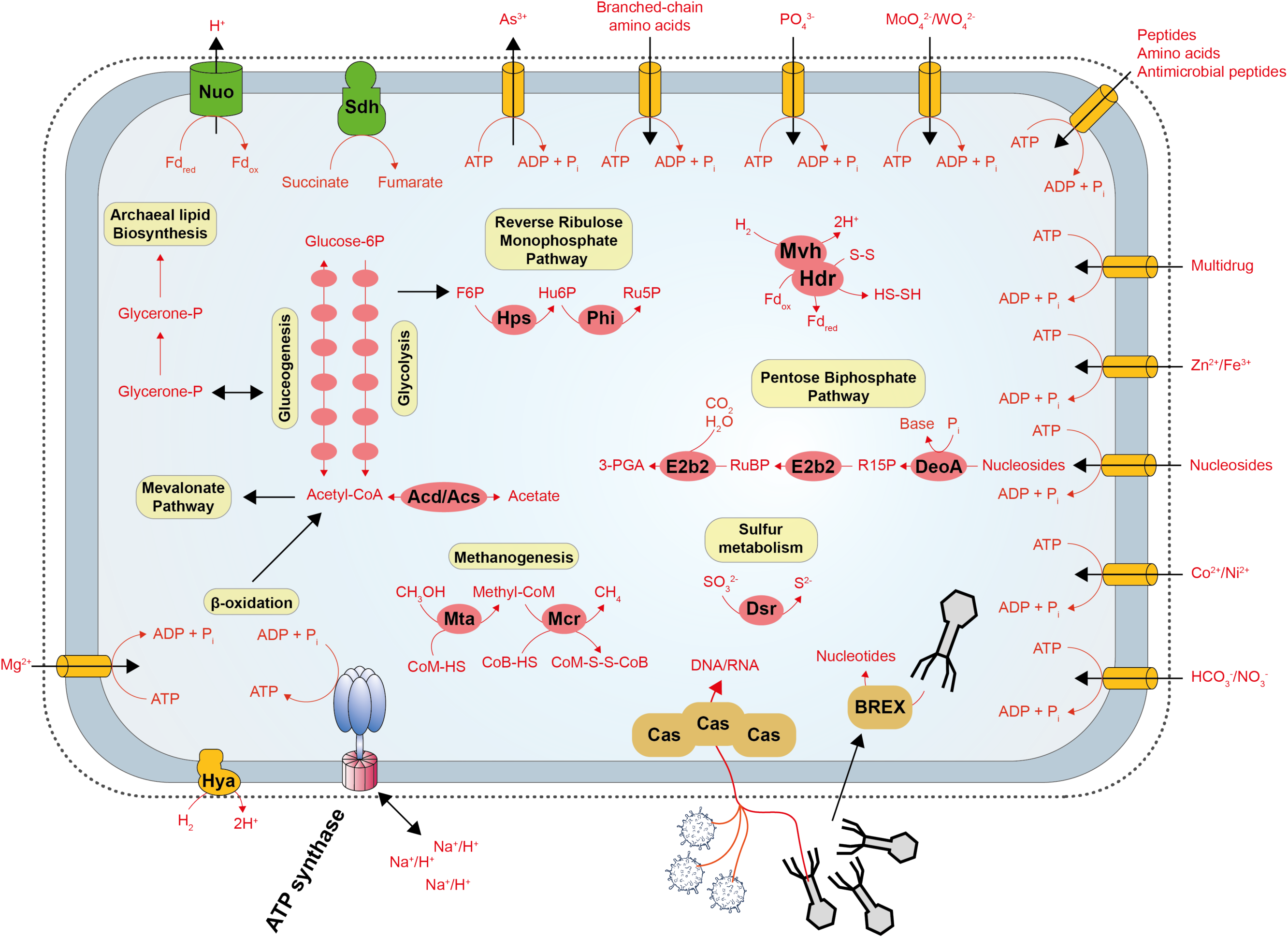
Ancestral gene content reconstruction of the Last *Ca*. Methanodesulfokora common ancestor (LMCA). Genes were considered present if it was inferred by ALE with a copy number ≥ 0.3. For protein complexes, at least half the subunits had to be inferred as mentioned before.

## Conclusion

Despite rapid advancements in metagenomic sequencing analysis which particularly expanded our view of the archaeal branch of the Tree of Life, our understanding of Korarchaeota genomics is limited. Here, the addition of five new MAGs improved the phylogenomic confidence, placing the phylum at the base of TACK and Asgard, creating an ATACK superphylum, and showed a clear korarchaeotal endemism and two habitat transition between marine and terrestrial geothermal habitats. The inclusion of MAGs from all currently known korarchaeotal genera in gene tree-aware ancestral genome reconstructions revealed that these environmental transitions were subject to significant metabolic changes, thereby completely reshaping their genetic composition. Our work provides new insights into the distribution, evolution and physiology of the Korarchaeota and guides future cultivation-dependent and - independent studies to further unravel its enigmatic biology and lifestyle.

## Supporting information

Etymology of (new) Korarchaeota taxa

## Acknowledgements

The authors would like to express their appreciation to Nina Dombrowski and Kassiani Panagiotou for their support. We thank Peter Vandamme for advice on the etymology of the new names proposed.

## Funding

This work was supported by grants from the European Research Council (ERC) (Consolidator grant 817834 to T.E. and Starting grant 803151 to L.E.), the Swedish Research Council (International Postdoc 2018-06609 to D.T.) and the Volkswagen Foundation (grant no. 96725 to T.E.). This work made use of the Dutch national e-infrastructure with the support of the SURF Cooperative using grant no. EINF-2953.

## Conflict of interest

The authors declare there is no conflict of interest.

## Notes

### Competing Interest Statement

The authors have declared no competing interest.

### Summary of Updates

Added Supplementary Material

