## Supplementary Table Captions for "Phylogenomics and ancestral reconstruction of Korarchaeota reveals genomic adaptation to habitat switching"

*** Correspondence:**

Guillaume Tahon

Archaea, microbial diversity, phylogeny, hyperthermophiles, hot spring, genome evolution

### Supplementary Table Captions

**Table S1. Metadata of the 119 non-Korarchaeota reference MAGs included in the analyses.**

**Table S2. Metadata for all *Candidatus* Korarchaeota MAGs used in this study.** Taxonomic descriptions of new taxa are given in the Supplementary Note.

**Table S3. Metadata of *Candidatus* Korarchaeota 16S rRNA gene sequences obtained from SILVA 138.1.** Taxonomic descriptions of new taxa are given in the Supplementary Note.

**Table S4. Global *Candidatus* Korarchaeota distribution based on 16S rRNA gene sequences retrieved using IMNGS.**

**Table S5. Coverage of universal and *Ca*. Korarchaeota specific primers with 16S rRNA gene sequences from *Ca*. Korarchaeota MAGs.** The first tab (Table S5a) shows the primer mismatches and coverage of universal and *Ca*. Korarchaeota-specific primers with partial 16S rRNA gene sequences retrieved from *Ca*. Korarchaeota MAGs. The second tab (Table S5b) lists *Ca*. Korarchaeota-specific primer sequences retrieved from literature with the publications they were retrieved from. Taxonomic descriptions of new taxa are given in the Supplementary Note.

**Table S6. Pairwise average amino acid identity (AAI) values between 32 *Ca*. Korarchaeota MAGs used in this study.**

**Table S7. Overview of Bayesian phylogenetic analyses results for the five Ribosomal Protein (RP56) alignments.**

**Table S8. Pairwise average amino acid identity (AAI) values between *Ca*. Panguiarchaeales and *Ca*. Njordarchaeales MAGs.**

**Table S9. Raw ALE values for the number of originations, transfers, duplications and losses for *Candidatus* Korarchaeota.**
