## Supplementary material for "Phylogenomics and ancestral reconstruction of Korarchaeota reveals genomic adaptation to habitat switching": Table S7

**Table S7. Overview of Bayesian phylogenetic analyses results for the five Ribosomal Protein (RP56) alignments.**

| | **Untreated** | **SR4 recoded** | **Custom recoded** | **50% least heterogeneous sites ($\chi^2$-trimmed)** | **50% fastest evolving sites trimmed** |
|---|---|---|---|---|---|
| *General alignment statistics* | | | | | |
| No. of sites | 7158 | 7158 | 7158 | 3602 | 3575 |
| Model of evolution | LG+C60+F+G4 | GTR+G4+C60SR4 | LG+C60+F+G4 | LG+C60+F+G4 | LG+C60+F+G4 |
| No. of informative sites | 6672 | 5730 | 6050 | 3145 | 3089 |
| Missing data (%) | 7.21 | 7.21 | 7.21 | 7.09 | 6.86 |
| *PhyloBayes sampling statistics* | | | | | |
| Chains used for optimal Maxdiff | 2 | 4 | 3 | 3 | 4 |
| Cycles | 14434 - 14669 | 68598 - 69795 | 57658 - 58701 | 49750 - 50669 | 31757 - 31914 |
| Burnin | 5000 | 20000 | 20000 | 20000 | 20000 |
| Step | 10 | 30 | 30 | 30 | 10 |
| Effective sampling size | >114 | >26927 | >13376 | >421 | >2013 |
| Optimal Maxdiff | 0.226 | 0.102 | 0.087 | 0.096 | 0.25 |
| Maxdiff 4 chains | 1 | 0.102 | 1 | 0.641 | 0.24 |
| *Posterior predictive tests* | | | | | |
| Maximum squared heterogeneity (p-value) | 0 | 0 | 0 | 0.39 - 0.42 | 0 |
| Maximum squared heterogeneity (z-score) | 73.99 - 74.42 | 32.80 -34.20 | 16.31 - 16.61 | 0.02 - 0.08 | 9.55 - 10.52 |
| Mean squared heterogeneity (p-value) | 0 | 0 | 0 | 1 | 0 |
| Mean squared heterogeneity (z-score) | 216.45 - 219.23 | 81.77 - 84.15 | 40.04 - 42.01 | -8.55 - -8.26 | 33.99 - 36.13 |
| Mean diversity per site (p-value) | 0 | 0.56 - 0.57 | 0.30 - 0.33 | 0 | 0 |
| Mean diversity per site (z-score) | 9.33 - 9.81 | -0.15 - -0.12 | 0.44 -0.49 | 5.44 - 5.63 | 5.29 - 5.40 |
