## Supplementary material for "Phylogenomics and ancestral reconstruction of Korarchaeota reveals genomic adaptation to habitat switching": Etymology of (new) Korarchaeota taxa

*** Correspondence:**

Guillaume Tahon

Archaea, microbial diversity, phylogeny, hyperthermophiles, hot spring, genome evolution

**Etymology of (new) *Candidatus* Korarchaeota taxa**

**Emended description of *Candidatus* Korarchaeum Elkins et al. 2008**

Kor.ar.chae’um. Gr. masc. n. *koros*, young man; N.L. neut. n. *archaeum*, archaeon; N.L. neut. n. *Korarchaeum*, young archaeon because of the early divergence of the group. Estimated genome size of *Ca*. Korarchaeum MAGs range from 1.18 to 1.74 Mb, their genomic G+C content from 47.46 to 56.65 %. The representative MAG of the genus is that of *Ca*. Korarchaeum cryptofilum OPF8. Its NCBI accession number is GCF_000019605.1.

**Emended description of *Candidatus* Korarchaeum cryptofilum Elkins et al. 2008**

cryp.to.fi’lum. Gr. masc. adj. *kryptos*, hidden; L. neut. n. *filum*, a thread; N.L. neut. nom. n. *cryptofilum*, a hidden thread. The representative MAG of the species is that of *Ca*. Korarchaeum cryptofilum OPF8. Its NCBI accession number is GCF_000019605.1.

DNA G+C content of the draft genomes is 48.68 – 49.15%, approximate genome sizes are 1.59 - 1.74 Mb.

**Description of *Candidatus* Korarchaeum calidifontum sp. nov.**

ca.li.di.fon’tum. L. masc. adj. *calidus*, hot; L. masc. n. *fons*, spring, fountain; N.L. gen. neut. n. *calidifontum*, originating from a hot spring.

The representative MAG of the species is that of *Ca*. Korarchaeum calidifontum LHC4 (JAVCBY000000000). Its estimated genome size and DNA G+C content are 1.45 Mb and 47.46%, respectively. A second MAG (*Ca*. Korarchaeum calidifontum SpSt-718) has an estimated genome size and DNA G+C content are 1.51 Mb and 47.68%, respectively.

**Description of *Candidatus* Korarchaeum tikiterianum sp. nov.**

ti.ki.te.ri.a’num. -*anum* adjective-forming suffix; N.L. neut. adj. *tikiterianum*, of or belonging to the Tikiteri hot spring in New Zealand.

The representative draft genome is that of *Ca*. Korarchaeum tikiterianum NZ13-K (NCBI accession number MAIU00000000). As derived from the draft genome the estimated genome size is 1.42 Mb, the DNA G+C content 56.65%.

**Description of *Candidatus* Korarchaeum nevadense sp. nov.**

ne.va.den’se. N.L. neut. adj. *nevadense*, pertaining to Nevada, where the first MAGs originate from.

The representative draft genome is that of *Ca*. Korarchaeum nevadense SpSt-15 (NCBI accession number DSGI00000000). As derived from the genome the estimated genome size is 1.60 Mb, the DNA G+C content 47.85%.

**Description of *Candidatus* Korarchaeaceae fam. nov.**

*Korarchaeaceae* (Kor.ar.chae.a.ce’ae, N.L. neut. n. *Korarchaeum*, the type genus of the family; suff. -*aceae*, ending to denote a family; N.L. fem. pl. n. *Korarchaeaceae*, the family of the genus *Korarchaeum*).

The description is the same as for the genus *Korarchaeum*. Type genus is *Korarchaeum*.

**Description of *Candidatus* Korarchaeales ord. nov.**

*Korarchaeales* (Kor.ar.chae.a’les, N.L. neut. n. *Korarchaeum*, the type genus of the order; suff. -*ales*, ending to denote an order; N.L. fem. pl. n. *Korarchaeales*, the order of the genus *Korarchaeum*).

The description is the same as for the genus *Korarchaeum*. Type genus is *Korarchaeum*.

**Description of *Candidatus* Korarchaeia class. nov.**

*Korarchaeia* (Kor.ar.chae’i.a, N.L. neut. n. *Korarchaeum*, the type genus of the order of the class; suff. -*ia*, ending to denote a class; N.L. neut. pl. n. *Korarchaeia*, the class of the order *Korarchaeales*).

The description is the same as for the genus *Korarchaeum*. Type genus is *Korarchaeum*.

**Description of Candidatus Korarchaeota phyl. nov.**

*Korarchaeota* (Kor.ar.chae.o’ta, N.L. neut. n. *Korarchaeum*, the type genus of the class of the phylum; suff. -*ota*, ending to denote a phylum; N.L. neut. pl. n. *Korarchaeota*, the phylum of the class *Korarchaeia*).

The description is the same as for the genus *Korarchaeum*. Type genus is *Korarchaeum*.

**Emended description of *Candidatus* Methanodesulfokora McKay et al. 2019**

Me.tha.no.de.sul.fo.ko’ra. N.L. neut. n. *methanum*, methane; N.L. pref. *methano-*, pertaining to methane; L. prep. *de*, from; N.L. pref. *sulfo-*, pertaining to sulfur; Gr. fem. n. *korê*, young woman; N.L. fem. n. *Methanodesulfokora*, a methane and sulfur metabolizing member of the Korarchaeota.

Estimated genome size of *Ca*. Methanodesulfokora MAGs range from 1.33 to 2.83 Mb, their genomic G+C content from 43.24 to 44.04 %. The representative MAG of the genus is that of *Ca*. Methanodesulfokora washburnensis NM4. Its NCBI accession number is RXII00000000.

**Emended description of *Candidatus* Methanodesulfokora washburnensis McKay et al. 2019**

wash.burn.en’sis. N.L. masc./fem. adj. *washburnensis*, pertaining to Washburn Hot Springs in Yellowstone National Park, USA.

The representative draft genome is that of *Ca*. Methanodesulfokora washburnensis NM4 (NCBI accession number RXII00000000). As derived from the draft genome the estimated genome size is 1.70 Mb, the DNA G+C content 43.43%.

**Description of *Candidatus* Methanodesulfokora zavarzinensis sp. nov.**

za.var.zin.en’sis. N.L. masc/fem. adj. *zavarzinensis*, pertaining to Zavarzin hot spring in the Uzon Caldera, Kamchatka, Russia.

The representative draft genome is that of *Ca*. Methanodesulfokora zavarzinensis ZAV-18 (NCBI accession number PNIB00000000). As derived from the draft genome the estimated genome size is 1.55 Mb, the DNA G+C content 44.04%.

**Description of *Candidatus* Fumariihydrokora gen. nov.**

Fu.ma.ri.i.hy.dro.ko’ra. L. neut. n. *fumarium*, chimney; Gr. neut. n. *hydôr*, water; Gr. fem. n. *korê*, young woman; N.L. fem. n. *Fumariihydrokora*, a member of the Korarchaeota found in a hydrothermal vent area.

Estimated genome size of *Ca*. Fumariihydrokora MAGs range from 1.15 to 1.84 Mb, their genomic G+C content from 43.83 to 50.25 %. The representative MAG of the genus is that of *Ca*. Fumariihydrokora guaymasensis HyVt-138. Its NCBI accession number is DQXR00000000.

**Description of *Candidatus* Fumariihydrokora guaymasensis sp. nov.**

guay.mas.en’sis. N.L. masc./fem. adj. *guaymasensis*, of or belonging to Guaymas, the location where the organism was discovered.

The representative draft genome is that of *Ca*. Fumariihydrokora guaymasensis HyVt-138 (NCBI accession number DQXR00000000). As derived from the draft genome the estimated genome size is 1.15 Mb, the DNA G+C content 50.25%.

**Description of *Candidatus* Fumariihydrokora hydrothermalis sp. nov.**

hy.dro.ther.ma’lis. Gr. neut. n. *hydôr*, water; Gr. masc. adj. *thermos*, hot; N.L. masc./fem. adj. *hydrothermalis*, to indicate that the organism originated from a hydrothermal vent.

The representative draft genome is that of *Ca*. Fumariihydrokora hydrothermalis B81_G16 (NCBI accession number QMVT00000000). As derived from the draft genome the estimated genome size is 1.84 Mb, the DNA G+C content 43.83%.

**Description of *Candidatus* Calidiprofundikora gen. nov.**

Ca.li.di.pro.fun.di.ko’ra. L. masc. adj. *calidus*, hot; L. masc. adj. *profundus*, pertaining to the depths of the sea; Gr. fem. n. *korê*, young woman; N.L. fem. n. *Calidiprofundikora*, a member of the Korarchaeota coming from a hot deep-sea area (i.e. a hydrothermal vent).

Estimated genome size of *Ca*. Calidiprofundikora MAGs range from 1.48 to 1.85 Mb, their genomic G+C content from 57.12 to 64.16 %. The representative MAG of the genus is that of *Ca*. Calidiprofundikora taketomiensis Kor2. Its NCBI accession number is JAVCBV000000000.

**Description of *Candidatus* Calidiprofundikora taketomiensis sp. nov.**

ta.ke.to.mi.en’sis. N.L. masc./fem. adj. *taketomiensis*, of or belonging to Taketomi, the location where the organism’s genome was recovered from.

The representative draft genome is that of *Ca*. Calidiprofundikora taketomiensis Kor2 (NCBI accession number JAVCBV000000000). As derived from the draft genome the estimated genome size is 1.48 Mb, the DNA G+C content 57.85%.

**Description of *Candidatus* Calidiprofundikora hydrothermalis sp. nov.**

hy.dro.ther.ma’lis. Gr. neut. n. *hydôr*, water; Gr. masc. adj. *thermos*, hot; N.L. masc./fem. adj. *hydrothermalis*, to indicate that the organism originated from a hydrothermal vent.

The representative draft genome is that of *Ca*. Calidiprofundikora hydrothermalis HyVt-239 (NCBI accession number DRBY00000000). As derived from the draft genome the estimated genome size is 1.53 Mb, the DNA G+C content 57.12%.

**Description of *Candidatus* Calidiprofundikora guaymasensis sp. nov.**

guay.mas.en’sis. N.L. masc./fem. adj. *guaymasensis*, of or belonging to Guaymas, the location where the organism was discovered.

The representative draft genome is that of *Ca*. Calidiprofundikora guaymasensis B85_G9 (NCBI accession number QMVS00000000). As derived from the draft genome the estimated genome size is 1.85 Mb, the DNA G+C content 59.35%. A second draft genome is that of MAG HyVt-161 (NCBI accession number DQYR00000000) which has an estimated genome size of 1.71 Mb and a DNA G+C content of 64.16%.

**Description of *Candidatus* Sedimenticalidikora gen. nov.**

Se.di.men.ti.ca.li.di.ko’ra. L. neut. n. *sedimentum*, sediment; L. masc. adj. *calidus*, hot; Gr. fem. n. *korê*, young woman; N.L. fem. n. *Sedimenticalidikora*, a Korarchaeota archaeon originating from hot sediment.

The representative draft genome is that of *Ca*. Sedimenticalidikora guaymasensis B35_G17 (NCBI accession number QMVY00000000).

**Description of *Candidatus* Sedimenticalidikora guaymasensis sp. nov.**

guay.mas.en’sis. N.L. masc./fem. adj. *guaymasensis*, of or belonging to Guaymas, the location where the organism was discovered.

The representative draft genome is that of *Ca*. Sedimenticalidikora guaymasensis B35_G17 (NCBI accession number QMVY00000000). As derived from the draft genome the estimated genome size is 1.24 Mb, the DNA G+C content 65.01%.

**Description of *Candidatus* Hydrocaminikora gen. nov.**

Hy.dro.ca.mi.ni.ko’ra. Gr. neut. n. *hydôr*, water; L. gen. masc. n. *caminus*, a chimney; Gr. fem. n. *korê*, young woman; N.L. fem. n. *Hydrocaminikora*, a Korarchaeota archaeon from a hydrothermal chimney.

Estimated genome size of *Ca*. Hydrocaminikora MAGs range from 1.10 to 2.48 Mb, their genomic G+C content from 43.56 to 47.46 %. The representative MAG of the genus is that of *Ca*. Hydrocaminikora taketomiensis Kor4. Its NCBI accession number is JAVCBX000000000.

**Description of *Candidatus* Hydrocaminikora taketomiensis Kor4 sp. nov.**

ta.ke.to.mi.en’sis. N.L. masc./fem. adj. *taketomiensis*, of or belonging to Taketomi, the location where the organism’s genome was recovered from.

The representative draft genome is that of *Ca*. Hydrocaminikora taketomiensis Kor4 (NCBI accession number JAVCBX000000000). As derived from the draft genome the estimated genome size is 1.45 Mb, the DNA G+C content 47.46%.

**Description of *Candidatus* Hydrocaminikora hydrothermalis sp. nov.**

hy.dro.ther.ma’lis. Gr. neut. n. *hydôr*, water; Gr. masc. adj. *thermos*, hot; N.L. masc./fem. adj. *hydrothermalis*, to indicate that the organism originated from a hydrothermal vent.

The representative draft genome is that of *Ca*. Hydrocaminikora hydrothermalis B8_G17 (NCBI accession number QMVU00000000). As derived from the draft genome the estimated genome size is 2.04 Mb, the DNA G+C content 46.86%. A second draft genome is that of MAG HyVt-231 (NCBI accession number DRBQ00000000) which has an estimated genome size of 1.10 Mb and a DNA G+C content of 45.96%.

**Description of *Candidatus* Hydrocaminikora caymanensis sp. nov.**

cay.man.en’sis. N.L. masc./fem. adj. *caymanensis*, of or belonging to the Mid-Cayman Rise hydrothermal vent.

The representative draft genome is that of *Ca*. Hydrocaminikora caymanensis UWMA-0234 (NCBI accession number DTYZ00000000). As derived from the draft genome the estimated genome size is 2.24 Mb, the DNA G+C content 43.59%.

**Description of *Candidatus* Hydrocaminikora guaymasensis sp. nov.**

guay.mas.en’sis. N.L. masc./fem. adj. *guaymasensis*, of or belonging to Guaymas, the location where the organism was discovered.

The representative draft genome is that of *Ca*. Hydrocaminikora guaymasensis B15_G15 (NCBI accession number QMVZ00000000). As derived from the draft genome the estimated genome size is 2.28 Mb, the DNA G+C content 43.56%. Two other MAGs (QMVX00000000 and QMVW00000000) have estimated genome sizes and DNA G+C contents of 2.09 and 2.48 MB, and 44.04 and 43.92% respectively.

**Description of *Candidatus* Hydrocaminikoraceae fam. nov.**

*Hydrocaminikoraceae* (Hy.dro.ca.mi.ni.ko.ra.ce’ae, N.L. fem. n. *Hydrocaminikora*, the type genus of the family; suff. -*aceae*, ending to denote a family; N.L. fem. pl. n. *Hydrocaminikoraceae*, the family of the genus *Hydrocaminikora*).

The description is the same as for the genus *Hydrocaminikora*. Type genus is *Hydrocaminikora*.

**Description of *Candidatus* Thermotainarokora gen. nov.**

Ther.mo.tai.na.ro.ko’ra. Gr. masc. adj. *thermos*, hot; Gr. neut. n. *Tainaron*, a town named after Tainaros, a son of Zeus. Tainaron was believed to be the gate to the Underworld; Gr. fem. n. *korê*, young woman; N.L. fem. n. *Thermotainarokora*, a member of the Korarchaeota from a hot underground site.

Estimated genome size of *Ca*. Thermotainarokora MAGs range from 1.85 to 1.93 Mb, their genomic G+C content from 42.30 to 44.76 %. The representative MAG of the genus is that of *Ca*. Thermotainarokora taketomiensis Kor1. Its NCBI accession number is JAVCBU000000000.

**Description of *Candidatus* Thermotainarokora taketomiensis Kor1 sp. nov.**

ta.ke.to.mi.en’sis. N.L. masc./fem. adj. *taketomiensis*, of or belonging to Taketomi, the location where the organism’s genome was recovered from.

The representative draft genome is that of *Ca*. Thermotainarokora taketomiensis Kor1 (NCBI accession number JAVCBU000000000). As derived from the draft genome the estimated genome size is 1.85 Mb, the DNA G+C content 44.76%.

**Description of *Candidatus* Thermotainarokora guaymasensis sp. nov.**

guay.mas.en’sis. N.L. masc./fem. adj. *guaymasensis*, of or belonging to Guaymas, the location where the organism was discovered.

The representative draft genome is that of *Ca*. Thermotainarokora guaymasensis B68_G1 (NCBI accession number QMVV00000000). As derived from the draft genome the estimated genome size is 1.93 Mb, the DNA G+C content 42.30%.

**Description of *Candidatus* Thermotainarokoraceae fam. nov.**

*Thermotainarokora*ceae (Ther.mo.tai.na.ro.co.ra.ce’ae, N.L. fem. n. *Thermotainarokora*, the type genus of the family; suff. -*aceae*, ending to denote a family; N.L. fem. pl. n. *Thermotainarokoraceae*, the family of the genus *Thermotainarokora*).

The description is the same as for the genus *Thermotainarokora*. Type genus is *Thermotainarokora*.

**Description of *Candidatus* Caldabyssikora gen. nov.**

Cal.da.bys.si.ko’ra. L. masc. adj. *caldus*, hot; L. fem. n. *abyssus*, an abyss, the deep sea; Gr. fem. n. *korê*, young woman; N.L. fem. n. *Caldabyssikora*, a Korarchaeon from the warm deep sea, referring to the hydrothermal vent areas where the organisms are found.

Estimated genome size of *Ca*. Caldabyssikora MAGs range from 1.88 to 2.02 Mb, their genomic G+C content from 49.71 to 52.84 %. The representative MAG of the genus is that of *Ca*. Caldabyssikora taketomiensis Kor3. Its NCBI accession number is JAVCBW000000000.

**Description of *Candidatus* Caldabyssikora taketomiensis sp. nov.**

ta.ke.to.mi.en’sis. N.L. masc./fem. adj. *taketomiensis*, of or belonging to Taketomi, the location where the organism’s genome was recovered from.

The representative draft genome is that of *Ca*. Caldabyssikora taketomiensis Kor3 (NCBI accession number JAVCBW000000000). As derived from the draft genome the estimated genome size is 1.88 Mb, the DNA G+C content 49.71%.

**Description of *Candidatus* Caldabyssikora guaymasensis sp. nov.**

guay.mas.en’sis. N.L. masc./fem. adj. *guaymasensis*, of or belonging to Guaymas, the location where the organism was discovered.

The representative draft genome is that of *Ca*. Caldabyssikora guaymasensis B14_G2 (NCBI accession number QMWA00000000). As derived from the draft genome the estimated genome size is 1.88 Mb, the DNA G+C content 52.83%. A second draft genome is that of MAG B10_G17 (NCBI accession number QMWB00000000) which has an estimated genome size of 2.02 Mb and a DNA G+C content of 52.84%.

**Description of *Candidatus* Caldabyssikoraceae fam. nov.**

*Caldabyssikoraceae* (Cal.da.bys.si.ko.ra.ce’ae, N.L. fem. n. *Caldabyssikora*, the type genus of the family; suff. -*aceae*, ending to denote a family; N.L. fem. pl. n. *Caldabyssikoraceae*, the family of the genus *Caldabyssikora*).

The description is the same as for the genus *Caldabyssikora*. Type genus is *Caldabyssikora*.
