## Supplementary Figures S1-S4 for "Phylogenomics and ancestral reconstruction of Korarchaeota reveals genomic adaptation to habitat switching"

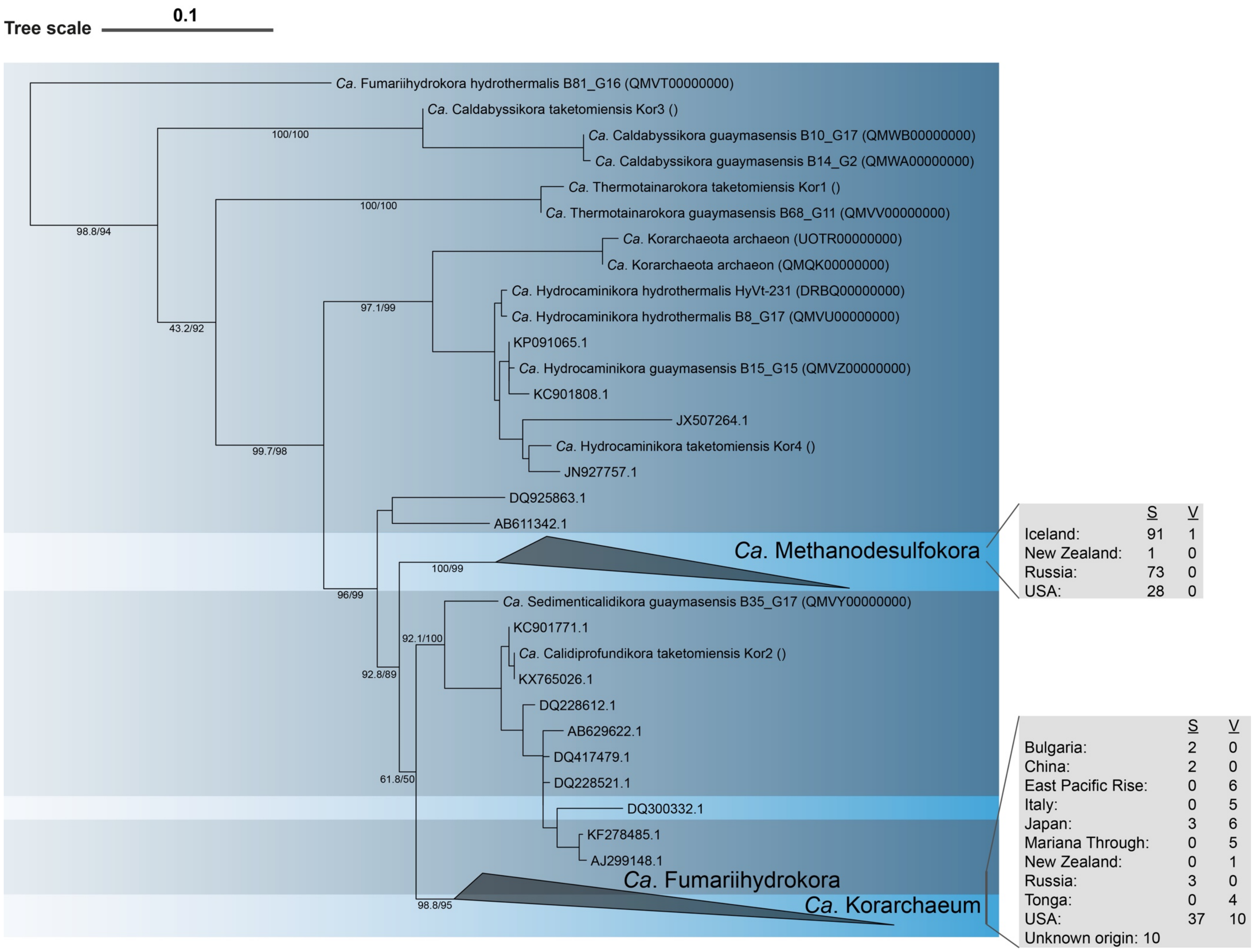
Tree scale
0.1
Ca. Fumariihydrokora hydrothermalis B81_G16 (QMVT00000000)
Ca. Caldabyssikora taketomiensis Kor3 ()
Ca. Caldabyssikora guaymasensis B10_G17 (QMWB00000000)
Ca. Caldabyssikora guaymasensis B14_G2 (QMWA00000000)
Ca. Thermotainarokora taketomiensis Kor1 ()
Ca. Thermotainarokora guaymasensis B68_G11 (QMVV00000000)
Ca. Korarchaeota archaeon (UOTR00000000)
Ca. Korarchaeota archaeon (QMQK00000000)
Ca. Hydrocaminikora hydrothermalis HyVt-231 (DRBQ00000000)
Ca. Hydrocaminikora hydrothermalis B8_G17 (QMVU00000000)
KP091065.1
Ca. Hydrocaminikora guaymasensis B15_G15 (QMVZ00000000)
KC901808.1
JX507264.1
Ca. Hydrocaminikora taketomiensis Kor4 ()
JN927757.1
DQ925863.1
AB611342.1
Ca. Methanodesulfokora
Ca. Sedimenticalidikora guaymasensis B35_G17 (QMVY00000000)
KC901771.1
Ca. Calidiprofundikora taketomiensis Kor2 ()
KX765026.1
DQ228612.1
AB629622.1
DQ417479.1
DQ228521.1
DQ300332.1
KF278485.1
AJ299148.1
Ca. Fumariihydrokora
Ca. Korarchaeum
100/100
98.8/94
100/100
43.2/92
97.1/99
99.7/98
100/99
96/99
92.1/100
92.8/89
61.8/50
98.8/95
S V
Iceland: 91 1
New Zealand: 1 0
Russia: 73 0
USA: 28 0
S V
Bulgaria: 2 0
China: 2 0
East Pacific Rise: 0 6
Italy: 0 5
Japan: 3 6
Mariana Through: 0 5
New Zealand: 0 1
Russia: 3 0
Tonga: 0 4
USA: 37 10
Unknown origin: 10

**Fig. S1.** Maximum-likelihood phylogenetic tree (IQ-TREE, GTR+F model, 1000 ultrafast bootstrap replicates, 1000 approximate likelihood-ratio test) of 16S rRNA gene sequences of *Ca*. Korarchaeota retrieved from SILVA 138.1 (Table S3) and MAGs (Table S2). Scale bar indicates 0.1 substitutions per nucleotide position. For collapsed clades, an overview of the number of sequences from hot springs (S) and hydrothermal vents (V) is shown together with their geographical origin. *Candidatus* Korarchaeota clades originating from hot springs and hydrothermal vents have a light and dark blue background shading, respectively.

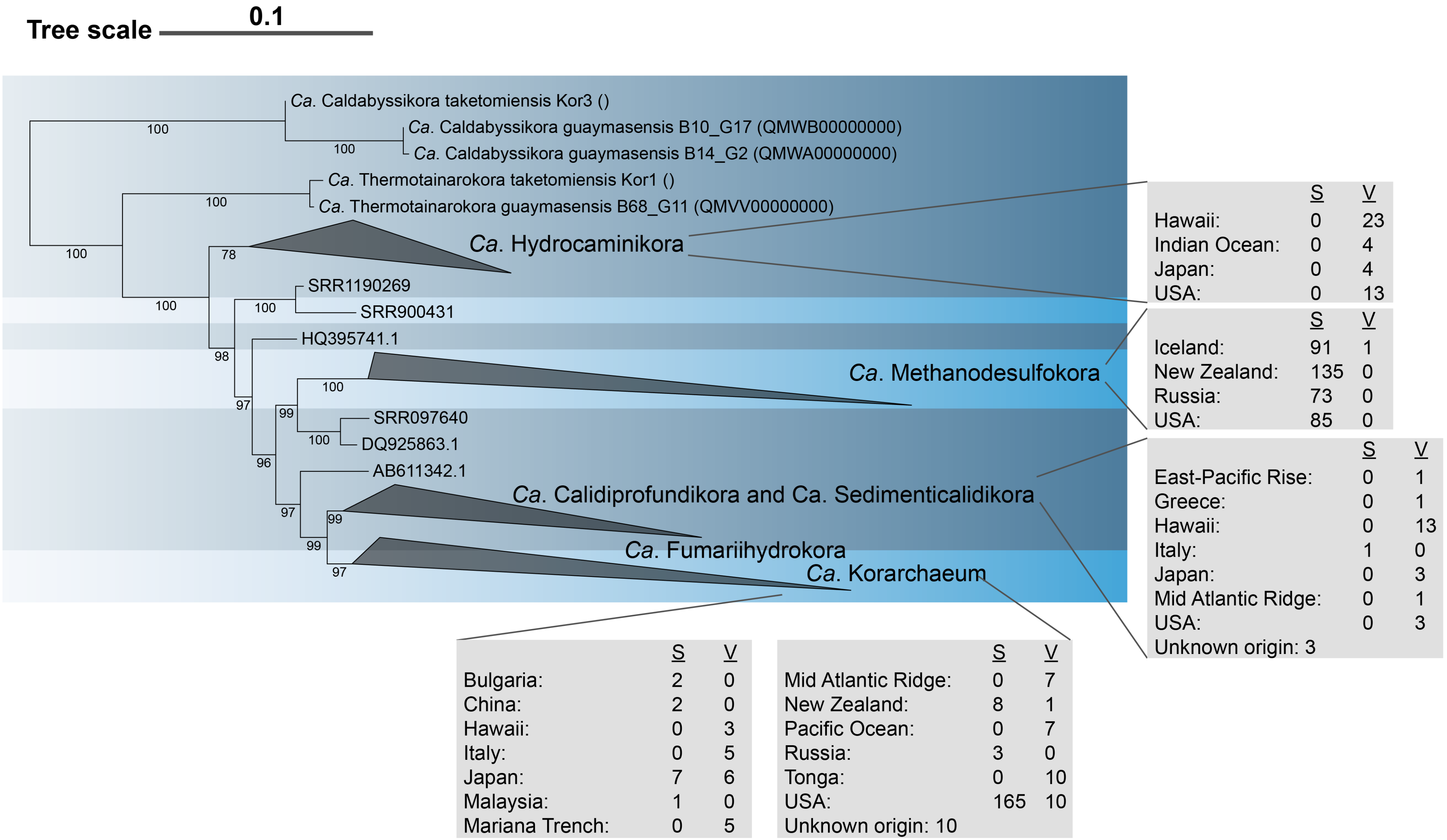


**Fig. S2.** Maximum-likelihood phylogenetic tree (IQ-TREE, GTR+F model, 1000 ultrafast bootstrap replicates) of 16S rRNA gene sequences of *Ca.* Korarchaeota retrieved from SILVA 138.1 (Table S3), MAGs (Table S2) and IMNGS datasets (Table S4). Scale bar indicates 0.1 substitutions per nucleotide position. For collapsed clades, an overview of the number of sequences from hot springs (S) and hydrothermal vents (V) is shown together with their geographical origin. *Candidatus* Korarchaeota clades originating from hot springs and hydrothermal vents have a light and dark blue background shading, respectively.

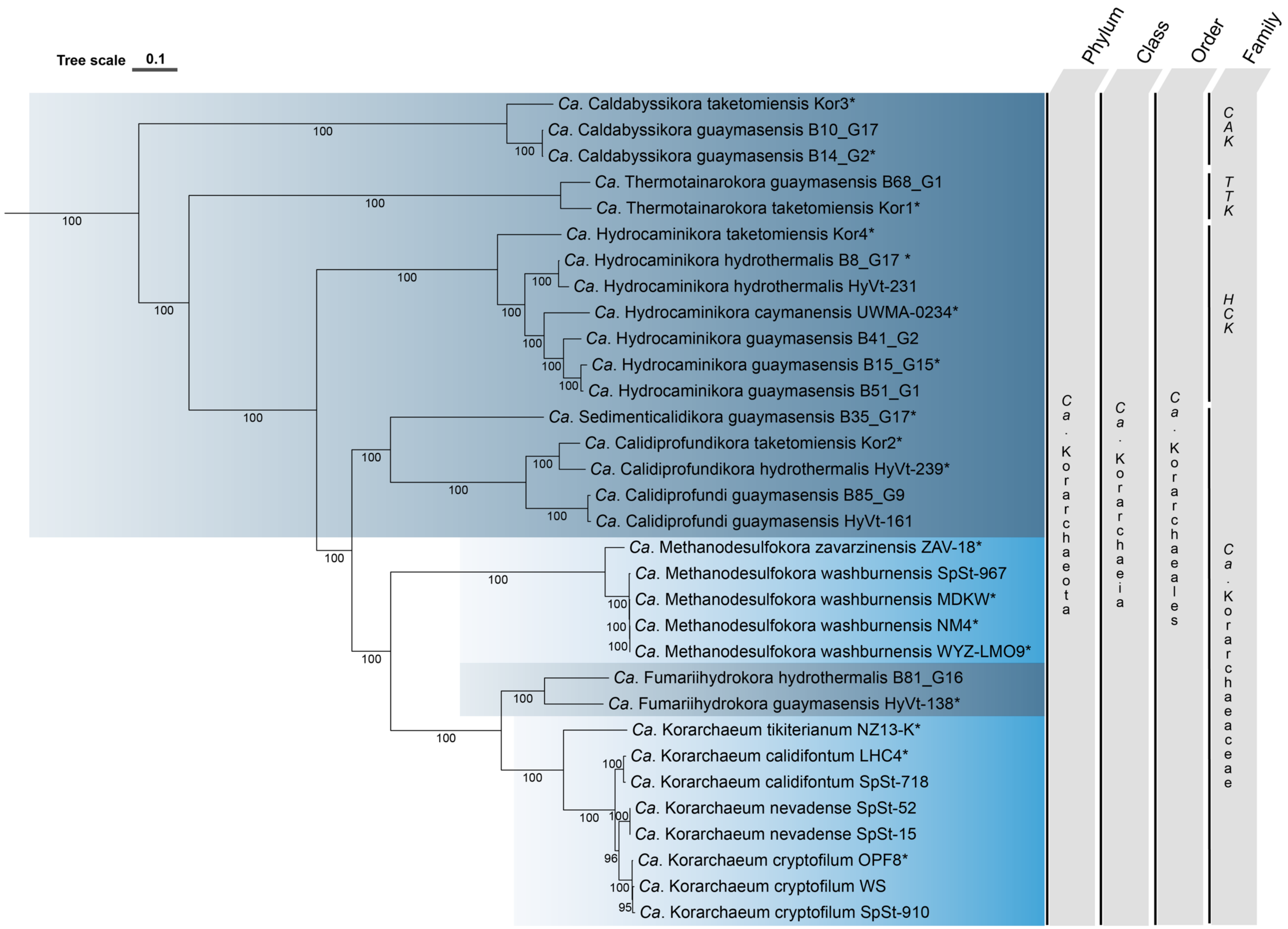

Tree scale 0.1
Phylum
Class
Order
Family
Ca. Caldabyssikora taketomiensis Kor3*
Ca. Caldabyssikora guaymasensis B10_G17
Ca. Caldabyssikora guaymasensis B14_G2*
Ca. Thermotainarokora guaymasensis B68_G1
Ca. Thermotainarokora taketomiensis Kor1*
Ca. Hydrocaminikora taketomiensis Kor4*
Ca. Hydrocaminikora hydrothermalis B8_G17 *
Ca. Hydrocaminikora hydrothermalis HyVt-231
Ca. Hydrocaminikora caymanensis UWMA-0234*
Ca. Hydrocaminikora guaymasensis B41_G2
Ca. Hydrocaminikora guaymasensis B15_G15*
Ca. Hydrocaminikora guaymasensis B51_G1
Ca. Sedimenticalidikora guaymasensis B35_G17*
Ca. Calidiprofundikora taketomiensis Kor2*
Ca. Calidiprofundikora hydrothermalis HyVt-239*
Ca. Calidiprofundi guaymasensis B85_G9
Ca. Calidiprofundi guaymasensis HyVt-161
Ca. Methanodesulfokora zavarzinensis ZAV-18*
Ca. Methanodesulfokora washburnensis SpSt-967
Ca. Methanodesulfokora washburnensis MDKW*
Ca. Methanodesulfokora washburnensis NM4*
Ca. Methanodesulfokora washburnensis WYZ-LMO9*
Ca. Fumariihydrokora hydrothermalis B81_G16
Ca. Fumariihydrokora guaymasensis HyVt-138*
Ca. Korarchaeum tikiterianum NZ13-K*
Ca. Korarchaeum calidifontum LHC4*
Ca. Korarchaeum calidifontum SpSt-718
Ca. Korarchaeum nevadense SpSt-52
Ca. Korarchaeum nevadense SpSt-15
Ca. Korarchaeum cryptofilum OPF8*
Ca. Korarchaeum cryptofilum WS
Ca. Korarchaeum cryptofilum SpSt-910
Ca. Korarchaeota
Ca. Korarchaeia
Ca. Korarchaeales
CAK
TTK
HCK
Ca. Korarchaeaceae
100
96
95

**Fig. S3.** Maximum-likelihood phylogenetic tree (IQ-TREE, LG+C60+F+G4 model, 1000 ultrafast bootstrap replicates) of *Ca*. Korarchaeota MAGs based on the concatenation of 76 core archaeal single-copy genes (1). *Candidatus* Korarchaeota clades originating from hot springs and hydrothermal vents have a light and dark blue background shading, respectively. Scale bar indicates 0.1 substitutions per position. Branch names ending with an * indicate the MAG was included in the dataset for the amalgamated likelihood estimation analysis. An overview of the taxonomic descriptions of new taxa is given in the Supplementary Note. HCK: *Candidatus* Hydrocaminikoraceae; TTK: *Candidatus* Thermotainarokora; CAK: *Candidatus* Caldabyssikora

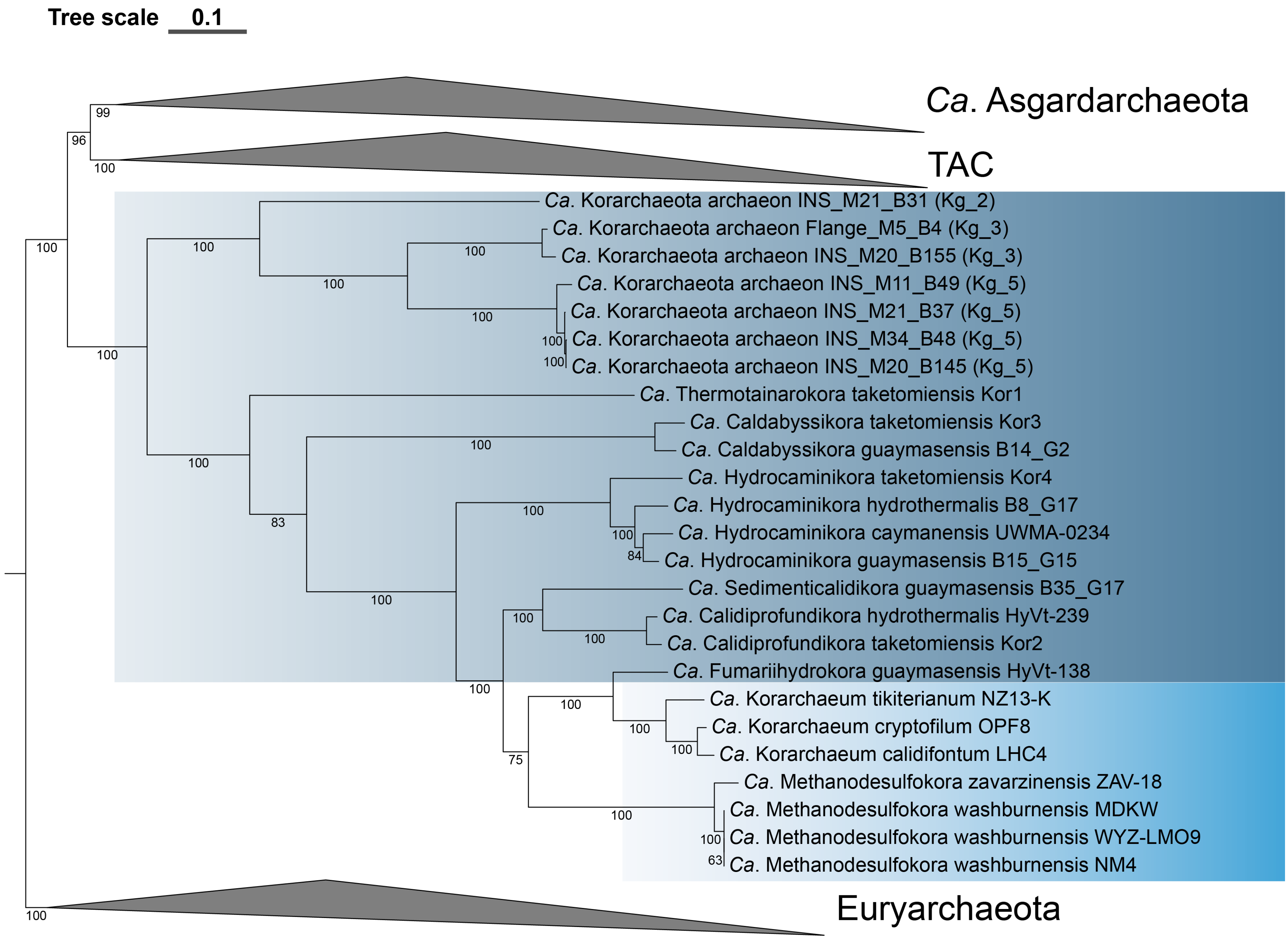

Tree scale 0.1
Ca. Asgardarchaeota
TAC
Ca. Korarchaeota archaeon INS_M21_B31 (Kg_2)
Ca. Korarchaeota archaeon Flange_M5_B4 (Kg_3)
Ca. Korarchaeota archaeon INS_M20_B155 (Kg_3)
Ca. Korarchaeota archaeon INS_M11_B49 (Kg_5)
Ca. Korarchaeota archaeon INS_M21_B37 (Kg_5)
Ca. Korarchaeota archaeon INS_M34_B48 (Kg_5)
Ca. Korarchaeota archaeon INS_M20_B145 (Kg_5)
Ca. Thermotainarokora taketomiensis Kor1
Ca. Caldabyssikora taketomiensis Kor3
Ca. Caldabyssikora guaymasensis B14_G2
Ca. Hydrocaminikora taketomiensis Kor4
Ca. Hydrocaminikora hydrothermalis B8_G17
Ca. Hydrocaminikora caymanensis UWMA-0234
Ca. Hydrocaminikora guaymasensis B15_G15
Ca. Sedimenticalidikora guaymasensis B35_G17
Ca. Calidiprofundikora hydrothermalis HyVt-239
Ca. Calidiprofundikora taketomiensis Kor2
Ca. Fumariihydrokora guaymasensis HyVt-138
Ca. Korarchaeum tikiterianum NZ13-K
Ca. Korarchaeum cryptofilum OPF8
Ca. Korarchaeum calidifontum LHC4
Ca. Methanodesulfokora zavarzinensis ZAV-18
Ca. Methanodesulfokora washburnensis MDKW
Ca. Methanodesulfokora washburnensis WYZ-LMO9
Ca. Methanodesulfokora washburnensis NM4
Euryarchaeota

**Fig. S4.** Non-parametric bootstrap maximum-likelihood phylogenomic tree (IQ-TREE, 100 bootstraps, LG+C60+F+G4 model) based on the χ2-trimmed alignment of 54 concatenated new marker proteins (NM54; 7584 alignment positions). The dataset included the original 137 MAGs (Tables S1-S2), *Ca*. Njord- and *Ca*. Panguiarchaeales MAGs, and seven MAGs, corresponding to the lineages Kg_2, Kg_3 and Kg_5, from the study of F. Vulcano et al. (2). Scale bar indicates 0.1 substitutions per site. Candidatus Korarchaeota originating from hot springs and hydrothermal vents have a light and dark blue background shading, respectively.
