## Supplementary Figures S5-S15 for "Phylogenomics and ancestral reconstruction of Korarchaeota reveals genomic adaptation to habitat switching"

*** Correspondence:**

Guillaume Tahon

Archaea, microbial diversity, phylogeny, hyperthermophiles, hot spring, genome evolution

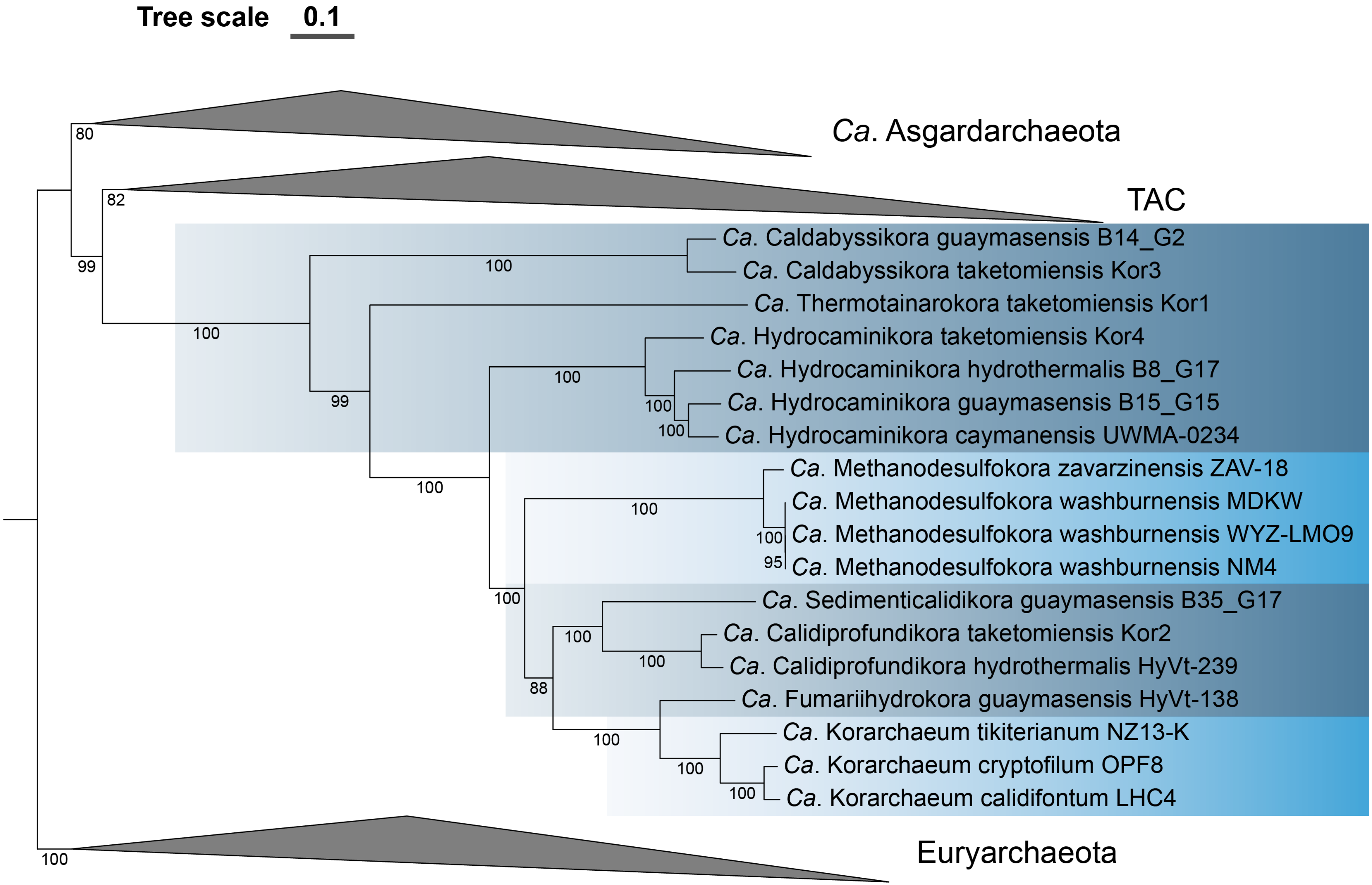

Tree scale 0.1
Ca. Asgardarchaeota
TAC
Ca. Caldabyssikora guaymasensis B14_G2
Ca. Caldabyssikora taketomiensis Kor3
Ca. Thermotainarokora taketomiensis Kor1
Ca. Hydrocaminikora taketomiensis Kor4
Ca. Hydrocaminikora hydrothermalis B8_G17
Ca. Hydrocaminikora guaymasensis B15_G15
Ca. Hydrocaminikora caymanensis UWMA-0234
Ca. Methanodesulfokora zavarzinensis ZAV-18
Ca. Methanodesulfokora washburnensis MDKW
Ca. Methanodesulfokora washburnensis WYZ-LMO9
Ca. Methanodesulfokora washburnensis NM4
Ca. Sedimenticalidikora guaymasensis B35_G17
Ca. Calidiprofundikora taketomiensis Kor2
Ca. Calidiprofundikora hydrothermalis HyVt-239
Ca. Fumariihydrokora guaymasensis HyVt-138
Ca. Korarchaeum tikiterianum NZ13-K
Ca. Korarchaeum cryptofilum OPF8
Ca. Korarchaeum calidifontum LHC4
Euryarchaeota

**Fig. S5.** Non-parametric bootstrap maximum-likelihood phylogenomic tree (IQ-TREE, 100 bootstraps, LG+C60+F+G4 model) based on the untreated alignment of 56 concatenated ribosomal proteins (RP56; 7158 alignment positions). Scale bar indicates 0.1 substitutions per site. *Candidatus* Korarchaeota originating from hot springs and hydrothermal vents have a light and dark blue background shading, respectively.

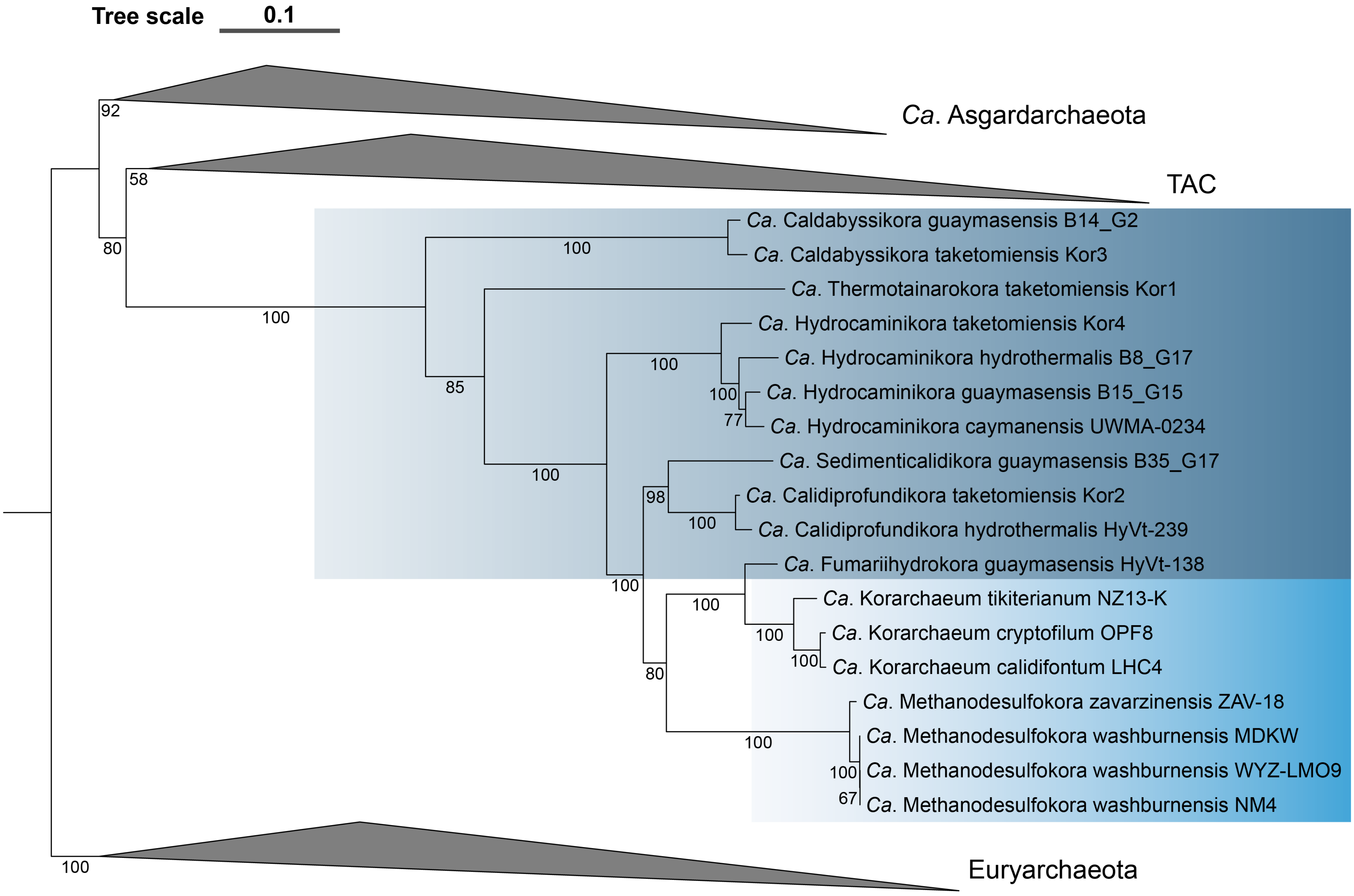
Tree scale
0.1
Ca. Asgardarchaeota
TAC
Ca. Caldabyssikora guaymasensis B14_G2
Ca. Caldabyssikora taketomiensis Kor3
Ca. Thermotainarokora taketomiensis Kor1
Ca. Hydrocaminikora taketomiensis Kor4
Ca. Hydrocaminikora hydrothermalis B8_G17
Ca. Hydrocaminikora guaymasensis B15_G15
Ca. Hydrocaminikora caymanensis UWMA-0234
Ca. Sedimenticalidikora guaymasensis B35_G17
Ca. Calidiprofundikora taketomiensis Kor2
Ca. Calidiprofundikora hydrothermalis HyVt-239
Ca. Fumariihydrokora guaymasensis HyVt-138
Ca. Korarchaeum tikiterianum NZ13-K
Ca. Korarchaeum cryptofilum OPF8
Ca. Korarchaeum calidifontum LHC4
Ca. Methanodesulfokora zavarzinensis ZAV-18
Ca. Methanodesulfokora washburnensis MDKW
Ca. Methanodesulfokora washburnensis WYZ-LMO9
Ca. Methanodesulfokora washburnensis NM4
Euryarchaeota
92
58
80
100
100
85
100
100
100
77
98
100
100
100
100
80
100
100
100
67
100

**Fig. S6.** Non-parametric bootstrap maximum-likelihood phylogenomic tree (IQ-TREE, 100 bootstraps, GTR+G4+C60SR4 model) based on an SR4-recoded concatenation of 56 ribosomal proteins (RP56; 7158 alignment positions). Scale bar indicates 0.1 substitutions per site. *Candidatus* Korarchaeota originating from hot springs and hydrothermal vents have a light and dark blue background shading, respectively.

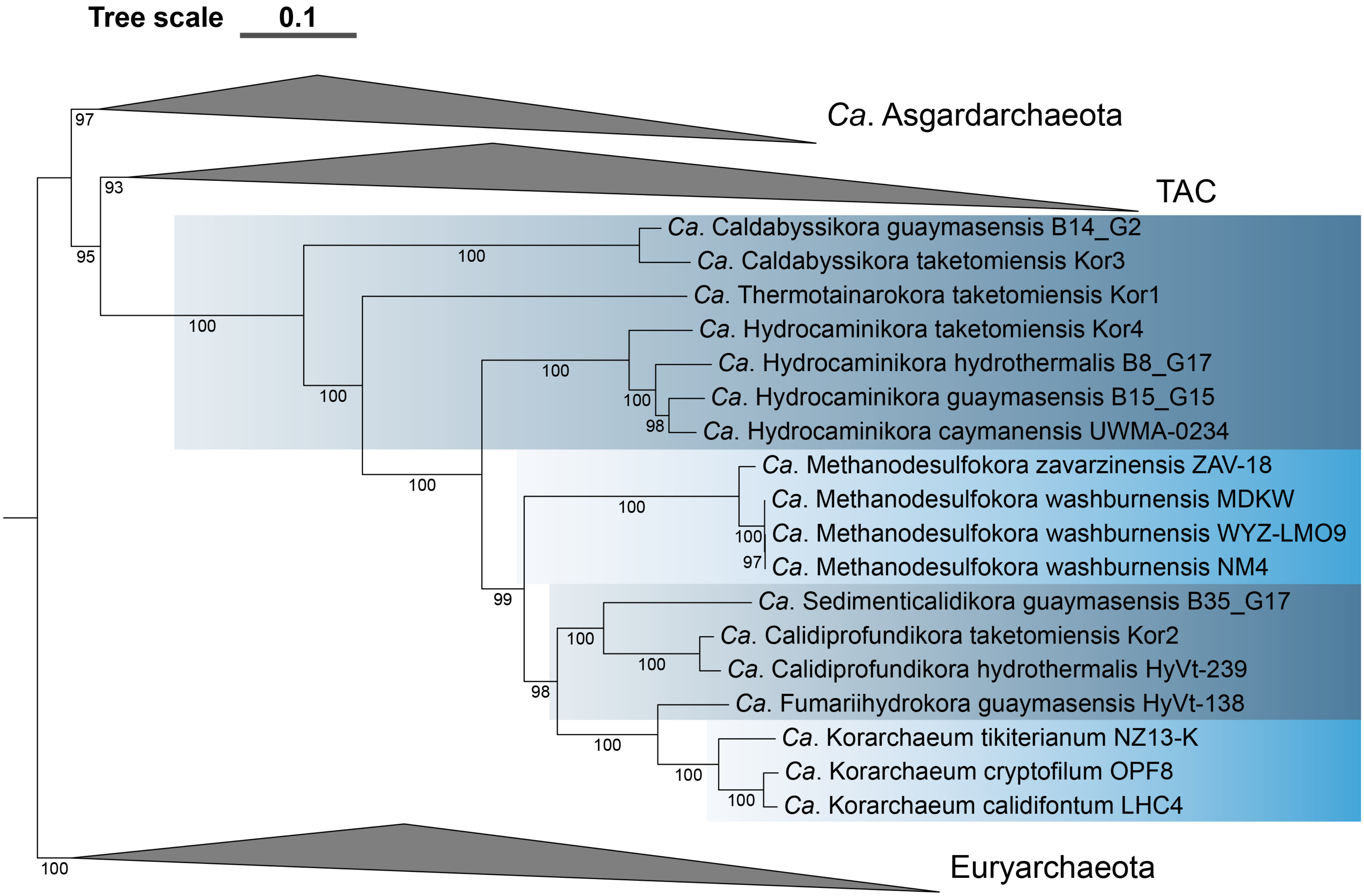

Tree scale 0.1
Ca. Asgardarchaeota
TAC
Ca. Caldabyssikora guaymasensis B14_G2
Ca. Caldabyssikora taketomiensis Kor3
Ca. Thermotainarokora taketomiensis Kor1
Ca. Hydrocaminikora taketomiensis Kor4
Ca. Hydrocaminikora hydrothermalis B8_G17
Ca. Hydrocaminikora guaymasensis B15_G15
Ca. Hydrocaminikora caymanensis UWMA-0234
Ca. Methanodesulfokora zavarzinensis ZAV-18
Ca. Methanodesulfokora washburnensis MDKW
Ca. Methanodesulfokora washburnensis WYZ-LMO9
Ca. Methanodesulfokora washburnensis NM4
Ca. Sedimenticalidikora guaymasensis B35_G17
Ca. Calidiprofundikora taketomiensis Kor2
Ca. Calidiprofundikora hydrothermalis HyVt-239
Ca. Fumariihydrokora guaymasensis HyVt-138
Ca. Korarchaeum tikiterianum NZ13-K
Ca. Korarchaeum cryptofilum OPF8
Ca. Korarchaeum calidifontum LHC4
Euryarchaeota

**Fig. S7.** Non-parametric bootstrap maximum-likelihood phylogenomic tree (IQ-TREE, 100 bootstraps, LG+C60+F+G4 model) based on a custom recoded concatenation of 56 ribosomal proteins (RP56; 7158 alignment positions). Scale bar indicates 0.1 substitutions per site. *Candidatus* Korarchaeota originating from hot springs and hydrothermal vents have a light and dark blue background shading, respectively.

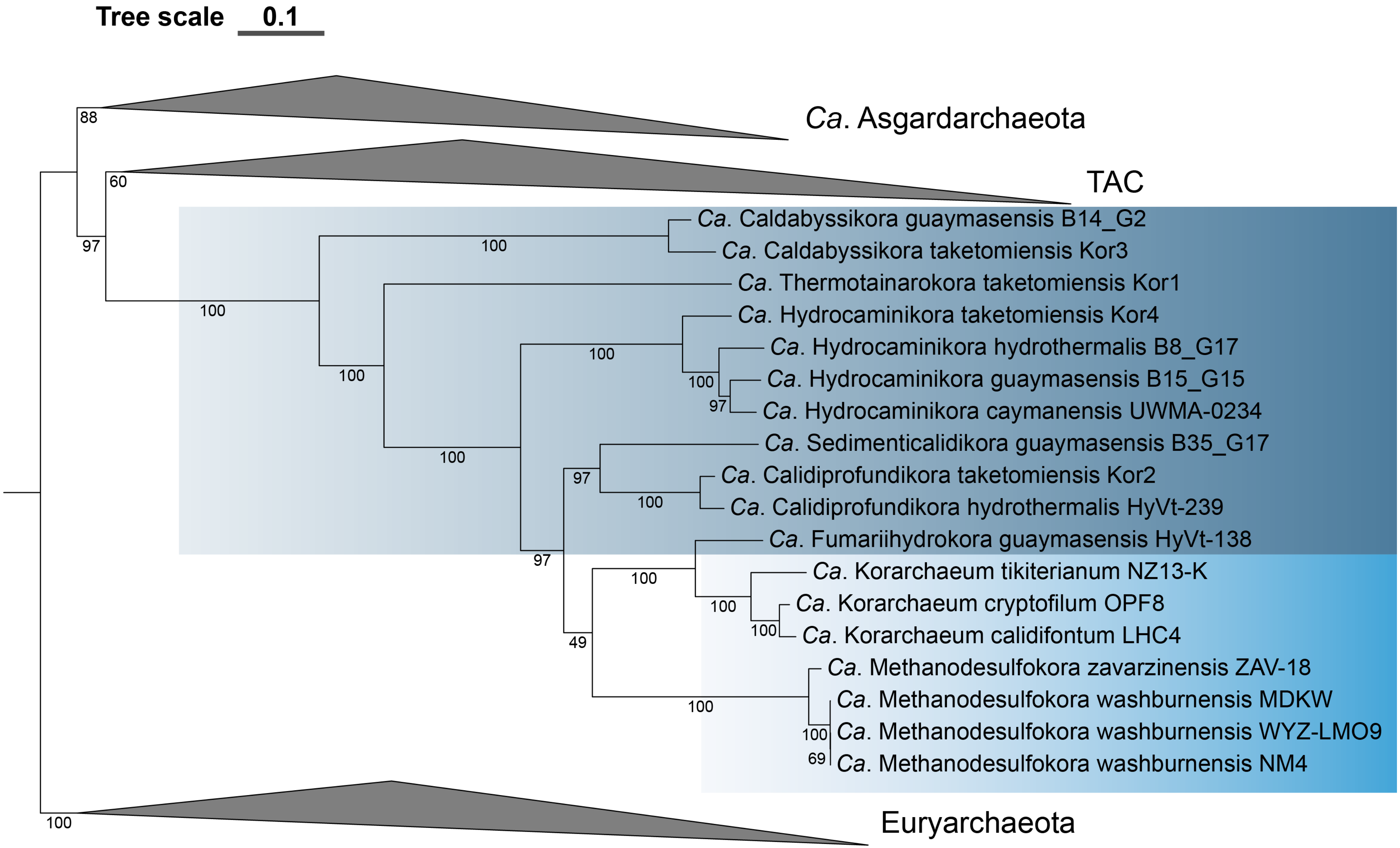
Tree scale 0.1
Ca. Asgardarchaeota
TAC
Ca. Caldabyssikora guaymasensis B14_G2
Ca. Caldabyssikora taketomiensis Kor3
Ca. Thermotainarokora taketomiensis Kor1
Ca. Hydrocaminikora taketomiensis Kor4
Ca. Hydrocaminikora hydrothermalis B8_G17
Ca. Hydrocaminikora guaymasensis B15_G15
Ca. Hydrocaminikora caymanensis UWMA-0234
Ca. Sedimenticalidikora guaymasensis B35_G17
Ca. Calidiprofundikora taketomiensis Kor2
Ca. Calidiprofundikora hydrothermalis HyVt-239
Ca. Fumariihydrokora guaymasensis HyVt-138
Ca. Korarchaeum tikiterianum NZ13-K
Ca. Korarchaeum cryptofilum OPF8
Ca. Korarchaeum calidifontum LHC4
Ca. Methanodesulfokora zavarzinensis ZAV-18
Ca. Methanodesulfokora washburnensis MDKW
Ca. Methanodesulfokora washburnensis WYZ-LMO9
Ca. Methanodesulfokora washburnensis NM4
Euryarchaeota
88
60
97
100
100
100
100
100
97
100
97
100
97
100
49
100
100
100
100
69
100

**Fig. S8.** Non-parametric bootstrap maximum-likelihood phylogenomic tree (IQ-TREE, 100 bootstraps, LG+C60+F+G4 model) based on a $\chi 2$-trimmed alignment (50% most heterogeneous sites removed, 3602 alignment positions) of 56 concatenated ribosomal proteins (RP56). Scale bar indicates 0.1 substitutions per site. *Candidatus* Korarchaeota originating from hot springs and hydrothermal vents have a light and dark blue background shading, respectively.

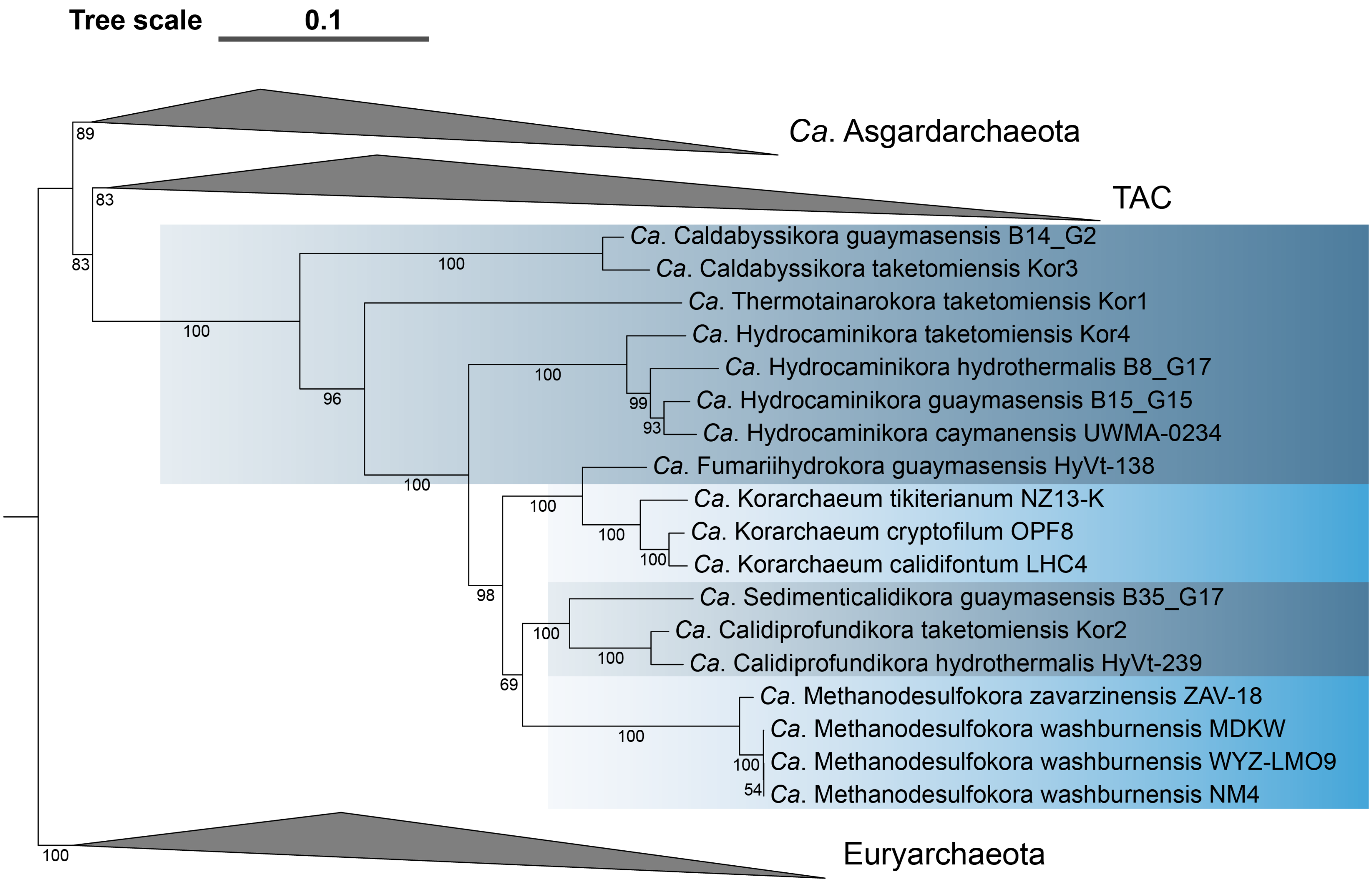

Tree scale
0.1
Ca. Asgardarchaeota
TAC
Ca. Caldabyssikora guaymasensis B14_G2
Ca. Caldabyssikora taketomiensis Kor3
Ca. Thermotainarokora taketomiensis Kor1
Ca. Hydrocaminikora taketomiensis Kor4
Ca. Hydrocaminikora hydrothermalis B8_G17
Ca. Hydrocaminikora guaymasensis B15_G15
Ca. Hydrocaminikora caymanensis UWMA-0234
Ca. Fumariihydrokora guaymasensis HyVt-138
Ca. Korarchaeum tikiterianum NZ13-K
Ca. Korarchaeum cryptofilum OPF8
Ca. Korarchaeum calidifontum LHC4
Ca. Sedimenticalidikora guaymasensis B35_G17
Ca. Calidiprofundikora taketomiensis Kor2
Ca. Calidiprofundikora hydrothermalis HyVt-239
Ca. Methanodesulfokora zavarzinensis ZAV-18
Ca. Methanodesulfokora washburnensis MDKW
Ca. Methanodesulfokora washburnensis WYZ-LMO9
Ca. Methanodesulfokora washburnensis NM4
Euryarchaeota
89
83
83
100
100
96
100
100
99
93
100
100
100
100
98
100
100
69
100
100
54
100

**Fig. S9.** Non-parametric bootstrap maximum-likelihood phylogenomic tree (IQ-TREE, 100 bootstraps, LG+C60+F+G4 model) based on a fast site trimmed alignment (50% fastest evolving sites removed, 3575 alignment positions) of 56 concatenated ribosomal proteins (RP56). Scale bar indicates 0.1 substitutions per site. *Candidatus* Korarchaeota originating from hot springs and hydrothermal vents have a light and dark blue background shading, respectively.

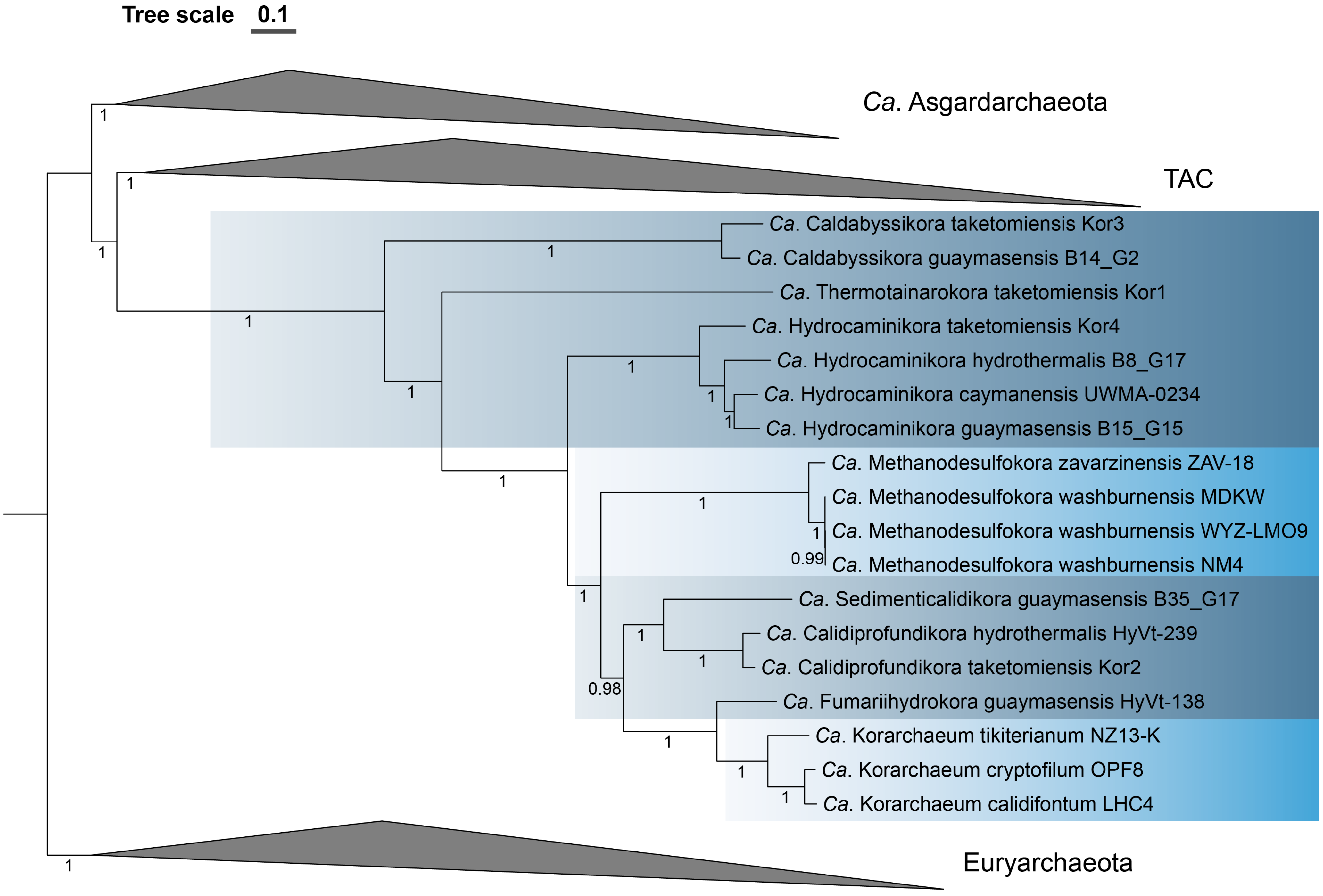

Tree scale 0.1
Ca. Asgardarchaeota
TAC
Ca. Caldabyssikora taketomiensis Kor3
Ca. Caldabyssikora guaymasensis B14_G2
Ca. Thermotainarokora taketomiensis Kor1
Ca. Hydrocaminikora taketomiensis Kor4
Ca. Hydrocaminikora hydrothermalis B8_G17
Ca. Hydrocaminikora caymanensis UWMA-0234
Ca. Hydrocaminikora guaymasensis B15_G15
Ca. Methanodesulfokora zavarzinensis ZAV-18
Ca. Methanodesulfokora washburnensis MDKW
Ca. Methanodesulfokora washburnensis WYZ-LMO9
Ca. Methanodesulfokora washburnensis NM4
Ca. Sedimenticalidikora guaymasensis B35_G17
Ca. Calidiprofundikora hydrothermalis HyVt-239
Ca. Calidiprofundikora taketomiensis Kor2
Ca. Fumariihydrokora guaymasensis HyVt-138
Ca. Korarchaeum tikiterianum NZ13-K
Ca. Korarchaeum cryptofilum OPF8
Ca. Korarchaeum calidifontum LHC4
Euryarchaeota
0.99
0.98

**Fig. S10.** Phylogenetic tree corresponding to the Bayesian consensus tree (4 chains; CAT+GTR model) reconstructed from the untreated alignment based on 56 concatenated ribosomal proteins (RP56; 7158 alignment positions). *Ca*. Korarchaeota clades originating from hot springs and hydrothermal vents have a light and dark blue background shading, respectively. Scale bar indicates 0.1 substitutions per site.

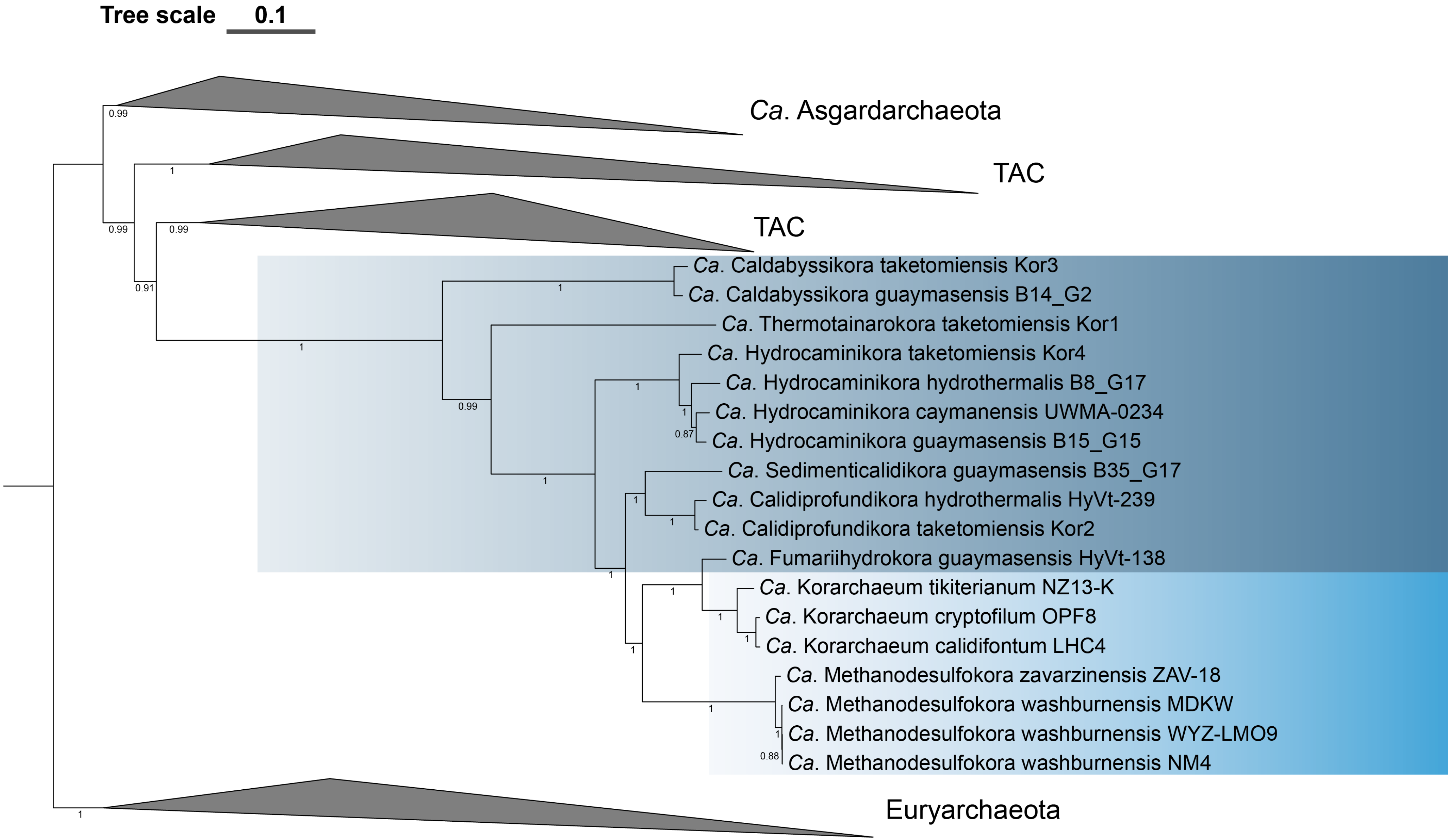
Tree scale 0.1
Ca. Asgardarchaeota
TAC
TAC
Ca. Caldabyssikora taketomiensis Kor3
Ca. Caldabyssikora guaymasensis B14_G2
Ca. Thermotainarokora taketomiensis Kor1
Ca. Hydrocaminikora taketomiensis Kor4
Ca. Hydrocaminikora hydrothermalis B8_G17
Ca. Hydrocaminikora caymanensis UWMA-0234
Ca. Hydrocaminikora guaymasensis B15_G15
Ca. Sedimenticalidikora guaymasensis B35_G17
Ca. Calidiprofundikora hydrothermalis HyVt-239
Ca. Calidiprofundikora taketomiensis Kor2
Ca. Fumariihydrokora guaymasensis HyVt-138
Ca. Korarchaeum tikiterianum NZ13-K
Ca. Korarchaeum cryptofilum OPF8
Ca. Korarchaeum calidifontum LHC4
Ca. Methanodesulfokora zavarzinensis ZAV-18
Ca. Methanodesulfokora washburnensis MDKW
Ca. Methanodesulfokora washburnensis WYZ-LMO9
Ca. Methanodesulfokora washburnensis NM4
Euryarchaeota

**Fig. S11.** Phylogenetic tree corresponding to the Bayesian consensus tree (4 chains; CAT+GTR model) reconstructed from the SR4-recoded alignment based on 56 concatenated ribosomal proteins (RP56; 7158 alignment positions). *Ca*. Korarchaeota clades originating from hot springs and hydrothermal vents have a light and dark blue background shading, respectively. Scale bar indicates 0.1 substitutions per site.

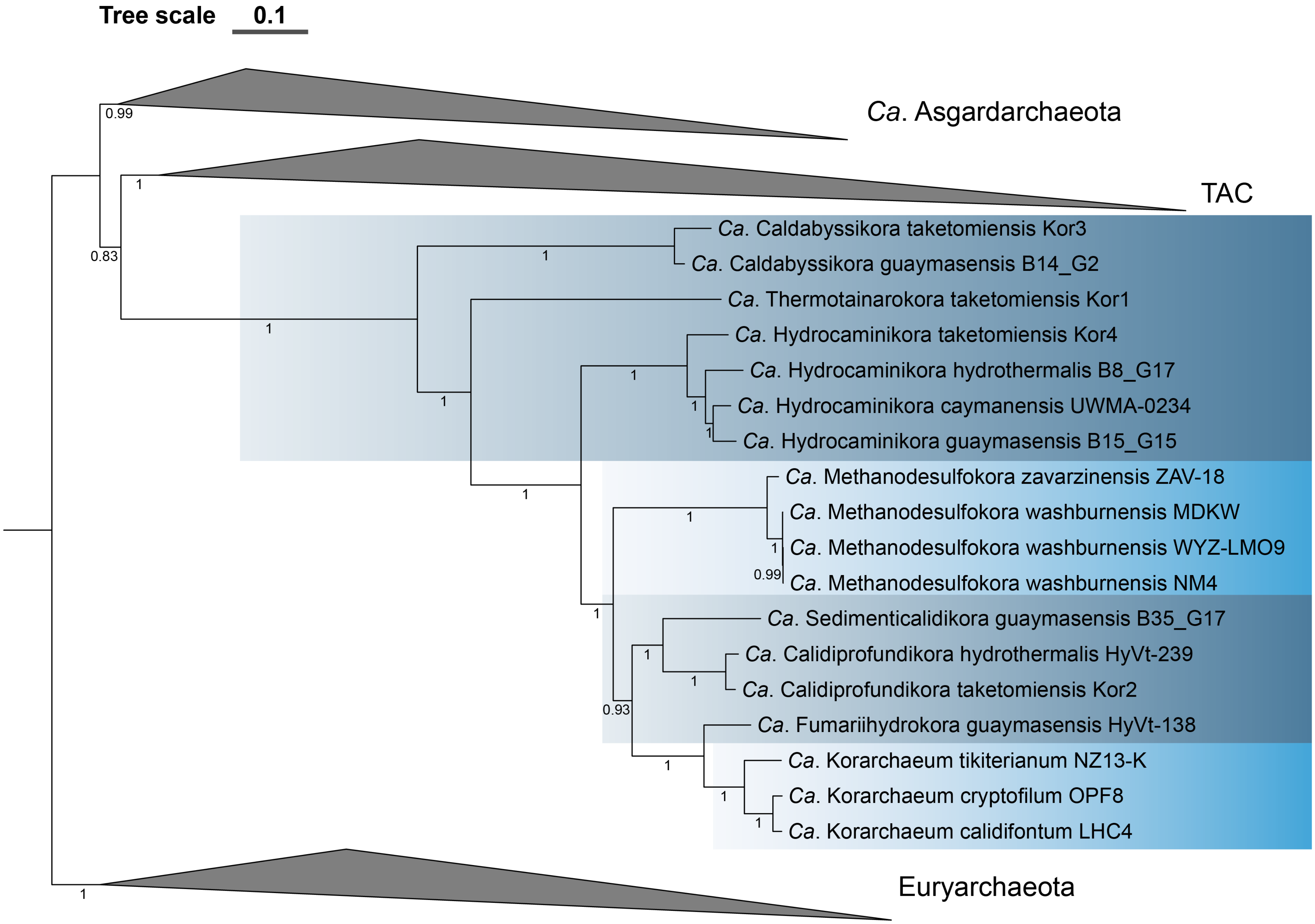
Tree scale 0.1
Ca. Asgardarchaeota
TAC
Ca. Caldabyssikora taketomiensis Kor3
Ca. Caldabyssikora guaymasensis B14_G2
Ca. Thermotainarokora taketomiensis Kor1
Ca. Hydrocaminikora taketomiensis Kor4
Ca. Hydrocaminikora hydrothermalis B8_G17
Ca. Hydrocaminikora caymanensis UWMA-0234
Ca. Hydrocaminikora guaymasensis B15_G15
Ca. Methanodesulfokora zavarzinensis ZAV-18
Ca. Methanodesulfokora washburnensis MDKW
Ca. Methanodesulfokora washburnensis WYZ-LMO9
Ca. Methanodesulfokora washburnensis NM4
Ca. Sedimenticalidikora guaymasensis B35_G17
Ca. Calidiprofundikora hydrothermalis HyVt-239
Ca. Calidiprofundikora taketomiensis Kor2
Ca. Fumariihydrokora guaymasensis HyVt-138
Ca. Korarchaeum tikiterianum NZ13-K
Ca. Korarchaeum cryptofilum OPF8
Ca. Korarchaeum calidifontum LHC4
Euryarchaeota
0.99
1
0.83
0.93

**Fig. S12.** Phylogenetic tree corresponding to the Bayesian consensus tree (4 chains; CAT+GTR model) reconstructed from the custom recoded alignment based on 56 concatenated ribosomal proteins (RP56; 7158 alignment positions). *Ca*. Korarchaeota clades originating from hot springs and hydrothermal vents have a light and dark blue background shading, respectively. Scale bar indicates 0.1 substitutions per site.

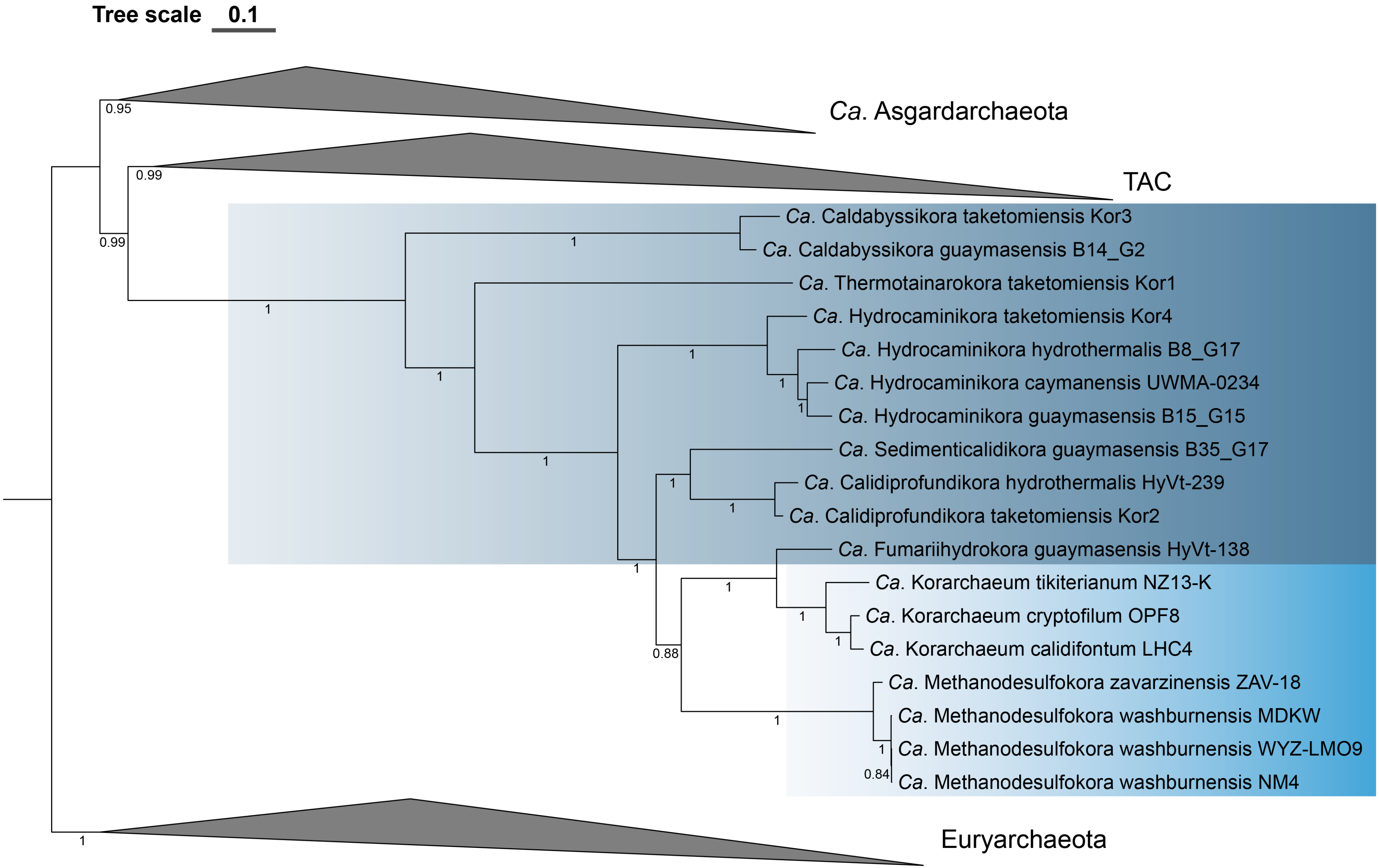
Tree scale 0.1
Ca. Asgardarchaeota
TAC
Ca. Caldabyssikora taketomiensis Kor3
Ca. Caldabyssikora guaymasensis B14_G2
Ca. Thermotainarokora taketomiensis Kor1
Ca. Hydrocaminikora taketomiensis Kor4
Ca. Hydrocaminikora hydrothermalis B8_G17
Ca. Hydrocaminikora caymanensis UWMA-0234
Ca. Hydrocaminikora guaymasensis B15_G15
Ca. Sedimenticalidikora guaymasensis B35_G17
Ca. Calidiprofundikora hydrothermalis HyVt-239
Ca. Calidiprofundikora taketomiensis Kor2
Ca. Fumariihydrokora guaymasensis HyVt-138
Ca. Korarchaeum tikiterianum NZ13-K
Ca. Korarchaeum cryptofilum OPF8
Ca. Korarchaeum calidifontum LHC4
Ca. Methanodesulfokora zavarzinensis ZAV-18
Ca. Methanodesulfokora washburnensis MDKW
Ca. Methanodesulfokora washburnensis WYZ-LMO9
Ca. Methanodesulfokora washburnensis NM4
Euryarchaeota
0.95
0.99
0.99
0.88
0.84
1

**Fig. S13.** Phylogenetic tree corresponding to the Bayesian consensus tree (4 chains; CAT+GTR model) reconstructed from the χ2-trimmed alignment (50% most heterogeneous sites removed, 3602 alignment positions) of 56 concatenated ribosomal proteins (RP56). *Ca*. Korarchaeota clades originating from hot springs and hydrothermal vents have a light and dark blue background shading, respectively. Scale bar indicates 0.1 substitutions per site.

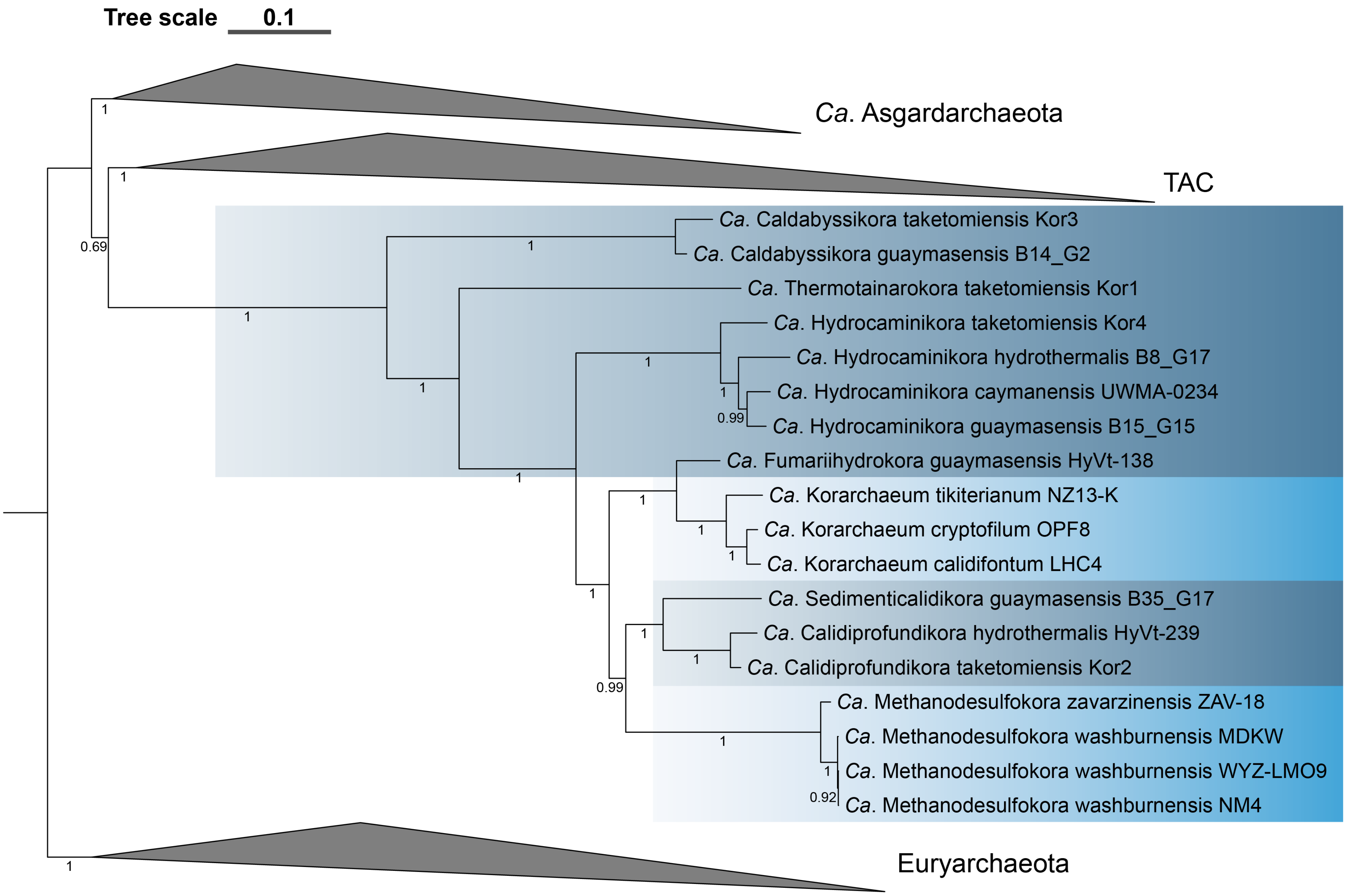
Tree scale 0.1
Ca. Asgardarchaeota
TAC
Ca. Caldabyssikora taketomiensis Kor3
Ca. Caldabyssikora guaymasensis B14_G2
Ca. Thermotainarokora taketomiensis Kor1
Ca. Hydrocaminikora taketomiensis Kor4
Ca. Hydrocaminikora hydrothermalis B8_G17
Ca. Hydrocaminikora caymanensis UWMA-0234
Ca. Hydrocaminikora guaymasensis B15_G15
Ca. Fumariihydrokora guaymasensis HyVt-138
Ca. Korarchaeum tikiterianum NZ13-K
Ca. Korarchaeum cryptofilum OPF8
Ca. Korarchaeum calidifontum LHC4
Ca. Sedimenticalidikora guaymasensis B35_G17
Ca. Calidiprofundikora hydrothermalis HyVt-239
Ca. Calidiprofundikora taketomiensis Kor2
Ca. Methanodesulfokora zavarzinensis ZAV-18
Ca. Methanodesulfokora washburnensis MDKW
Ca. Methanodesulfokora washburnensis WYZ-LMO9
Ca. Methanodesulfokora washburnensis NM4
Euryarchaeota
0.69
0.99
0.99
0.92

**Fig. S14.** Phylogenetic tree corresponding to the Bayesian consensus tree (4 chains; CAT+GTR model) reconstructed from the fast site trimmed alignment (50% fastest evolving sites removed, 3575 alignment positions) of 56 concatenated ribosomal proteins (RP56). *Ca*. Korarchaeota clades originating from hot springs and hydrothermal vents have a light and dark blue background shading, respectively. Scale bar indicates 0.1 substitutions per site.

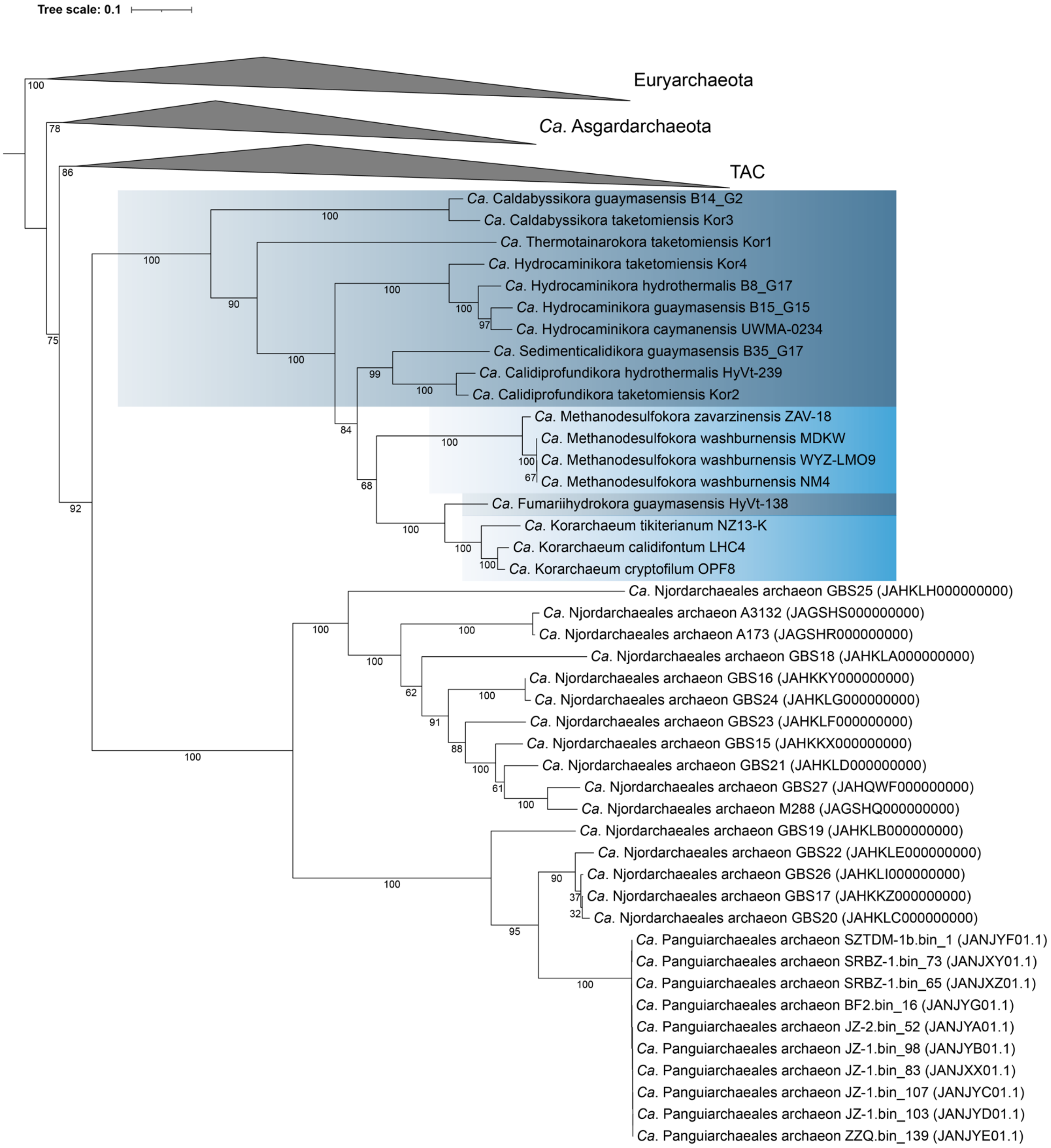


**Fig. S15.** Non-parametric bootstrap maximum-likelihood phylogenomic tree (IQ-TREE, 100 bootstraps, LG+C60+F+G4 model) based the χ2-trimmed alignment (50% most heterogeneous sites removed; 3328 alignment positions) of 56 concatenated ribosomal proteins (RP56) to which *Ca*. Njord- and *Ca*. Panguiarchaeales MAGs were added. Scale bar indicates 0.1 substitutions per site. *Candidatus* Korarchaeota originating from hot springs and hydrothermal vents have a light and dark blue background shading, respectively.
