## Supplementary Figures S16-S25 for "Phylogenomics and ancestral reconstruction of Korarchaeota reveals genomic adaptation to habitat switching"

*** Correspondence:**

Guillaume Tahon

Archaea, microbial diversity, phylogeny, hyperthermophiles, hot spring, genome evolution

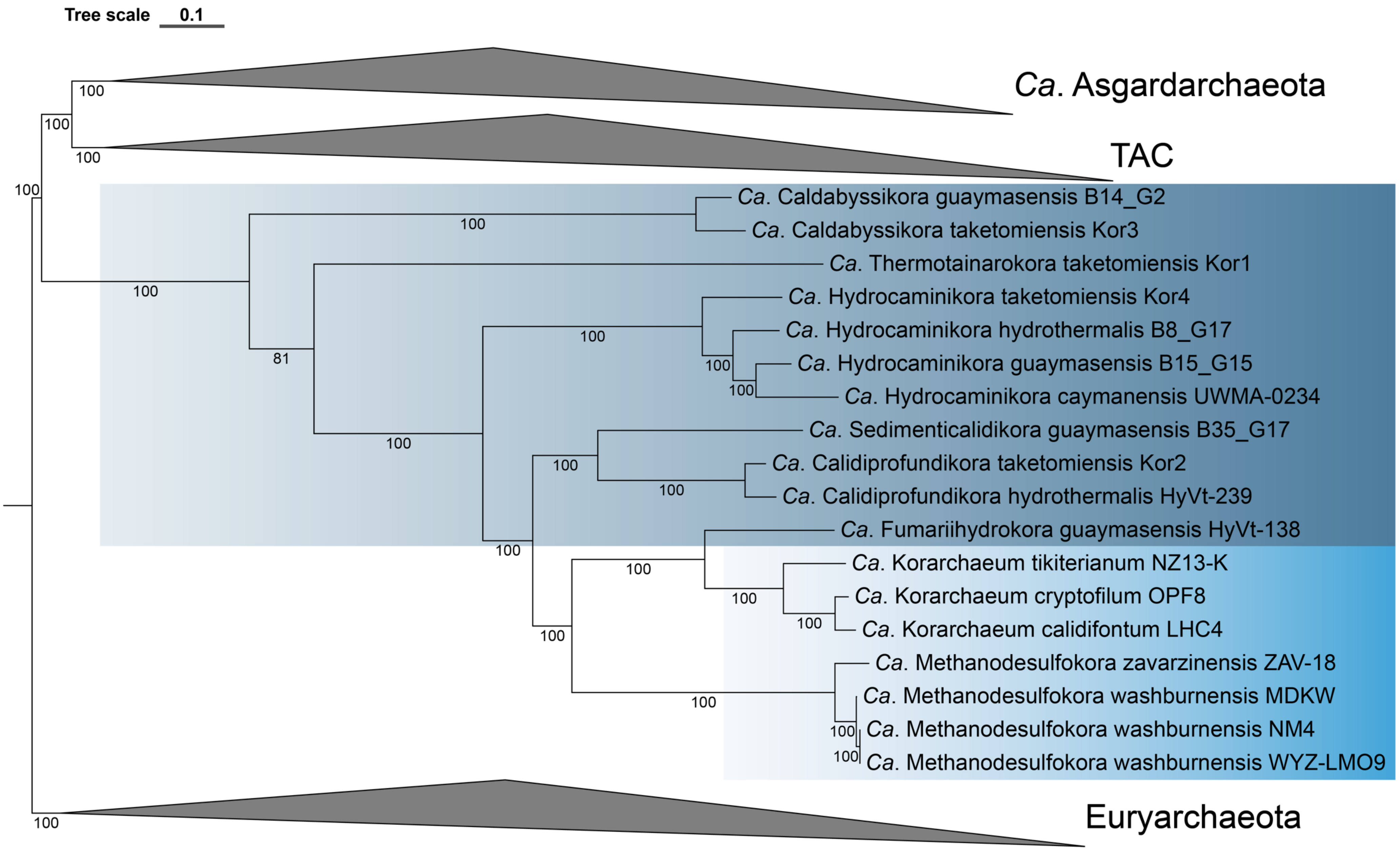

Tree scale 0.1
Ca. Asgardarchaeota
TAC
Ca. Caldabyssikora guaymasensis B14_G2
Ca. Caldabyssikora taketomiensis Kor3
Ca. Thermotainarokora taketomiensis Kor1
Ca. Hydrocaminikora taketomiensis Kor4
Ca. Hydrocaminikora hydrothermalis B8_G17
Ca. Hydrocaminikora guaymasensis B15_G15
Ca. Hydrocaminikora caymanensis UWMA-0234
Ca. Sedimenticalidikora guaymasensis B35_G17
Ca. Calidiprofundikora taketomiensis Kor2
Ca. Calidiprofundikora hydrothermalis HyVt-239
Ca. Fumariihydrokora guaymasensis HyVt-138
Ca. Korarchaeum tikiterianum NZ13-K
Ca. Korarchaeum cryptofilum OPF8
Ca. Korarchaeum calidifontum LHC4
Ca. Methanodesulfokora zavarzinensis ZAV-18
Ca. Methanodesulfokora washburnensis MDKW
Ca. Methanodesulfokora washburnensis NM4
Ca. Methanodesulfokora washburnensis WYZ-LMO9
Euryarchaeota

**Fig. S16.** Non-parametric bootstrap maximum-likelihood phylogenomic tree (IQ-TREE, 100 bootstraps, LG+C60+F+G4 model) based the untreated alignment of 54 concatenated new marker proteins (NM54; 15142 alignment positions). Scale bar indicates 0.1 substitutions per site. *Candidatus* Korarchaeota originating from hot springs and hydrothermal vents have a light and dark blue background shading, respectively.

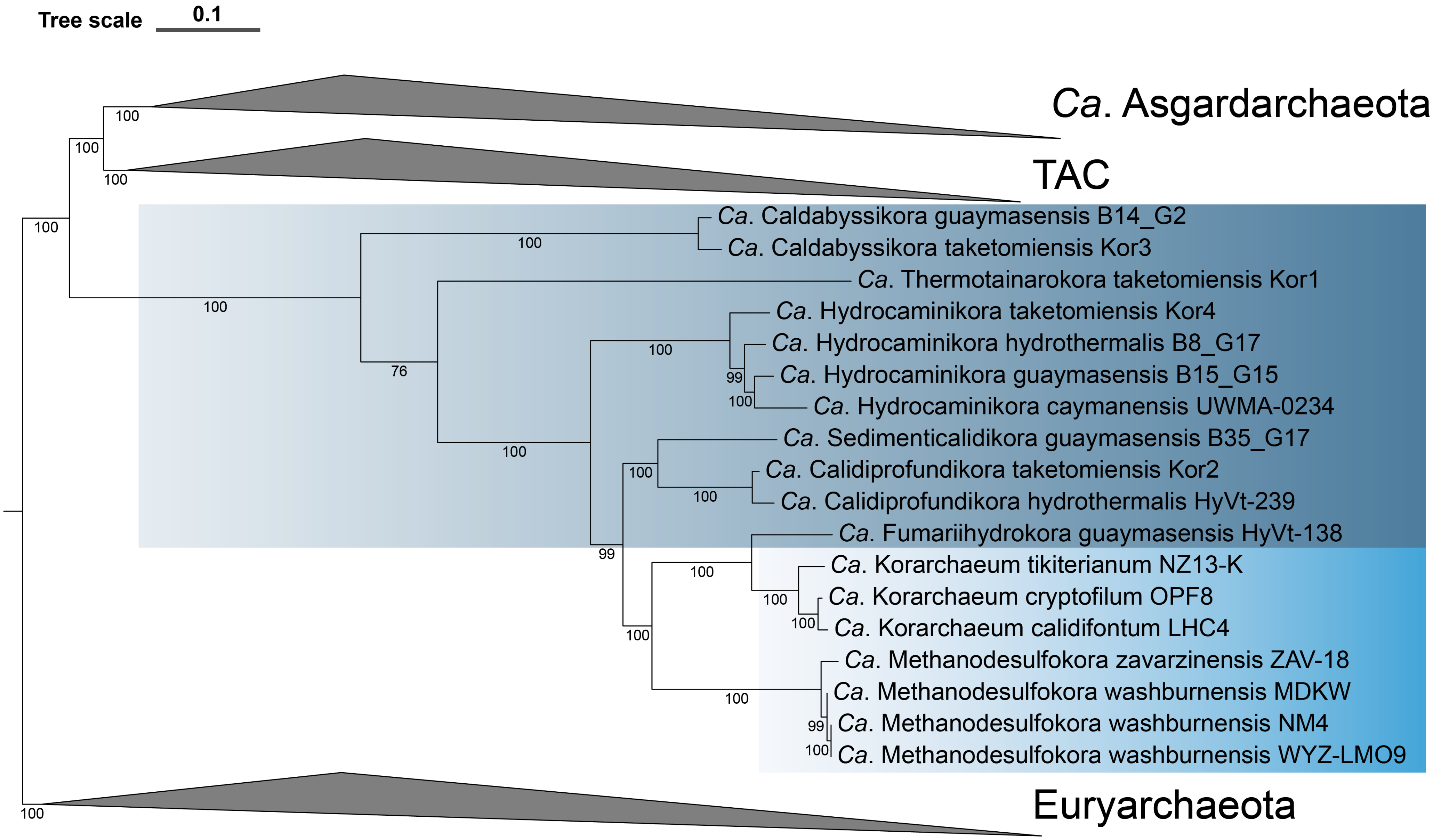
Tree scale 0.1
Ca. Asgardarchaeota
TAC
Ca. Caldabyssikora guaymasensis B14_G2
Ca. Caldabyssikora taketomiensis Kor3
Ca. Thermotainarokora taketomiensis Kor1
Ca. Hydrocaminikora taketomiensis Kor4
Ca. Hydrocaminikora hydrothermalis B8_G17
Ca. Hydrocaminikora guaymasensis B15_G15
Ca. Hydrocaminikora caymanensis UWMA-0234
Ca. Sedimenticalidikora guaymasensis B35_G17
Ca. Calidiprofundikora taketomiensis Kor2
Ca. Calidiprofundikora hydrothermalis HyVt-239
Ca. Fumariihydrokora guaymasensis HyVt-138
Ca. Korarchaeum tikiterianum NZ13-K
Ca. Korarchaeum cryptofilum OPF8
Ca. Korarchaeum calidifontum LHC4
Ca. Methanodesulfokora zavarzinensis ZAV-18
Ca. Methanodesulfokora washburnensis MDKW
Ca. Methanodesulfokora washburnensis NM4
Ca. Methanodesulfokora washburnensis WYZ-LMO9
Euryarchaeota

**Fig. S17.** Non-parametric bootstrap maximum-likelihood phylogenomic tree (IQ-TREE, 100 bootstraps, GTR+G4+C60SR4 model) based on an SR4-recoded concatenation of 54 new marker proteins (NM54; 15142 alignment positions). Scale bar indicates 0.1 substitutions per site. *Candidatus* Korarchaeota originating from hot springs and hydrothermal vents have a light and dark blue background shading, respectively.

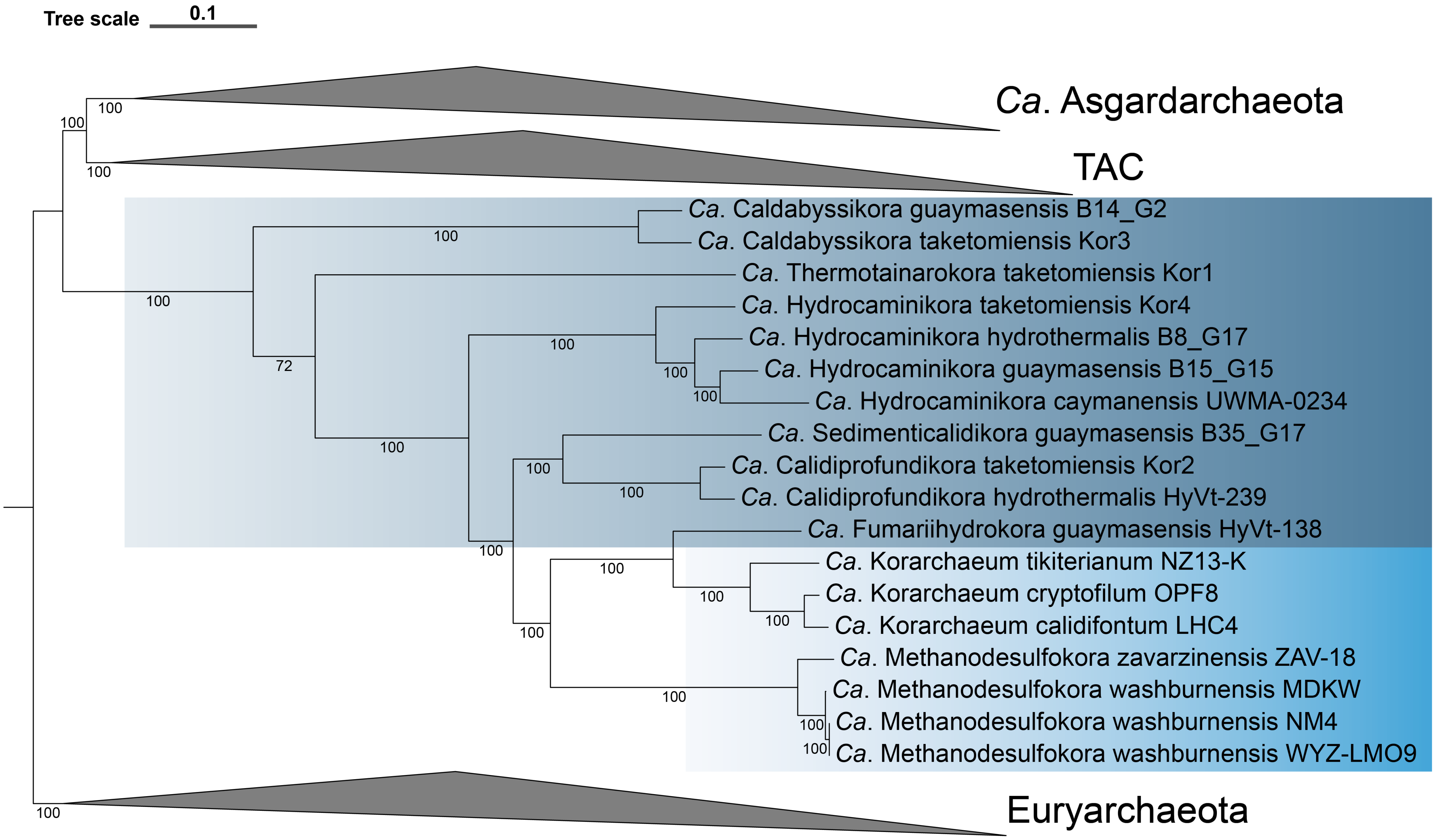
Tree scale
0.1
Ca. Asgardarchaeota
TAC
Ca. Caldabyssikora guaymasensis B14_G2
Ca. Caldabyssikora taketomiensis Kor3
Ca. Thermotainarokora taketomiensis Kor1
Ca. Hydrocaminikora taketomiensis Kor4
Ca. Hydrocaminikora hydrothermalis B8_G17
Ca. Hydrocaminikora guaymasensis B15_G15
Ca. Hydrocaminikora caymanensis UWMA-0234
Ca. Sedimenticalidikora guaymasensis B35_G17
Ca. Calidiprofundikora taketomiensis Kor2
Ca. Calidiprofundikora hydrothermalis HyVt-239
Ca. Fumariihydrokora guaymasensis HyVt-138
Ca. Korarchaeum tikiterianum NZ13-K
Ca. Korarchaeum cryptofilum OPF8
Ca. Korarchaeum calidifontum LHC4
Ca. Methanodesulfokora zavarzinensis ZAV-18
Ca. Methanodesulfokora washburnensis MDKW
Ca. Methanodesulfokora washburnensis NM4
Ca. Methanodesulfokora washburnensis WYZ-LMO9
Euryarchaeota
100
72

**Fig. S18.** Non-parametric bootstrap maximum-likelihood phylogenomic tree (IQ-TREE, 100 bootstraps, LG+C60+F+G4 model) based on a custom recoded concatenation of 54 new marker proteins (NM54; 15142 alignment positions). Scale bar indicates 0.1 substitutions per site. *Candidatus* Korarchaeota originating from hot springs and hydrothermal vents have a light and dark blue background shading, respectively.

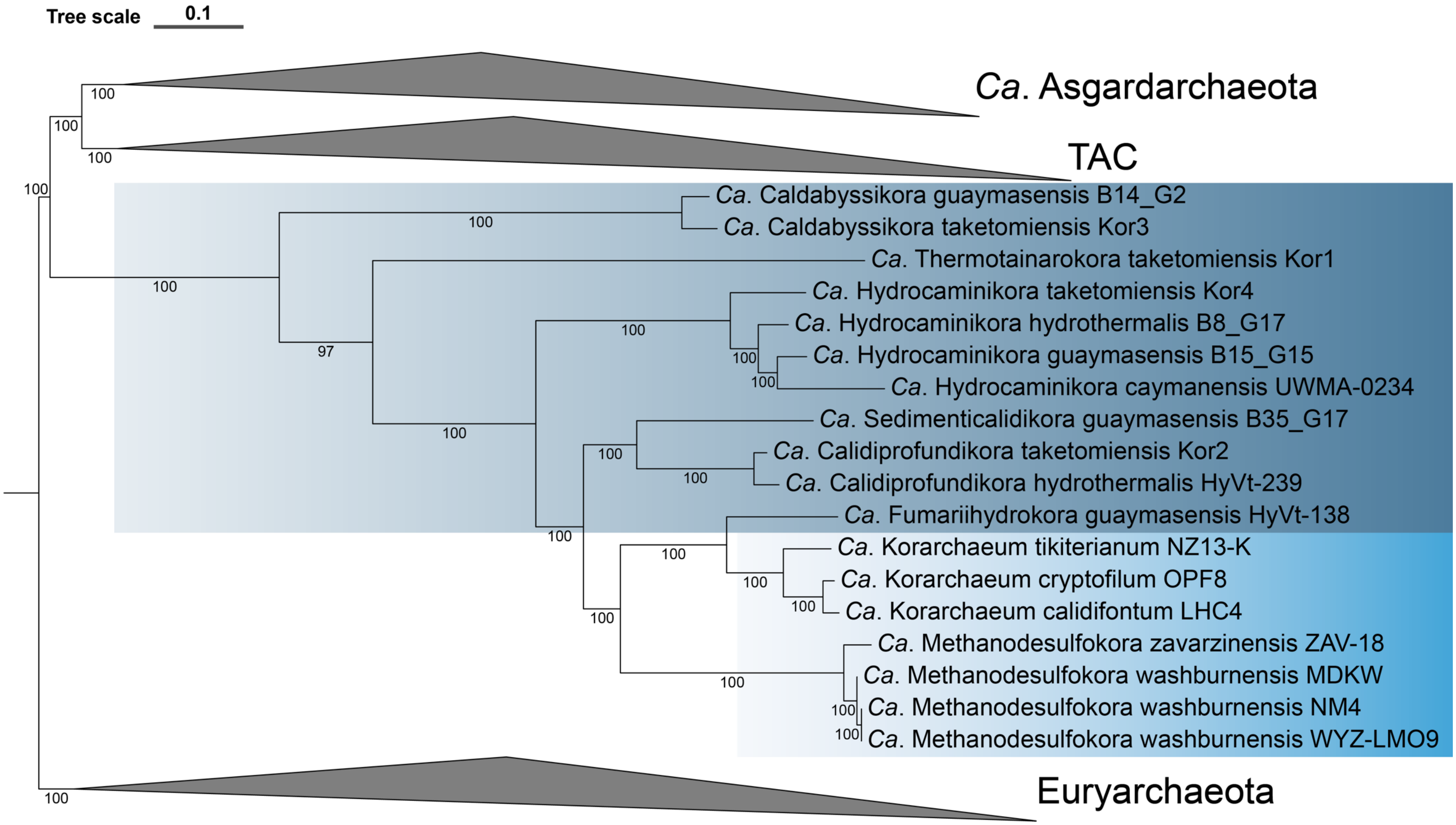


**Fig. S19.** Non-parametric bootstrap maximum-likelihood phylogenomic tree (IQ-TREE, 100 bootstraps, LG+C60+F+G4 model) based on a χ2-trimmed alignment (50% most heterogeneous sites removed, 7594 alignment positions) of 54 concatenated new marker proteins (NM54). Scale bar indicates 0.1 substitutions per site. *Candidatus* Korarchaeota originating from hot springs and hydrothermal vents have a light and dark blue background shading, respectively.

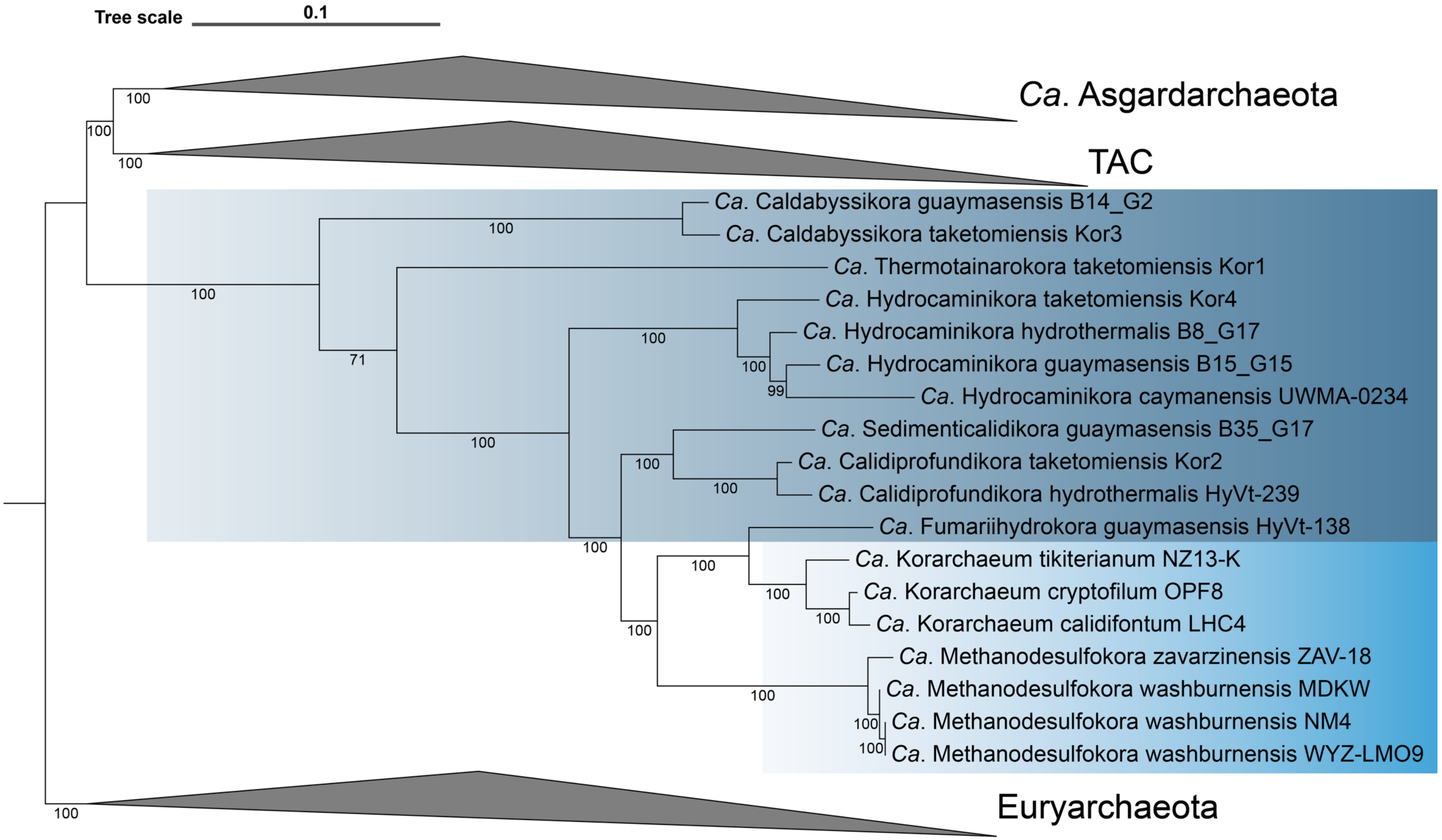


**Fig. S20.** Non-parametric bootstrap maximum-likelihood phylogenomic tree (IQ-TREE, 100 bootstraps, LG+C60+F+G4 model) based on a fast site trimmed alignment (50% fastest evolving sites removed, 7571 alignment positions) of 54 concatenated new marker proteins (NM54). Scale bar indicates 0.1 substitutions per site. *Candidatus* Korarchaeota originating from hot springs and hydrothermal vents have a light and dark blue background shading, respectively.

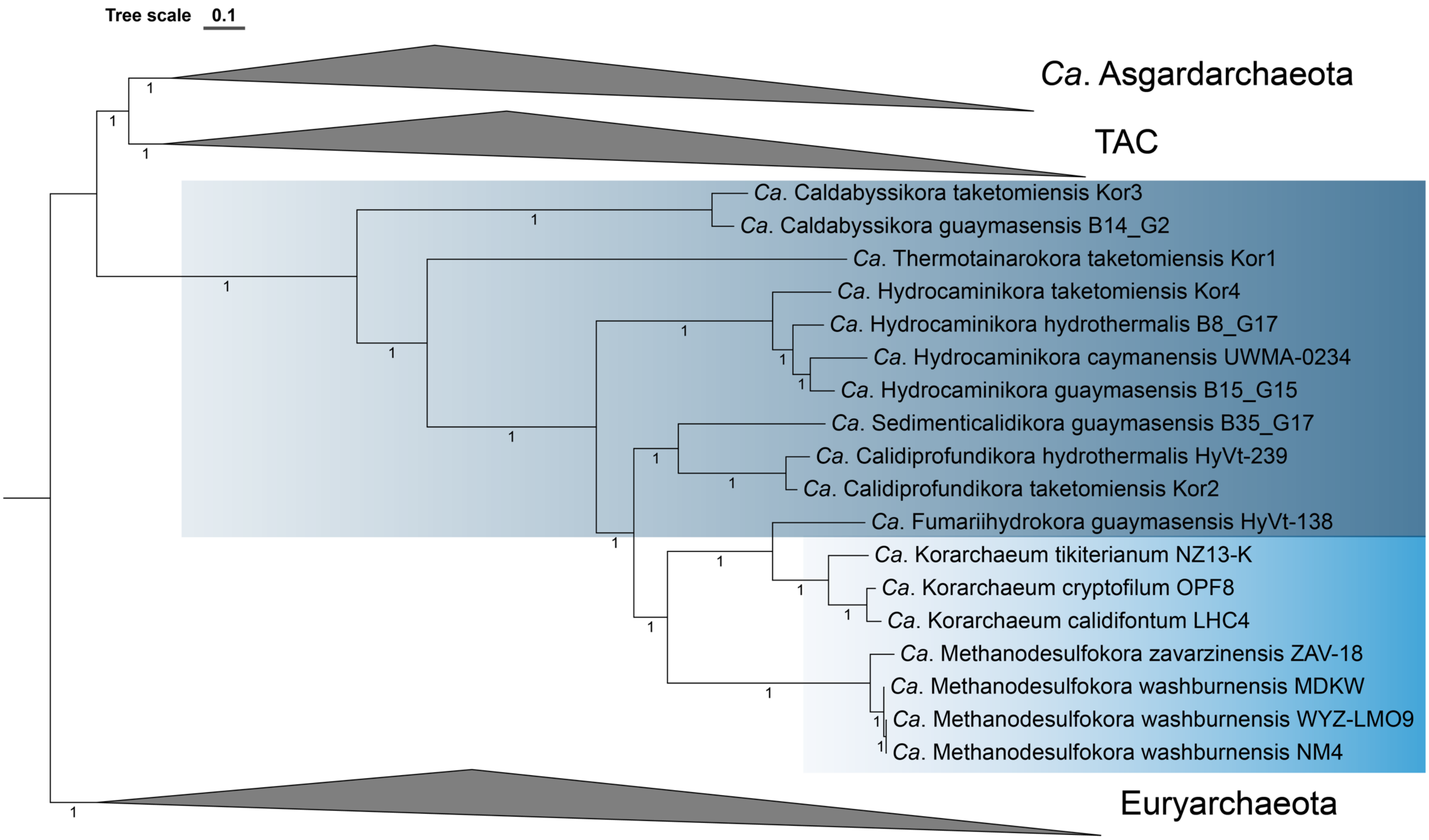
Tree scale 0.1
Ca. Asgardarchaeota
TAC
Ca. Caldabyssikora taketomiensis Kor3
Ca. Caldabyssikora guaymasensis B14_G2
Ca. Thermotainarokora taketomiensis Kor1
Ca. Hydrocaminikora taketomiensis Kor4
Ca. Hydrocaminikora hydrothermalis B8_G17
Ca. Hydrocaminikora caymanensis UWMA-0234
Ca. Hydrocaminikora guaymasensis B15_G15
Ca. Sedimenticalidikora guaymasensis B35_G17
Ca. Calidiprofundikora hydrothermalis HyVt-239
Ca. Calidiprofundikora taketomiensis Kor2
Ca. Fumariihydrokora guaymasensis HyVt-138
Ca. Korarchaeum tikiterianum NZ13-K
Ca. Korarchaeum cryptofilum OPF8
Ca. Korarchaeum calidifontum LHC4
Ca. Methanodesulfokora zavarzinensis ZAV-18
Ca. Methanodesulfokora washburnensis MDKW
Ca. Methanodesulfokora washburnensis WYZ-LMO9
Ca. Methanodesulfokora washburnensis NM4
Euryarchaeota

**Fig. S21.** Phylogenetic tree corresponding to the Bayesian consensus tree (4 chains; CAT+GTR model) reconstructed from the untreated alignment based on 54 concatenated new marker proteins (NM54; 15142 alignment positions). *Ca*. Korarchaeota clades originating from hot springs and hydrothermal vents have a light and dark blue background shading, respectively. Scale bar indicates 0.1 substitutions per site.

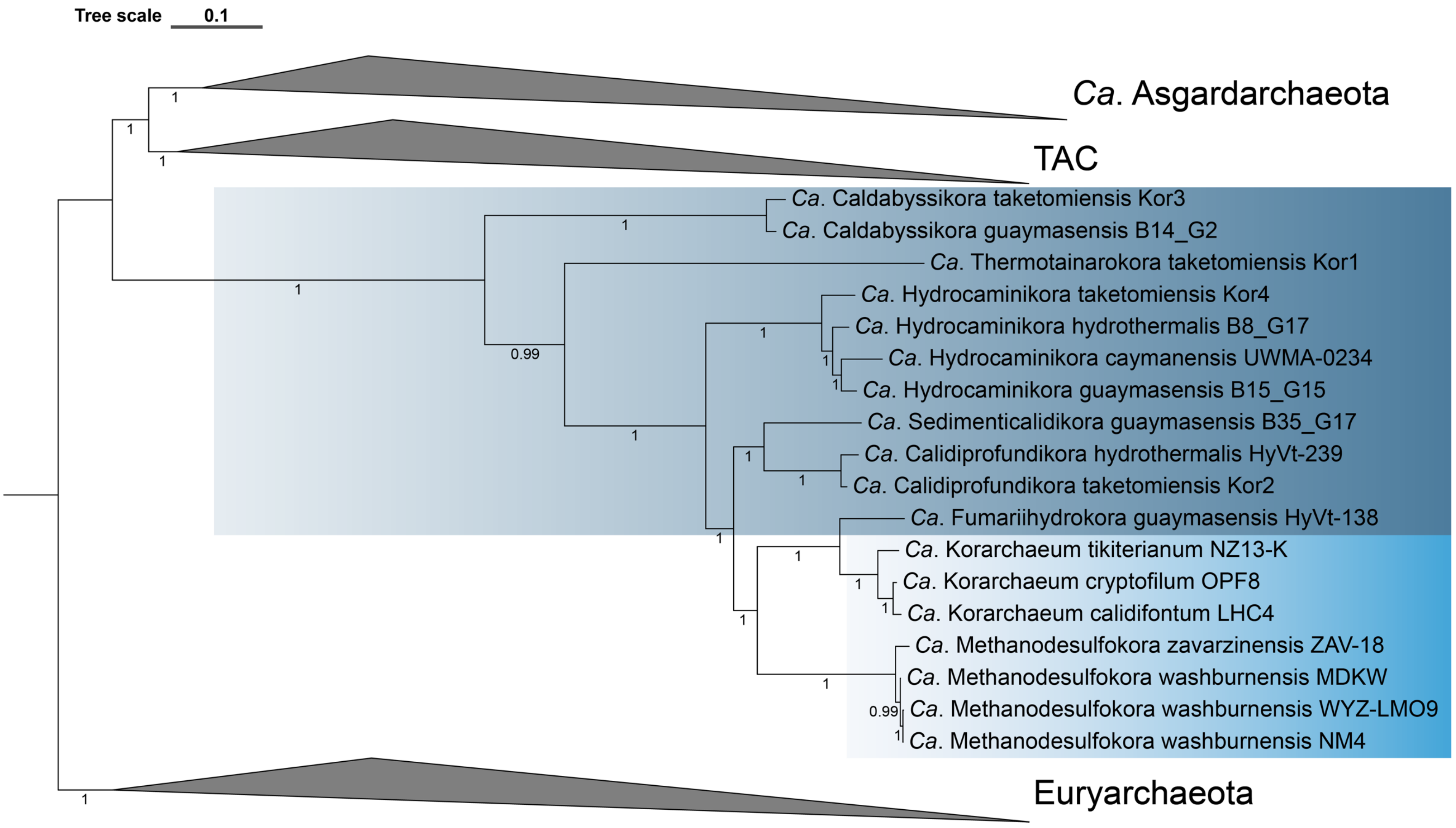


**Fig. S22.** Phylogenetic tree corresponding to the Bayesian consensus tree (4 chains; CAT+GTR model) reconstructed from the SR4-recoded alignment based on 54 concatenated new marker proteins (NM54; 15142 alignment positions). *Ca*. Korarchaeota clades originating from hot springs and hydrothermal vents have a light and dark blue background shading, respectively. Scale bar indicates 0.1 substitutions per site.

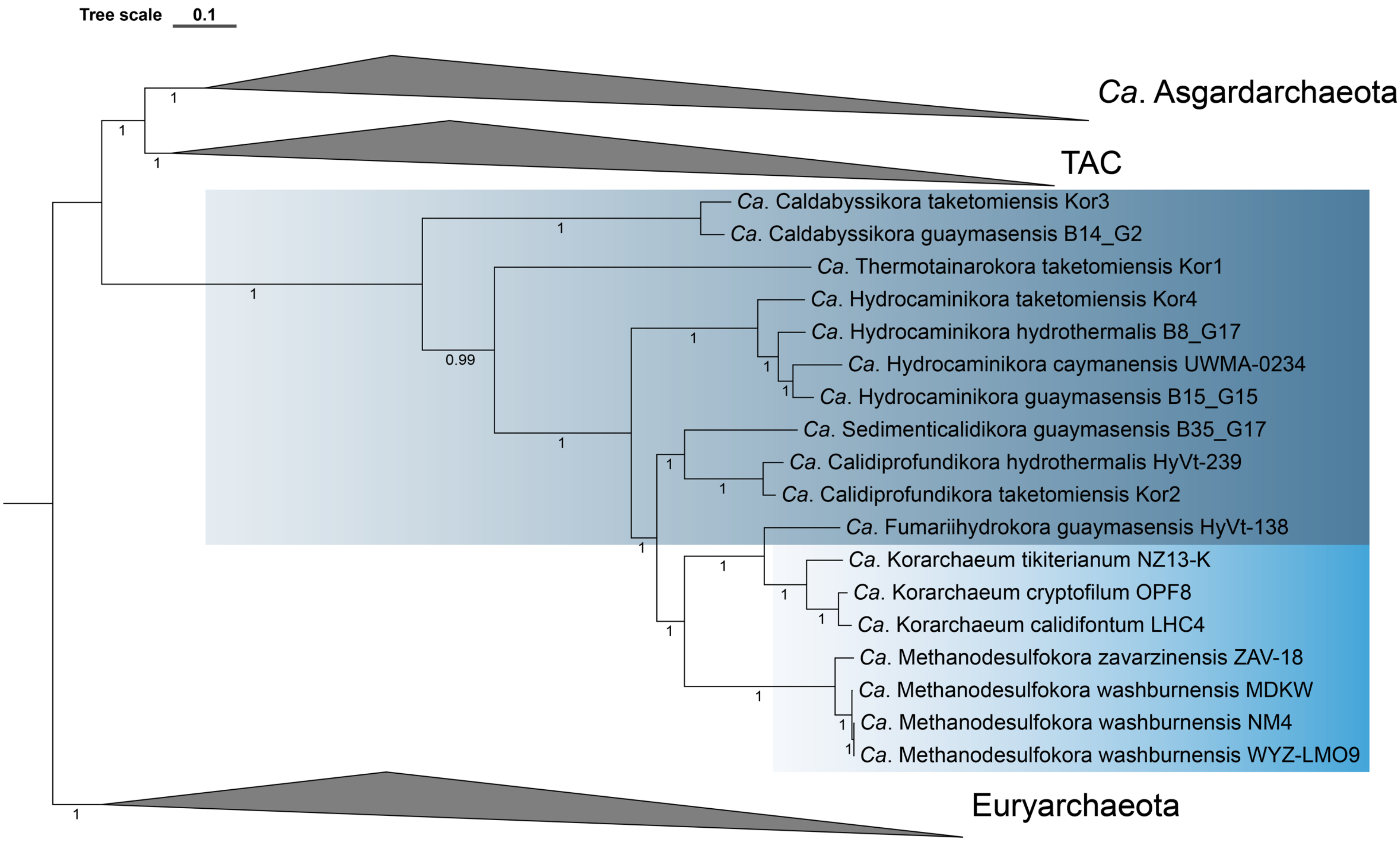
Tree scale 0.1
Ca. Asgardarchaeota
TAC
Ca. Caldabyssikora taketomiensis Kor3
Ca. Caldabyssikora guaymasensis B14_G2
Ca. Thermotainarokora taketomiensis Kor1
Ca. Hydrocaminikora taketomiensis Kor4
Ca. Hydrocaminikora hydrothermalis B8_G17
Ca. Hydrocaminikora caymanensis UWMA-0234
Ca. Hydrocaminikora guaymasensis B15_G15
Ca. Sedimenticalidikora guaymasensis B35_G17
Ca. Calidiprofundikora hydrothermalis HyVt-239
Ca. Calidiprofundikora taketomiensis Kor2
Ca. Fumariihydrokora guaymasensis HyVt-138
Ca. Korarchaeum tikiterianum NZ13-K
Ca. Korarchaeum cryptofilum OPF8
Ca. Korarchaeum calidifontum LHC4
Ca. Methanodesulfokora zavarzinensis ZAV-18
Ca. Methanodesulfokora washburnensis MDKW
Ca. Methanodesulfokora washburnensis NM4
Ca. Methanodesulfokora washburnensis WYZ-LMO9
Euryarchaeota
0.99

**Fig. S23.** Phylogenetic tree corresponding to the Bayesian consensus tree (4 chains; CAT+GTR model) reconstructed from the custom recoded alignment based on 54 concatenated new marker proteins (NM54; 15142 alignment positions). *Ca*. Korarchaeota clades originating from hot springs and hydrothermal vents have a light and dark blue background shading, respectively. Scale bar indicates 0.1 substitutions per site.

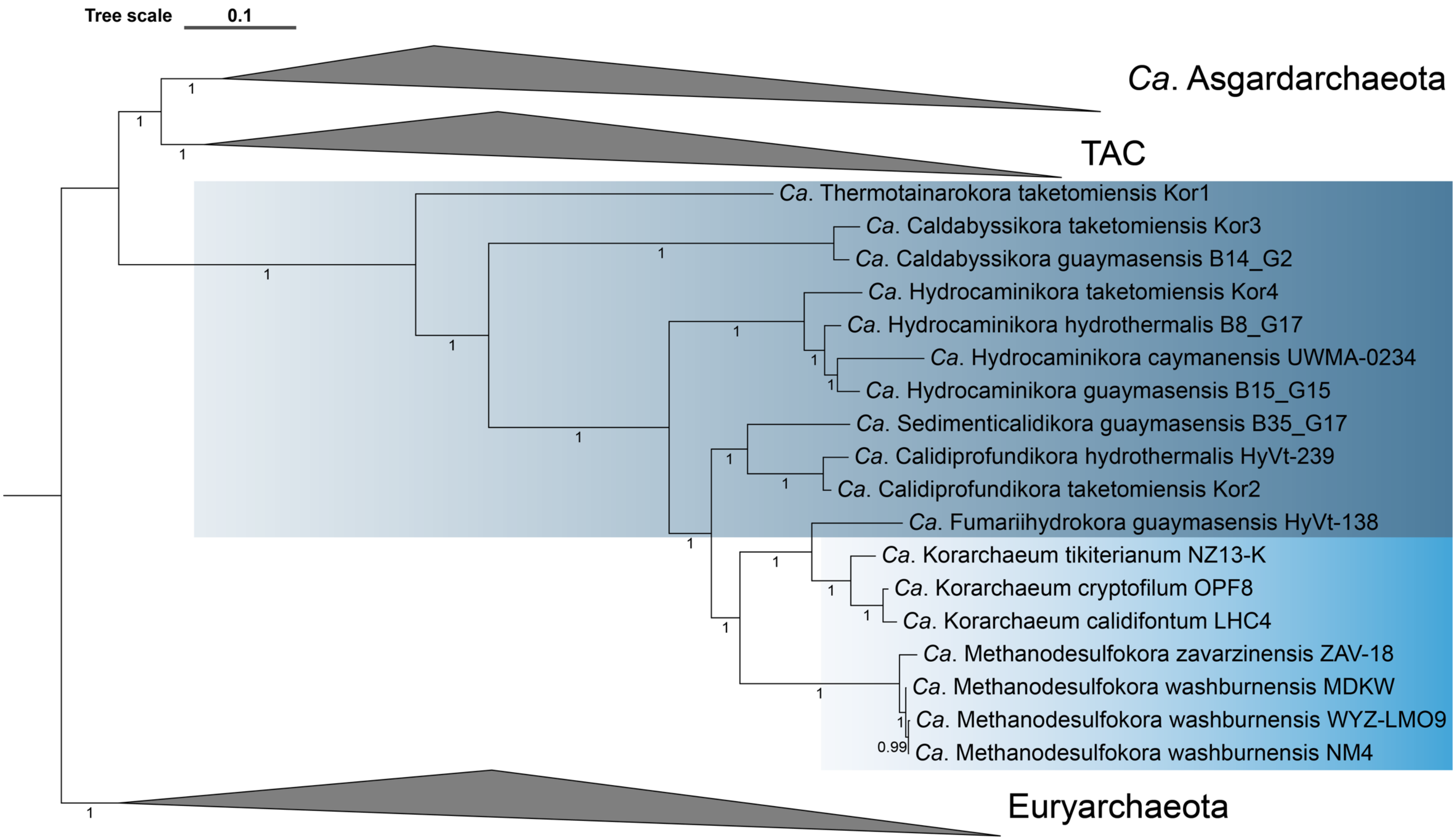


**Fig. S24.** Phylogenetic tree corresponding to the Bayesian consensus tree (4 chains; CAT+GTR model) reconstructed from the fast site trimmed alignment (50% fastest evolving sites removed, 7571 alignment positions) of 54 concatenated new marker proteins (NM54). *Ca.* Korarchaeota clades originating from hot springs and hydrothermal vents have a light and dark blue background shading, respectively. Scale bar indicates 0.1 substitutions per site.

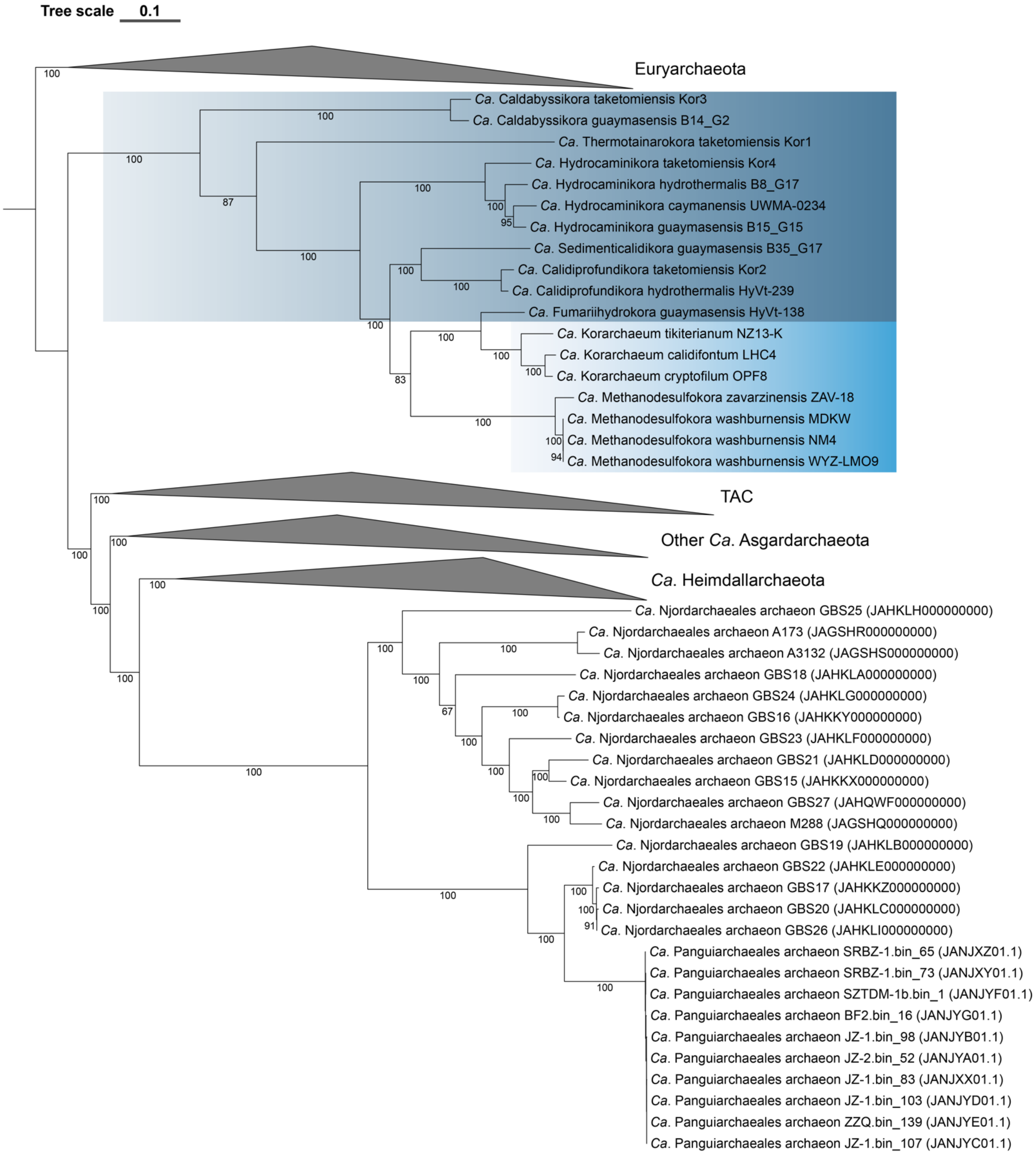


**Fig. S25.** Non-parametric bootstrap maximum-likelihood phylogenomic tree (IQ-TREE, 100 bootstraps, LG+C60+F+G4 model) based the χ2-trimmed alignment (50% most heterogeneous sites removed; 7045 alignment positions) of 54 concatenated new marker proteins (NM54) to which *Ca*. Njord- and *Ca*. Panguiarchaeales MAGs were added. Scale bar indicates 0.1 substitutions per site. *Candidatus* Korarchaeota originating from hot springs and hydrothermal vents have a light and dark blue background shading, respectively.
