## Supplementary Figures S26-S27 for "Phylogenomics and ancestral reconstruction of Korarchaeota reveals genomic adaptation to habitat switching"

*** Correspondence:**

Guillaume Tahon

Archaea, microbial diversity, phylogeny, hyperthermophiles, hot spring, genome evolution

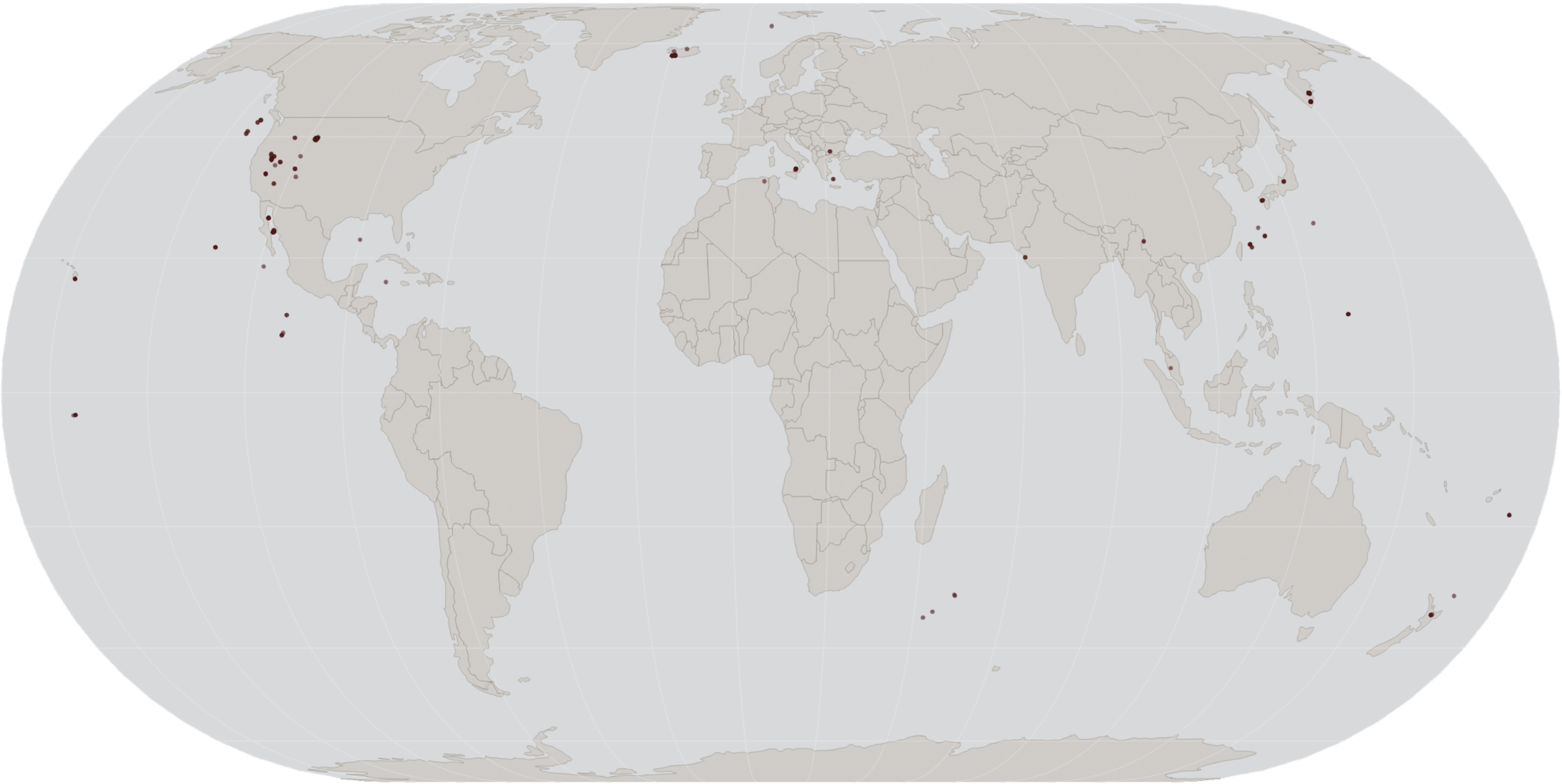

**Fig. S26.** Geographic distribution of *Ca*. Korarchaeota based on sampling locations where MAGs and 16S rRNA genes of the phylum were found (Table S2-S4).

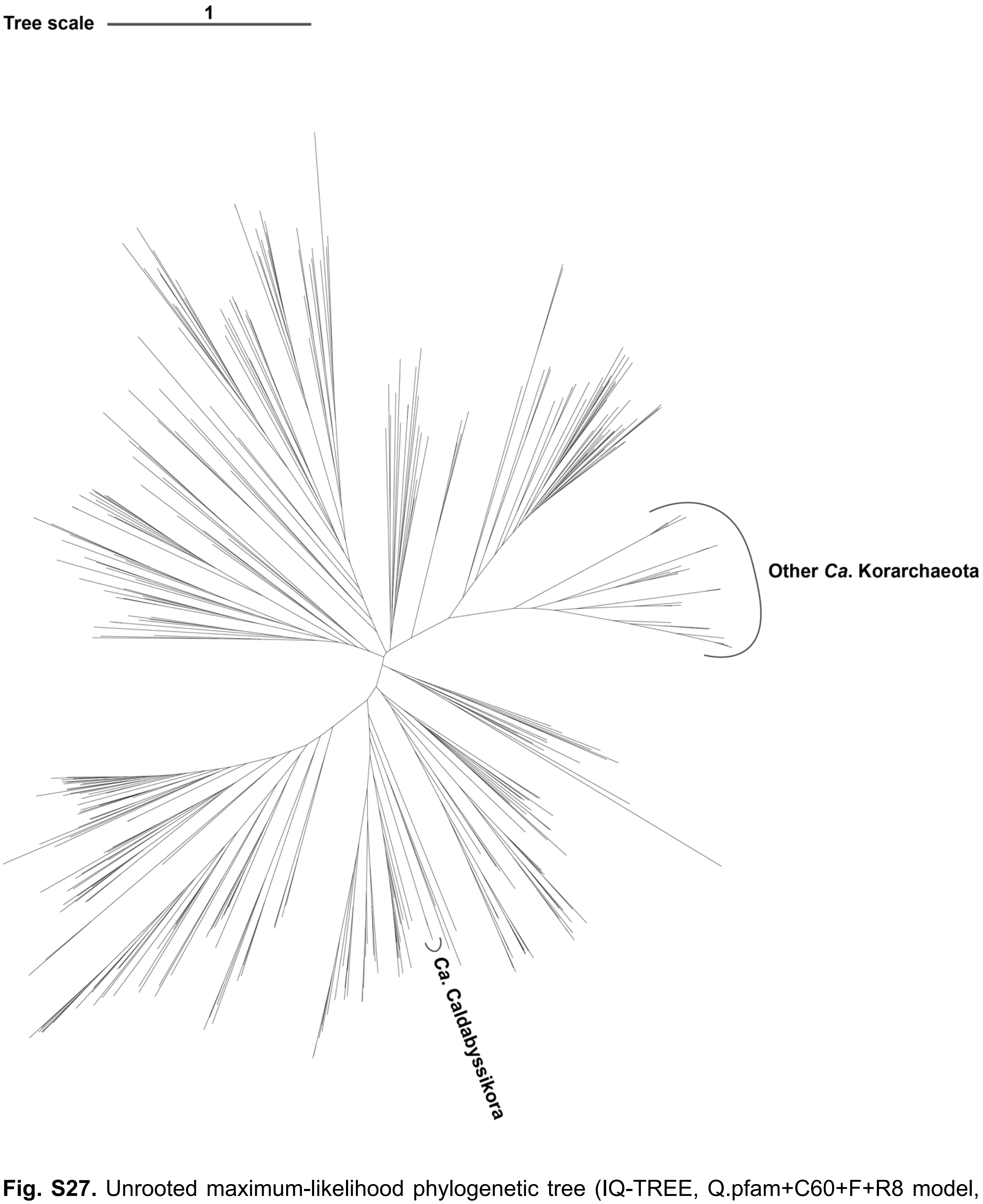


**Fig. S27.** Unrooted maximum-likelihood phylogenetic tree (IQ-TREE, Q.pfam+C60+F+R8 model, 1000 ultrafast bootstrap replicates, 1000 approximate likelihood-ratio test, 1245 alignment positions) of reverse gyrase sequences retrieved from UniProt and the 137 MAGs included in this study (Table S1-S2).
